# Estrus-Tracking Cortical Neurons Integrate Social cues and Reproductive states to Adaptively Control Sexually Dimorphic Sociosexual Behaviors

**DOI:** 10.1101/2024.08.30.610466

**Authors:** Yuping Wang, Xinli Song, Xiangmao Chen, Ying Zhou, Jihao Ma, Fang Zhang, Liqiang Wei, Guoxu Qi, Nakul Yadav, Benjie Miao, Yiming Yan, Guohua Yuan, Da Mi, Priyamvada Rajasethupathy, Ines Ibañez-Tallon, Xiaoxuan Jia, Nathaniel Heintz, Kun Li

## Abstract

Female sociosexual behaviors, essential for survival and reproduction, are adaptively modulated by ovarian hormones. However, the neural mechanisms integrating internal hormonal states with external social cues to guide these behaviors remain poorly understood. Here we identified primary estrous-sensitive *Cacna1h*-expressing medial prefrontal (mPFC^Cacna1h+^) neurons that orchestrate adaptive sociosexual behaviors. Bidirectional manipulation of mPFC^Cacna1h+^ neurons drives opposite-sex-directed behavioral shifts between estrus and diestrus females. In males, these neurons serve opposite functions compared to estrus females, mediating sexually dimorphic effects via anterior hypothalamic outputs. Miniscope imaging reveals mixed-representation of self-estrous states and social target sex in distinct mPFC^Cacna1h+^ subpopulations, with biased-encoding of opposite-sex social cues in estrus females and males. Mechanistically, ovarian hormone-driven upregulation of *Cacna1h*-encoded T-type calcium channels underlies estrus-specific activity changes and sexual-dimorphic function of mPFC^Cacna1h+^ neurons. These findings uncover a prefrontal circuit that integrates internal hormonal states and target-sex information to exert sexually bivalent top-down control over adaptive social behaviors.

## INTRODUCTION

Ovarian hormone fluctuations intricately orchestrate female internal physiological states^1,2^, and distinctly influence sex-specific social and emotional responses^3–5^, significantly contributing to the observed sex disparities in psychiatric disorder susceptibilities^6–8^. Across species, female social preference and sexual receptivity towards males are synchronized with the estrous cycle, specifically ovulation, optimizing reproductive success^9^. This suggests that female brains integrate social cues related to sex with their reproductive states to guide adaptive sociosexual interactions, such as social approach, investigation, avoidance, or sexual receptivity toward males^10–13^. However, the neural mechanisms underlying this integration in sociosexual behavior remain largely unknown.

Female adaptive sociosexual behaviors toward males require flexible adjustment across estrous states, aligning with the role of the medial prefrontal cortex (mPFC) in integrating diverse information^14^ and modulating behavioral flexibility^15–17^. The mPFC, a critical node in the social brain network, governs various aspects of social behavior, including social recognition^18^, investigation^19^, representation of conspecific sex^20^, and perception of social status^21,22^. The mPFC receives substantial inputs from olfactory pathways and projects to discrete subcortical areas mediating social reward, memory, aggression, and avoidance^23^. The evidence above suggests that the mPFC may exert a top-down role in the social brain network by integrating internal and external social contexts and controlling hardwired subcortical circuits to guide adaptive social behaviors.

Neuroimaging studies in humans have linked mPFC activity fluctuations to cognitive stimuli throughout the menstrual cycle^24–27^. Functional MRI studies in female rats have shown that the estrous cycle significantly alters neural activity across the brain^25–28^. Recent work identified increased activity of Esr1+/Cckar+ neurons in the ventrolateral subdivision of the ventromedial hypothalamus (VMHvl) as a key network node to control female sexual behaviors during estrus^4,29,30^. Despite low estrogen receptor expression, the estrous cycle in rodents has been shown to modulate stress-triggered immediate early gene (IEG) expression, dendritic remodeling, synaptic plasticity and behavioral responses to stress in the mPFC^3,31–33^. Specifically, conditional deletion of the Oxtr gene or Oxtr+/Sst+ interneurons (OxtrINs) in the mPFC attenuates females’ social preference for males during estrus, emphasizing the mPFC’s significant role in estrus-specific sociosexual preference^3^. Yet, how mPFC neurons encode self-estrous states, and integrate external social cue with self-estrous states to adaptively regulate male-directed sociosexual behaviors in females remain unknown.

To address this critical knowledge gap, we employed comprehensive unbiased screening techniques to identify a subset of *Cacna1h*-expressing layer 5 pyramidal tract neurons in the mPFC (mPFC^Cacna1h+^ neurons) that exhibit high sensitivity to estrous cycle changes in gene transcription and neuronal activity. Through bidirectional neural modulation, in vivo calcium imaging, computational analysis and mechanistic investigations, we demonstrate that mPFC^Cacna1h+^ neurons integrate social cues with self-estrous states to orchestrate increased male-directed sociosexual behaviors in estrus females, while conversely suppressing female-directed behaviors in males. We further provide cellular and molecular mechanistic insights into the estrus-dependent and sex-specific cortical mechanism underlying the intricate interplay between physiological processes and social adaptation. These findings provide direct evidence for the mPFC’s role in integrating internal and external social contexts to guide adaptive sociosexual behaviors, offering novel insights into the top-down regulation of adaptive innate social behaviors. The sexually bivalent roles of estrous-sensitive mPFC^Cacna1h+^ neurons and *Cacna1h* gene suggest that prefrontal dysfunction may underlie the high prevalence of hypersexuality in males and hyposexuality in females, elucidating the neural basis of sexual dimorphism and highlighting potential therapeutic targets for sexual disorders.

## RESULTS

### Estrous cycle shapes mPFC neural type transcriptomes

Single-cell RNA sequencing of adult mPFC samples from estrus females, diestrus females, and males was performed to characterize the molecular features of mPFC neurons responsive to estrous cycle changes (Figure 1A). t-SNE analysis identified six excitatory and four inhibitory neuron clusters (Figure 1B), annotated based on established gene markers (Figure 1C) ^34–37^. Sex hormone receptor expression was low for estrogen receptor and distributed for progesterone and androgen receptors across all mPFC neural types (Figure 1D). Differential gene expression (DEG) analysis revealed that Pvalb+ and Lamp5+ interneurons, and layer 5 pyramidal tract (L5_PT) and layer 5/6 near-projecting (L5/6_NP) excitatory neurons, exhibited the highest number of DEGs when comparing estrus to diestrus (Figure S1F), suggesting heightened susceptibility to the diestrus-to-estrus transition.

**Figure 1.**
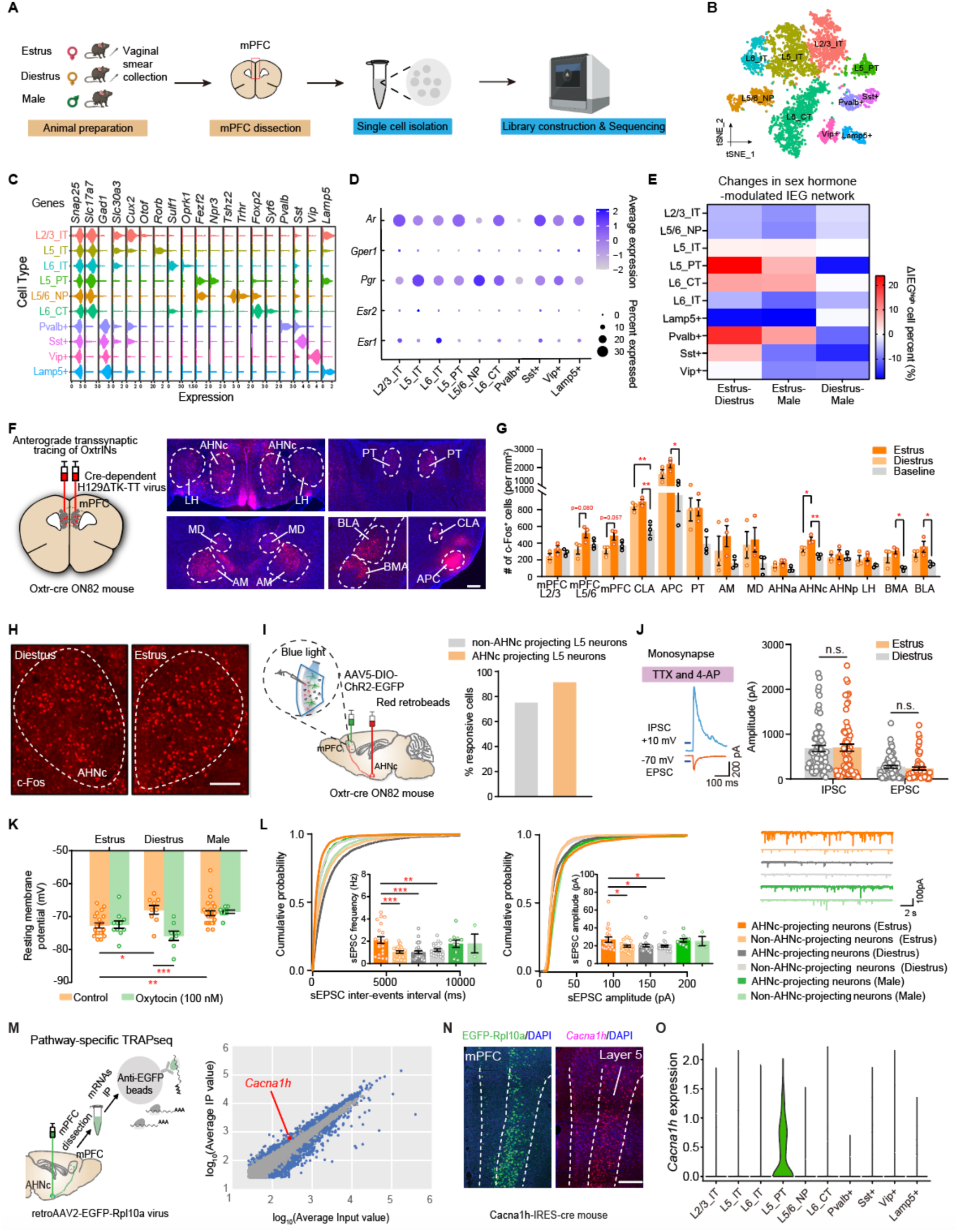
Comprehensive unbiased screenings identify *Cacna1h*-defined L5_PT neurons highly sensitive to estrus cycle oscillation. (A) Single cell RNA-seq strategy of the mPFC in estrus, diestrus female and male mice. (B) t-SNE space visualization of defined mPFC neuron clusters, colored by cell types (IT, intratelencephalically projecting; PT, pyramidal tract; NP, near-projecting; CT, corticothalamic). (C) Violin plot showing the marker gene expression levels across mPFC neuron clusters. (D) Dot plot indicating hormone receptor gene expression in each mPFC neural cluster (Esr1, estrogen receptor 1; Esr2, estrogen receptor 2; Pgr, progesterone receptor; Gper1, G protein-coupled estrogen receptor 1; Ar, androgen receptor). (E) Heatmap showing ΔIEG^high^ cell percent within different estrus/sex comparison (estrus vs diestrus, estrus vs male, diestrus vs male) for each neuron cluster. (F) Schematic illustration of anterograde transsynaptic tracing of OxtrINs in the mPFC (left). Right: representative images depicting H129ΔTK-TT virus-labeled output neurons that are targeted by inputs from OxtrIN-innervated projection neurons in the mPFC. These downstream outputs are located within the lateral hypothalamus (LH), the central part of anterior hypothalamic nucleus (AHNc), parataenial thalamic nucleus (PT), mediodorsal nucleus of the thalamus (MD), anteromedial thalamic nucleus (AM), basolateral amygdala (BLA), basomedial amygdala (BMA), claustrum (CLA), and the anterior piriform cortex (APC). Scale bar, 200 µm. DAPI, blue. (G) Quantification of c-Fos+ cell counts across different brain regions (n = 3 mice for each group. Two-tailed unpaired t-test, * p < 0.05, ** p < 0.01). (H) Immunofluorescence images of c-Fos expression in the AHNc activated by sociosexual interaction during diestrus (left) and estrus (right). Scale bar, 200 µm. (I) Schematic diagram of the strategy for measuring synaptic connectivity between OxtrINs and mPFC^AHN-projecting^ neurons (left). Right: percentage of responsive cells in AHN-projecting and non-AHN-projecting layer 5 pyramidal neurons upon OxtrINs activation. N = 21 to 29 cells per group. (J) Example traces showing the light-evoked monosynaptic excitatory and inhibitory connectivity between OxtrINs and mPFC^AHN-projecting^ neurons (left). Comparison of the amplitudes of light-evoked IPSCs and EPSCs in mPFC^AHNc-projecting^ neurons between estrus and diestrus females (right). N = 60 to 67 cells recorded per group (right). Two-way ANOVA, Bonferroni multiple comparisons tests, n.s. = not significant, p > 0.05. (K) Summarized data of RMP recorded from mPFC^AHNc-projecting^ neurons. n = 8 to 24 cells per group. Two-way ANOVA, Bonferroni’s multiple comparisons test. * p < 0.05, ** p < 0.01, *** p < 0.001. (L) sEPSC recording in AHNc-projecting and non-AHNc-projecting mPFC neurons. Cumulative probability of frequency (left) and amplitude (middle) and representative traces (right) of sEPSCs. Insets in cumulative probability graphs represent average sEPSC frequency (left) and amplitudes (right). One-way ANOVA, Bonferroni multiple comparisons test, * p < 0.05, ** p < 0.01, *** p < 0.001. (M) Schematic of profiling translating mRNAs in mPFC^AHNc-projecting^ neurons using pathway-specific TRAPseq. AHNc infected with retrograde AAV2 virus expressing EGFP-tagged Rpl10a (left). Right: scatterplot of TRAP-seq results showing normalized counts from INPUT (x-axis) and IP (y-axis). Dots represent individual genes (blue, significantly enriched genes in IP; red dot, *Cacna1h* gene) (N) Coronal sections showing the expression of EGFP-Rpl10a (green) and *Cacna1h* mRNA (magenta) in the mPFC. Scale bar, 200 µm. (O) Violin plot showing *Cacna1h* gene expression across mPFC neuronal types.

To elucidate the impact of ovarian hormone fluctuations on neural activity indicated by transcriptomic changes, we analyzed the expression of immediate early gene (IEG) networks known to be regulated by ovarian hormones^38–40^. L5_PT neurons exhibited the most dramatic IEG network changes between estrus and diestrus, indicating high sensitivity in neural activity to ovarian hormone changes during the diestrus-to-estrus transition (Figure 1E). Functional implications were investigated by screening DEG clusters between estrus and diestrus specific to each cell type using hierarchical clustering and GO-term analysis. Intriguingly, among the top 10 enriched terms in L5_PT neurons were synaptic organization, protein phosphorylation, and GTPase activity (Figure S1I), indicating that estrous cycle transitions primarily affect synaptic plasticity of this cell type. Moreover, L5_PT neurons displayed the most DEGs between diestrus females and males, suggesting their sex-specific functions (Figure S1F). As L5_PT neurons primarily project to the hypothalamus ^34–37^, further suggesting their role in social behaviors.

### *Cacna1h*-defined L5_PT neurons relaying OxtrINs and AHN sense estrous status by strengthening synaptic efficacy under hyperpolarization

To identify estrus-sensitive cell types involved in female sociosexual behaviors, we employed additional multi-faceted screening approaches. Previous work reported that OxtrINs in the mPFC are essential for female sociosexual preference toward males during estrus^3^. In females, OxtrINs potentiated inhibition on postsynaptic layer 5 pyramidal neurons compared to males^41^. As Oxtr mRNA expression remains constant across the estrous cycle (Figure S4D), we hypothesized that layer 5 pyramidal neurons, which receive input from OxtrINs, rather than presynaptic OxtrINs, may sense estrous states and mediate estrus-dependent sociosexual behaviors in females.

To characterize neural populations connected with OxtrINs, we expressed an anterograde transsynaptic tracer, H129DTK-TT^42^, in the mPFC of Oxtr-Cre ON82 transgenic mice^3^ (Figure 1F). Dense mCherry+ viral signals were observed in subcortical regions (Figures 1F and Figure S2A), with significantly increased c-Fos+ cells evoked by sociosexual interactions in the central part of the anterior hypothalamic nucleus (AHNc) of estrus females compared to diestrus females (Figures 1G-1H). Notably, almost 100% of layer 5 mPFC neurons which project to AHN are responsive to OxtrIN activations (Figure 1I). The AHN, a component of a sexually dimorphic nucleus complex in many species^43–49^, is crucial for female rats’ sexual behavior^50^ and ovulation^51^. Bilateral lesions disrupt female sexual behavior^50^, while blocking AHN muscarinic receptors prevents ovulation by suppressing ovarian follicular formation^51^, These results suggest that OxtrINs-innervated, AHNc-projecting layer 5 neurons in the mPFC (mPFC^AHNc-projecting^) may play an estrus-dependent role in sociosexual behaviors.

To assess whether the electrophysiological properties of mPFC^AHNc-projecting^ neurons change over the estrous cycle, we performed slice physiology recordings. Optogenetic stimulation of OxtrINs elicited predominantly inhibitory postsynaptic responses in mPFC^AHNc-projecting^ neurons in the presence of tetrodotoxin (TTX) and 4-aminopyridine (4-AP), confirming their monosynaptic connectivity (Figure 1J). The synaptic strength (Figure 1J) and neurotransmitter release probability from presynaptic OxtrINs to mPFC^AHNc-projecting^ neurons (Figure S4C) were similar between estrus and diestrus, suggesting that postsynaptic mPFC^AHNc-projecting^ neurons, rather than presynaptic OxtrINs, may primarily track estrous states in females.

Notably, the resting membrane potential (RMP) of mPFC^AHNc-projecting^ neurons was significantly hyperpolarized in estrus females compared to diestrus females and males (Figure 1K). As *Oxtr+* mRNA was predominantly enriched in inhibitory Sst+ neurons (Figure S3A-S3D), oxytocin treatment selectively hyperpolarized these neurons in diestrus females but had no further effect in estrus females (Figure 1K), suggesting oxytocin’s inhibitory impact and an already hyperpolarized state during estrus. Intriguingly, the hyperpolarized mPFC^AHNc-projecting^ neurons displayed enhanced synaptic efficacy, with potentiated sEPSC frequency and amplitude specifically in estrus females but not diestrus females or in males (Figure 1L). The estrus-dependent enhancement of synaptic strength in hyperpolarized mPFC^AHNc-projecting^ neurons was not observed in non-AHN projecting layer 5 pyramidal neurons (Figure 1L), highlighting their unique properties across the estrous cycle.

To identify candidate genes in mPFC^AHNc-projecting^ neurons that could explain estrus-dependent electrophysiological changes, we employed a pathway-specific translating ribosome affinity purification (TRAP) approach to profile the translating mRNAs of these neurons (Figure 1K). We applied three criteria: (1) specific expression in deep mPFC layers; (2) possible link to enhanced synaptic transmission under hyperpolarization; and (3) implication in social behavior. Among the enriched genes, calcium voltage-gated channel subunit alpha1H (*Cacna1h*) emerged as the only candidate fulfilling all criteria (Figure 1L). *Cacna1h*, highly expressed in layer 5 pyramidal neurons of the mPFC (Figures 1M-1N), encodes the low-voltage-activated T-type calcium channel, Cav3.2 subunit, which mediates calcium influx near RMP^52,53^. Previous research indicates that *Cacna1h* mediates enhanced synaptic strength induced by hyperpolarization^54^ and synaptic upscaling activated by TTX-mediated suppression^55^. Moreover, *Cacna1h* mutations are associated with autism spectrum disorders in humans^56,57^. Given the evidence above, we hypothesized that *Cacna1h* might contribute to the synaptic strength potentiation in hyperpolarized mPFC^AHNc-projecting^ neurons during estrus.

Single-cell RNA sequencing (sc-RNAseq) revealed that *Cacna1h* is highly expressed in L5_PT neurons, serving as a gene marker defining this cell type (Figures 1M-1N). Notably, L5_PT neurons exhibit the most pronounced changes in DEGs (Figure S1F), the ovarian hormone-regulated IEG network (Figure 1E), and synapse-related gene modules between estrous cycle transitions (Figure S1I). Furthermore, the electrophysiological properties of mPFC^Cacna1h+^ neurons closely resemble mPFC^AHNc-projecting^ neurons during estrus, characterized by enhanced synaptic efficacy and hyperpolarized membrane potential (Figures S5C-S5F). Taken together, these results indicate that OxtrIN-innervated, *Cacna1h*-expressing L5_PT neurons projecting to the AHN are highly sensitive to changes in estrous status and may regulate estrus-dependent sociosexual behaviors.

### mPFC^Cacna1h+^ neurons bidirectionally regulate sociosexual behaviors in estrus-dependent and sexually dimorphic manners

To elucidate the impact of mPFC^Cacna1h+^ neurons on opposite-sex sociosexual behaviors, we employed a chemogenetic approach to selectively inhibit these neurons in the mPFC (Figures 2A-2C). Male and female Cacna1h-IRES-Cre mice, injected with either AAV2-DIO-mCherry or AAV2-DIO-hM4Di viruses, underwent several behavioral assays assessing sociosexual interests and sexual behavior. Post-behavioral assessment, estrous cycle phases of experimental females were determined via blind assignment following vaginal cytology (Figure 2D).

**Figure 2.**
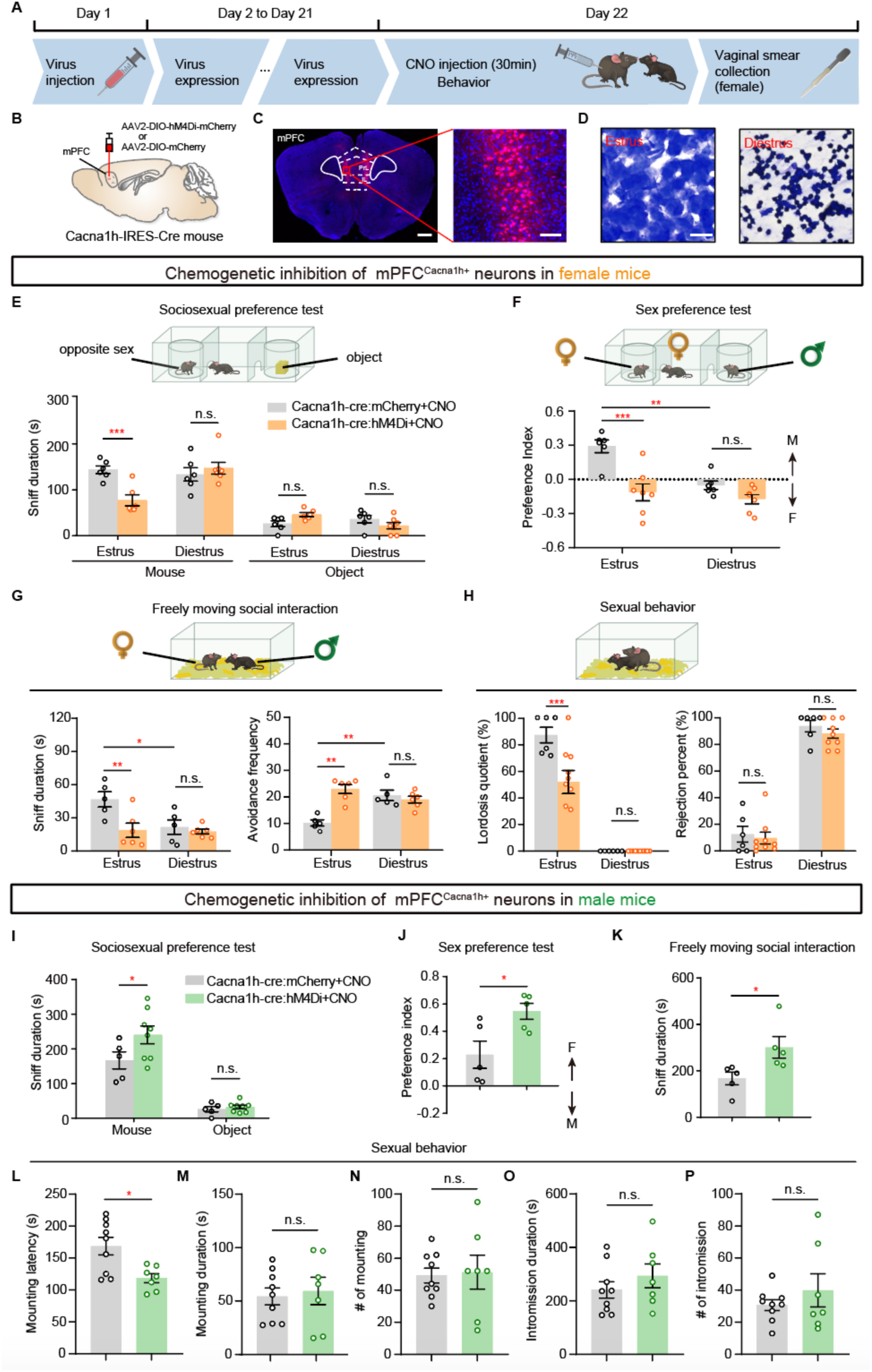
Chemogenetic inhibition of mPFC^Cacna1h+^ neurons suppresses sociosexual behaviors in estrus females but enhances them in males. (A) Timeline of behavioral testing for female and male Cacna1h-IRES-cre mice. (B) Schematic of virus injection into the mPFC of Cacna1h-IRES-cre mice. (C) Representative coronal section showing hM4Di-mCherry expression (magenta) in the mPFC^Cacna1h+^ neurons. Scale bars, 500 µm (left) and 100 µm (right). (D) Representative images of estrus and diestrus stages. Scale bar, 25 µm. (E) Sociosexual preference assay (top). Quantification of sniffing behaviors towards male or object (bottom). N = 6 to 7 mice per group. Two-way ANOVA, Bonferroni’s multiple comparisons test, ** p < 0.01. (F) Sex preference assay (top). Quantification of sniffing preference index towards male or female stimuli (bottom). N = 6 to 7 mice per group. Two-way ANOVA, Bonferroni’s multiple comparisons test, ** p < 0.01, *** p < 0.001. (G) Freely moving opposite-sex social interaction assay (top). Quantification of female-initiated social interactions (bottom left) and frequency of female avoidance responses to male approaches (bottom right). N = 5 to 6 mice per group. Two-way ANOVA, Bonferroni’s multiple comparisons test, * p < 0.05, ** p < 0.01. (H) Female sexual receptivity assessment (top). Quantification of the lordosis quotient (bottom left) and the percentage of male mating attempts rejected by females (bottom right). N = 6 to 9 mice per group. Two-way ANOVA, Bonferroni’s multiple comparisons test, *** p < 0.001. (I-K) Quantification of male social behaviors. Sniffing behaviors towards female or object in the sociosexual preference assay (I), sniffing preference index for male versus female stimuli in the sex preference assay (J), and male-initiated social interaction duration with females during freely moving social interaction (K). N = 5 to 8 mice per group. Two-way ANOVA, Bonferroni’s multiple comparisons test, * p < 0.05. (L-P) Quantification of male sexual behavior parameters, including mounting latency (L), mounting duration (M), number of mounts (N), intromission duration (O), and number of intromissions (P). N = 7 to 9 mice per group. Two-way ANOVA, Bonferroni’s multiple comparisons test, * p < 0.05.

In sociosexual preference assays, female mice were presented with a choice between a cup-restrained mouse of opposite sex and a novel object. Silencing mPFC^Cacna1h+^ neurons elicited a 47% reduction in male-directed sniffing duration specifically in estrus females without affecting other estrous phases (Figures 2E and S7C). Moreover-estrus females demonstrated a marked preference for sexually experienced males over females during the sex preference test, which was absent in diestrus females (Figure 2F). Remarkably, chemogenetic inhibition of mPFC^Cacna1h+^ neurons significantly attenuated the estrus-driven preference for males, reducing the preference index by 139% in hM4Di-expressing estrus females compared to controls (Figure 2F). During freely social interactions, females in estrus exhibited significantly increased investigation time and markedly reduced avoidance behaviors towards males compared to diestrus females (Figure 2G). Silencing mPFC^Cacna1h+^ neurons in estrus females specifically reduced female-initiated interactions with males, and increased avoidance frequency in response to male pursuits (Figure 2G). Furthermore, to investigate the role of mPFC^Cacna1h+^ neurons in modulating sexual receptivity, females across different estrous stages were introduced to sexually primed males. Inhibiting these neurons diminished the heightened sexual receptivity observed in estrus-phase females, as indicated by a reduced lordosis quotient (Figure 2H).

Conversely, in males, mPFC^Cacna1h+^ neuron suppression increased female-directed investigation in both sociosexual and sex preference assays, dramatically elevating interaction with freely moving females (Figures 2I-2K). Moreover, chemogenetic inhibition of mPFC^Cacna1h+^ neurons in males led to a 30% reduction in the latency to initiate mounting (Figure 2L), suggesting heightened sexual motivation. However, mPFC^Cacna1h+^ neuron suppression did not significantly affect the number of mountings, intromission duration, or ejaculation latency in males (Figures 2M-2P).

To determine if mPFC^Cacna1h+^ neurons are sufficient for the increased sociosexual interests and sexual receptivity in females, we chemogenetically activated these neurons by expressing AAV2-DIO-hM3Dq viruses in the mPFC of Cacna1h-IRES-cre mice. This intervention augmented female-initiated social investigation towards males across estrus and diestrus phases, and enabled diestrus females to exhibit estrus-like male preference (Figures 3A-3B). During freely social interactions, activating mPFC^Cacna1h+^ neurons in diestrus females enhanced their male investigation and reduced avoidance of male approaches (Figure 3C). Furthermore, enhanced mPFC^Cacna1h+^ neuronal activity also significantly reduced rejection rates in diestrus females during male mounting attempts (Figure 3D). In males, however, mPFC^Cacna1h+^ neuron activation significantly reduced the interaction duration with females without affecting their preference to female targets (Figures 3E-3G). Notably, enhanced mPFC^Cacna1h+^ neural activity dramatically suppressed male sexual behaviors, including mounting and intromission duration and frequency, along with a marked reduction in ejaculation incidence (Figures 3H-3L).

**Figure 3.**
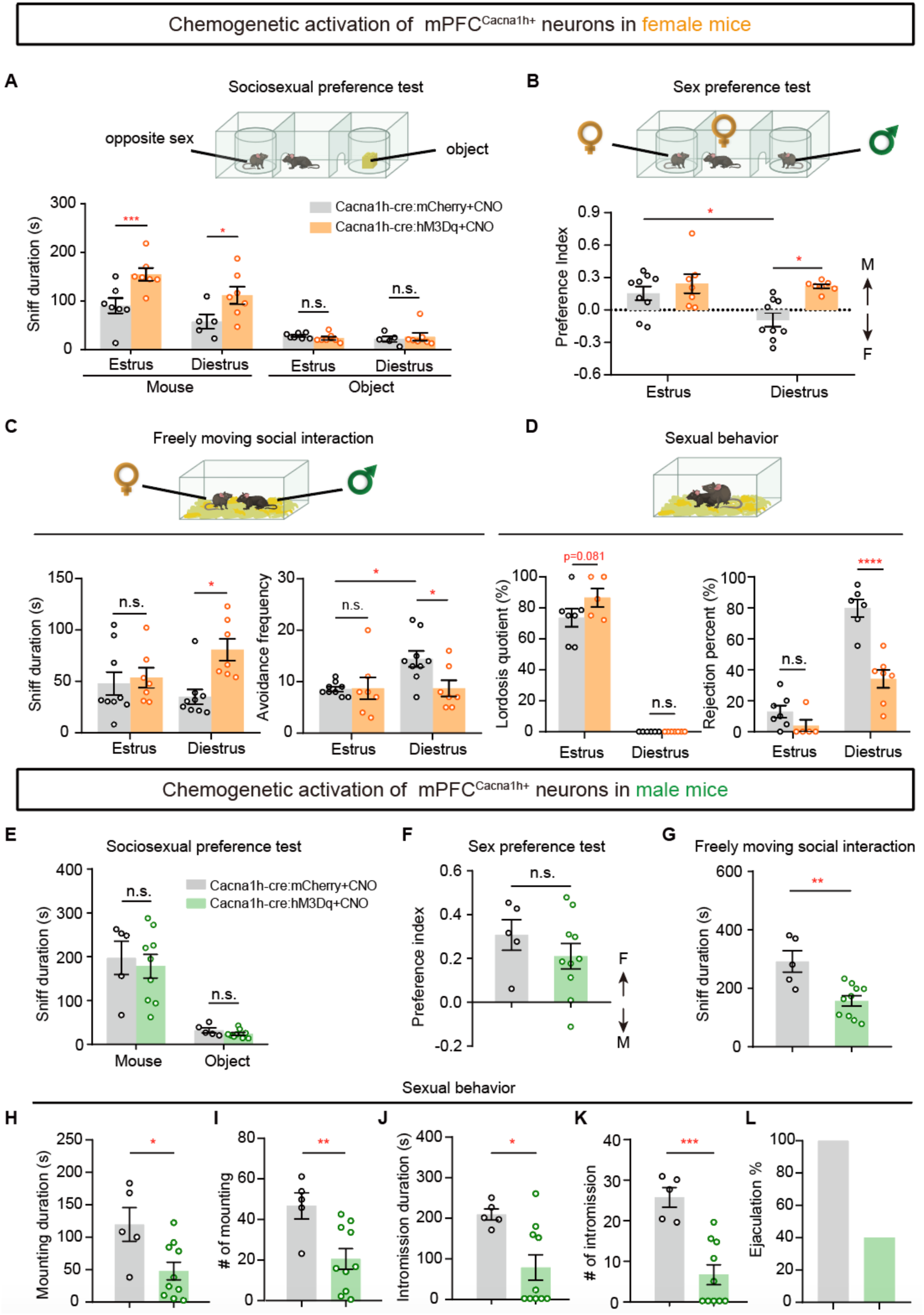
Chemogenetic activation of mPFC^Cacna1h+^ neurons elevates sociosexual behaviors in diestrus females but suppresses them in males. (A) Sociosexual preference assay. Quantification of sniffing behaviors towards male or object in estrus and diestrus females. N = 5 to 7 mice for each group. Two-way ANOVA, Bonferroni multiple comparisons test, * p < 0.05, *** p < 0.001. (B) Sex preference assay. Quantification of sniffing preference index towards male or female stimuli. N = 7 to 9 mice for each group. Two-way ANOVA, Bonferroni multiple comparisons test, * p < 0.05. (C) Opposite-sex social interaction assay. Quantification of female-initiated social interactions (Bottom Left) and frequency of female avoidance responses to male approaches (Bottom Right). N = 7 to 9 mice for each group. Two-way ANOVA, Bonferroni multiple comparisons test, n.s. = not significant, p > 0.05; * p < 0.05. (D) Female sexual receptivity assessment. Quantification of the lordosis quotient (Bottom left) and the percentage of male mating attempts rejected by females (Bottom right). N = 3 to 7 mice for each group. Two-way ANOVA, Bonferroni multiple comparisons test, * p < 0.05. (E-G) Quantification of male social behaviors. Sniffing behaviors towards female or object in the sociosexual preference assay (E), sniffing preference index in sex preference assay (F) and male-initiated interaction duration during social interaction assay (G). N = 5 to 10 mice for each group. Two-way ANOVA, Bonferroni multiple comparisons test, n.s. = not significant, p > 0.05, ** p < 0.01. (H-L) Quantification of male sexual behavior. Parameters analyzed including mounting duration (H), numbers of mounts (I), intromission duration (J), number of intromissions (K) and percentage of ejaculation (L). N = 5 to 10 mice for each group. Two-way ANOVA, Bonferroni multiple comparisons test, * p < 0.05, ** p < 0.01, *** p < 0.001.

Collectively, these findings unveil the pivotal role of mPFC^Cacna1h+^ neurons in orchestrating sociosexual behaviors, including opposite-sex preference, interaction and copulatory behaviors, in estrus-specific and sexually dimorphic manners.

### The representation of target sex by mPFC^Cacna1h+^ neurons is estrus-dependent and sex-specific

To investigate the neural response of mPFC^Cacna1h+^ neurons during sociosexual preference behavior, fiber photometry recordings were performed in Cacna1h-IRES-Cre mice with AAV-DIO-GCaMP6f virus infected into the mPFC (Figures S9A-S9B). These recordings revealed distinct mPFC^Cacna1h+^ neuron responses to opposite-sex conspecifics and novel objects. In estrus females, mPFC^Cacna1h+^ neurons showed a decrease in activity prior to sniffing males, followed by a fivefold increase during sniffing compared to diestrus (Figures S9C-S9D). This response was absent during novel object exploration (Figures S9E-S9F). In contrast, males displayed a modest decreasing trend in activity when sniffing females (Figures S9H).

To further dissect the estrus- and sex-dependent encoding mechanisms of social cues by mPFC^Cacna1h+^ neurons at the single-neuron level, we performed micro-endoscopic calcium imaging using a GRIN lens in Cacna1h-IRES-Cre mice of both sexes (Figure 4A). The calcium dynamics of GCaMP7b-expressing mPFC^Cacna1h+^ neurons were recorded during sociosexual and sex preference behaviors (Figures 4D and 4H). Individual cell activity was extracted with Minian based on constrained non-negative matrix factorization (CNMF) algorithm^58^, and reported as relative change in fluorescence (ΔF/F) (Figures 4B-4C).

**Figure 4.**
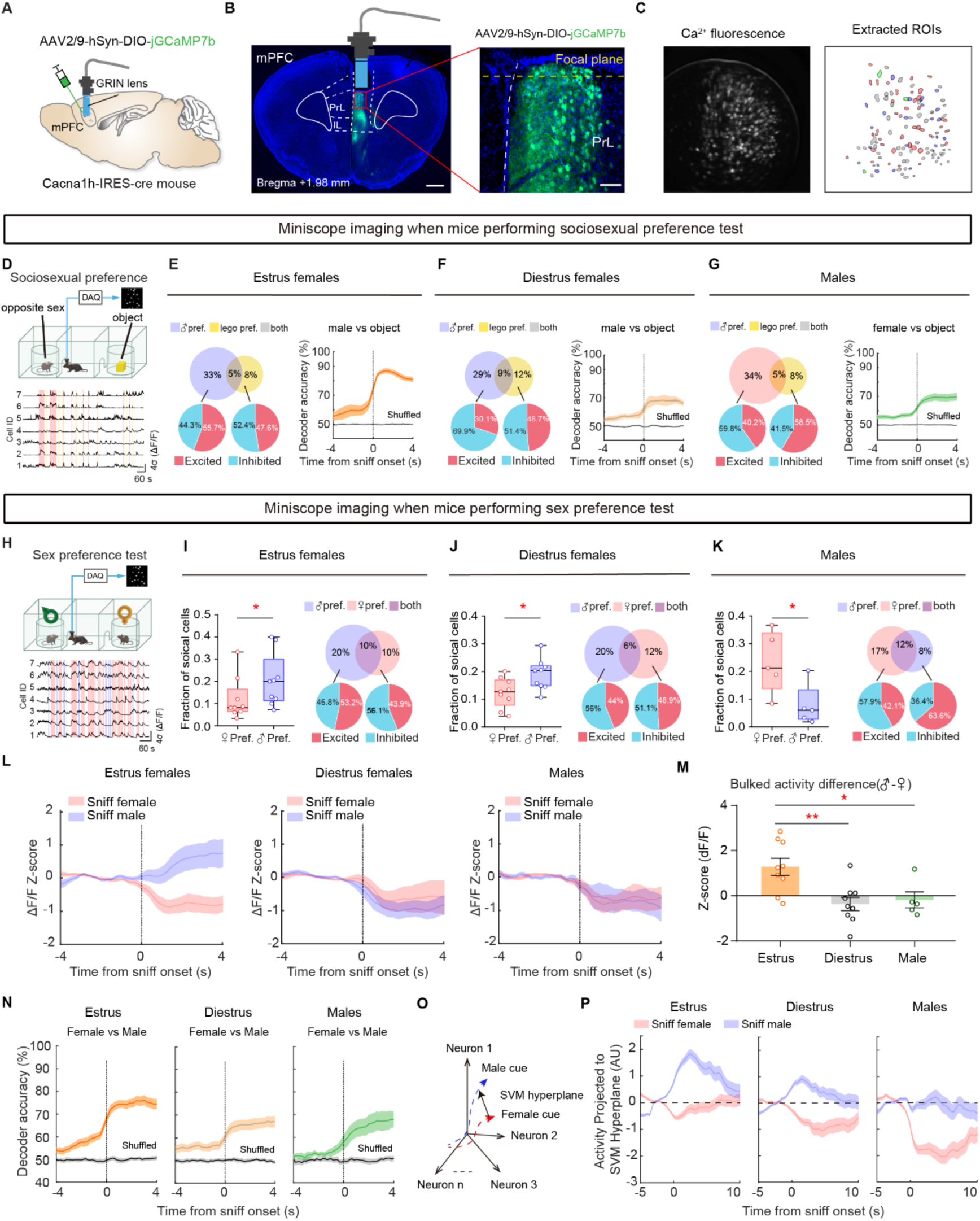
Estrus- and sex-specific representation of target-sex information by mPFC^Cacna1h+^ neurons. (A) Schematic of AAV2/9-hSyn-DIO-jGCaMP7b injection into the mPFC of Cacna1h-IRES-cre mice. (B) Expression of AAV2/9-hSyn-DIO-jGCaMP7b in the mPFC with a GRIN lens implanted above the infection site (left). GFP-positive cells represent GCaMP7b-expressing mPFC^Cacna1h+^ neurons (right). Blue, DAPI; scale bars, 500 µm (left) and 20 µm (right). (C) Raw calcium fluorescence from a single imaging field (left). Extracted ROIs corresponding to single neurons from the imaging field (right). (D) Sociosexual preference test paradigm (top) and representative calcium traces from mPFC^Cacna1h+^ neurons (bottom). (E-G) Fraction of mPFC^Cacna1h+^ neurons responding to sniffing mouse of opposite sex versus object in estrus females (E left, n = 510 neurons from 8 mice), diestrus females (F left, n = 319 neurons from 5 mice), and males (G left, n = 703 neurons from 10 mice). Time-wise classifier performance for distinguishing between sniffing male and object (-4 to 4s relative to sniff onset; E-G, right). (H) Sex preference test paradigm (top) and representative calcium traces from mPFC^Cacna1h+^ neurons (bottom). (I-K) Fraction of mPFC^Cacna1h+^ neurons responding to sniffing female versus male in individual (left) and pooled (right) estrus females (I, n = 394 neurons from 9 mice), diestrus females (J, n = 384 neurons from 9 mice), and males (K, n = 266 neurons from 5 mice). Pie charts show the percentage of neurons preferring female, male, or both. (L) Averaged Z-scored calcium signal aligned to sniff onset for investigating female and male trials in estrus females (n=9, left), diestrus females (n=9, middle), and males (n=5, right). (M) Quantification of Z-scored calcium signal difference between sniffing male and female trials (0-4s post-sniff onset) in estrus females, diestrus females, and males. (N) Time-wise classifier performance for distinguishing between sniffing male and female (-4 to 4s relative to sniff onset) in estrus females (left), diestrus females (middle), and males (right). (O) Schematic of mPFC^Cacna1h+^ neural activity projection onto an SVM-optimized hyperplane during male/female investigation. (P) mPFC^Cacna1h+^ neural activity during sniffing male (blue) and female (red) trials projected onto the SVM hyperplane in estrus females, diestrus females, and males.

During sociosexual preference tests (Figure 4D), mPFC^Cacna1h+^ neurons responsive to opposite-sex targets significantly outnumbered those responding to non-social objects in both sexes (Figures 4E-4G). Intriguingly, while the proportions of socially-tuned neurons were similar across groups, opposite-sex stimuli induced a higher percentage of suppression in diestrus females and males compared to estrus females (Figures 4E-4G). In contrast, male-responsive mPFC^Cacna1h+^ neurons were predominantly activated during estrus, indicating an estrus state-specific shift in the excitation/inhibition (E/I) balance towards excitation (Figure 4E). To assess the encoding capabilities of mPFC^Cacna1h+^ neurons, we trained a linear support vector machine (SVM) classifier to differentiate between opposite-sex and object trials based on population activity. Classifiers trained with estrus state data significantly outperformed those trained with diestrus and male data, and this superior encoding was consistently observed in pseudo-population analyses controlling for ensemble size effects (Figures 4E-4G, S11D-S11E), highlighting the estrus-dependent more effective encoding of male cues by mPFC^Cacna1h+^ neurons.

To investigate the neural correlates of sex-specific social cues in the mPFC^Cacna1h+^ neurons, we analyzed the calcium dynamics of these neurons during animals’ investigation of male and female targets in sex preference tests (Figure 4H). Across all groups, a higher proportion of neurons responded to opposite-sex compared to same-sex targets (Figures 4I-4K). Notably, in estrus females, opposite-sex cues excited mPFC^Cacna1h+^ neurons, while in diestrus females and males, opposite-sex cues predominantly suppressed their activity, aligning with the observations in sociosexual preference assay (Figures 4I-4L). Quantification of the averaged neuronal activity confirmed the selective response to male cues in estrus females, with significantly higher activation during male investigation trials (Figures 4L-4M). In contrast, the averaged neuronal activity in males showed decreased responses during female investigation trials (Figures 4L).

To investigate population-level encoding of social target sex, we trained a SVM classifier to discriminate between male and female investigation trials based on neural activity around sniff onset. Classification accuracy was above chance for all groups, with the highest performance achieved in estrus females (Figure 4N). Projecting the neural activity onto the SVM hyperplane, which maximally separated male and female trials, revealed that the discrimination was primarily driven by opposite-sex responses in estrus females and males (Figures 4O-4P). Same-sex preferring neurons in males exhibited a higher auROC bias towards opposite-sex targets compared to diestrus females (Figure S11J). Furthermore, dimensionality reduction using PCA demonstrated a more distinct separation of opposite-sex trials from baseline compared to same-sex trials in both estrus females and males (Figures S11K-S11L). These analyses unveil a significant bias in the tuning direction of mPFC^Cacna1h+^ neurons towards opposite-sex cues in estrus females and males. Collectively, these findings demonstrate that mPFC^Cacna1h+^ neurons encode social cues in an estrus- and sex-specific manner, with opposite-sex cues driving increased responses in estrus females and predominantly suppressed responses in males. The estrus-dependent excitatory response to male cues and the biased tuning towards opposite-sex targets in mPFC^Cacna1h+^ neurons suggest a neural substrate for male-directed preferences in estrus females. The differential encoding of opposite-sex cues by mPFC^Cacna1h+^ neurons in estrus females and males may underlie the sexually dimorphic modulation of sociosexual behaviors.

### Distinct mPFC^Cacna1h+^ neuron subpopulations engage in both representing self-estrous states and distinguishing sex of sniffing targets

To determine whether mPFC^Cacna1h+^ neurons encode self-estrous states in females, given their estrus-dependent representation of target-sex information, we analyzed calcium imaging data collected at three stages: baseline in home cages, habituation in a three-chamber apparatus, and investigation of social targets of both sexes (Figure 5A). The aligned mPFC^Cacna1h+^ neurons, imaged across estrus and diestrus phases in the same female mouse, underwent further analysis (Figure 5B). These results with aligned neurons across estrous states mirrored the previous analysis with all neurons, revealing a significant increase in estrus females relative to their diestrus phases during male target sniffing, and superior performance of the linear SVM decoder in predicting male versus female directed sniffing behaviors using estrus data (Figures 5D-5E).

**Figure 5.**
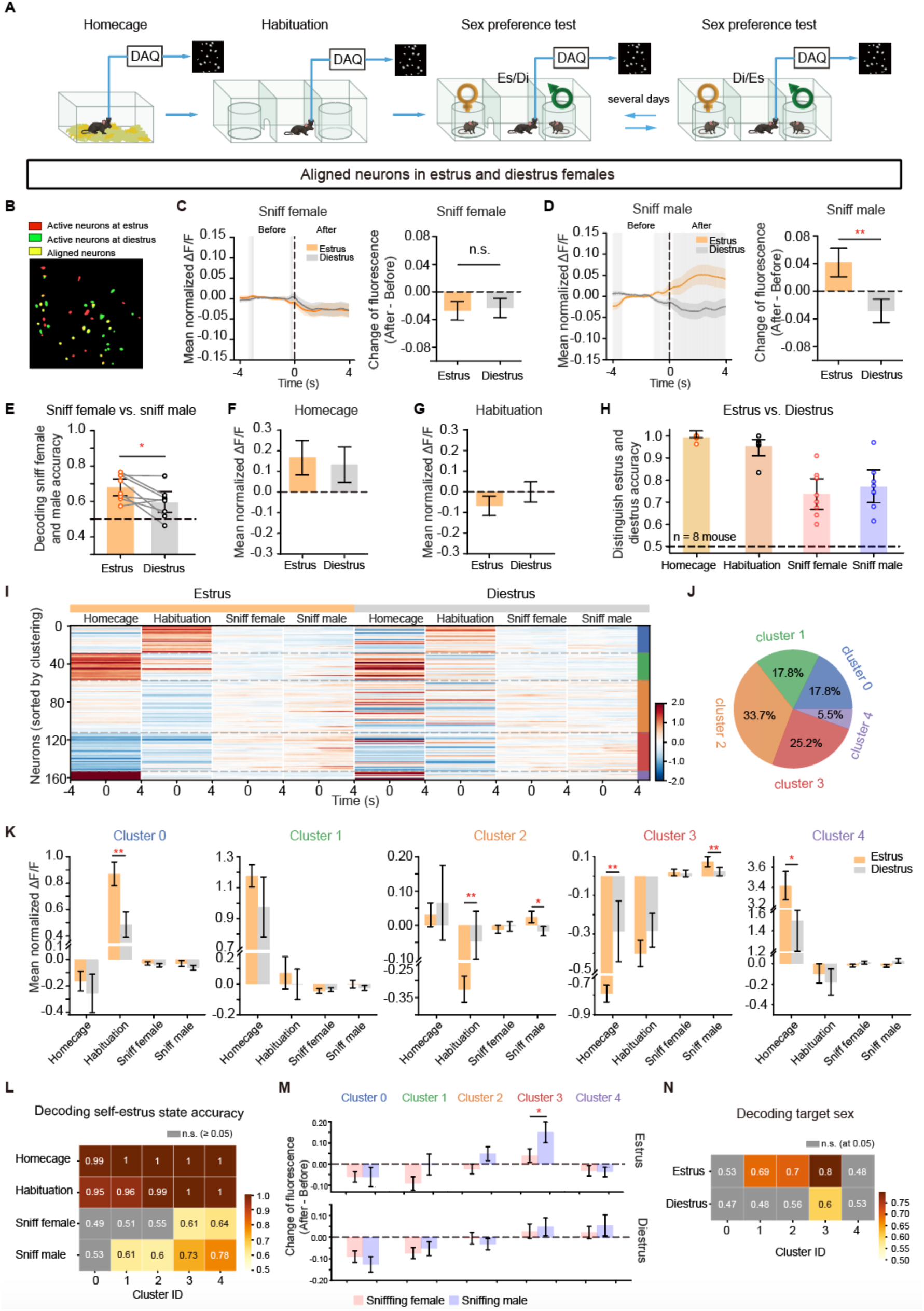
Distinct mPFC^Cacna1h+^ neuron subpopulations engage in both representing self-estrous states and distinguishing sex of sniffing targets. (A) Experimental design for calcium imaging of mPFC^Cacna1h+^ neurons in female mice across three distinct stages: home cage baseline, habituation to a three-chamber apparatus, and investigation of male and female social targets. Calcium imaging was performed during both the estrus and diestrus phases in the same female mouse. (B) Representative calcium imaging fields of view showing aligned mPFC^Cacna1h+^ neurons recorded in estrus and diestrus females. (C-D) Calcium activity of aligned mPFC^Cacna1h+^ neurons during sniffing trials in estrus and diestrus. (C) Left: Average calcium traces of aligned mPFC^Cacna1h+^ neurons during sniff-female trials. Right: Fluorescence change ratio before and after behavior onset for sniff-female trials. (D) Left: Average calcium traces of aligned neurons during sniff-male trials. Right: Fluorescence change ratio before and after behavior onset for sniff-male trials. Colored shaded areas: SEM. Gray shading: time points with significant differences between estrus and diestrus (n = 8 mice, **, p = 0.0006). (E) Decoding accuracy of sniffing target sex using aligned mPFC^Cacna1h+^ neural activity in estrus and diestrus. (n = 8 mice, *, p = 0.0142, Wilcoxon matched-pairs signed-rank test). (F-G) Spontaneous activity of aligned mPFC^Cacna1h+^ neurons in home cage (F) and 3-chamber habituation (G) (n = 162 neurons from 8 mice). (H) Decoding self-estrous states from mPFC^Cacna1h+^ neural activity across behavioral stages. Decoder trained on calcium activity predicted self-estrous state above chance level during all stages. (I) Clustering of mPFC^Cacna1h+^ neurons based on activity profiles across behavioral stages. Heatmap shows mean activity profiles of clustered neurons, with rows representing single neurons and column representing behavioral stages. Color scale indicates normalized mean activity; colors on the right axis denote different clusters; colors on the top distinguish estrus and diestrus. (J) Proportion of mPFC^Cacna1h+^ neurons assigned to each cluster. The color of each slice corresponds to the cluster colors in Figure I. (K) Mean activity of clusters in estrus and diestrus states during different behavioral stages. (L) Decoding accuracy of self-estrous state using clusters across behavioral contexts. Heatmap showing decoding accuracy for each cluster (columns) and behavior (rows). Colored entries: p < 0.05; grey entries: non-significant. Color bar indicates accuracy range. (M) Fluorescence change ratio of clusters before and after the onset of sniff-female and sniff-male events in estrus (top) and diestrus (bottom) states. Data: mean ± SEM; *, p < 0.05. (N) Decoding accuracy of sniffing target sex using mPFC^Cacna1h+^ neuron clusters in estrus and diestrus. Matrix depicting decoding accuracy of each cluster (columns) during different estrous states (rows). Colored entries: p < 0.05; grey entries: non-significant. Color bar indicates accuracy range.

The aligned mPFC^Cacna1h+^ neurons exhibited similar spontaneous activities across estrus and diestrus phases in both homecage and habituation stages (Figures 5F-5G), yet neural decoding analysis reliably distinguished between estrus and diestrus with notable accuracy (Figure 5H), highlighting their capability to encode self-estrous states. Despite similar mean spontaneous activities, their capacity to differentiate between these states indicates subpopulations with distinct activities specific to each estrous phase.

To address this, we employed K-means clustering (see Methods) to partition the aligned mPFC^Cacna1h+^ neurons recorded in all female mice into functionally similar subpopulations. The features of each neuron were constructed according to its activity across homecage, habituation, and social behavior sessions in both estrus and diestrus states. This approach enabled us to compare the ‘role’ of each neuron during all conditions and states. An optimal clustering result gives out K = 5 clusters (Figure S12B). The profile heatmap revealed distinct unique activity patterns among these clusters (Figure 5I), implying that distinct subpopulations may perform unique roles in encoding functions.

Analysis of mean activity of mPFC^Cacna1h+^ neurons by cluster, correlated with the self-estrus states, revealed marked differences in spontaneous activity between the estrus and diestrus states during homecage or habituation in most subpopulations (Figure 5K). Notably, clusters 2 and 3 demonstrated distinct activity differences during male sniffing behaviors between estrus and diestrus phases. Further decoding analysis of neural activity per cluster (Figures 5L and 5N) and fluorescence change ratio (Figures 5M) yielded consistent findings. Most neuronal clusters demonstrated optimal performance in decoding self-estrus states based on spontaneous activity data (Figure 5L). Notably, cluster 3 achieved the highest accuracy in decoding the sex of investigated target during the sex preference test (Figure 5N). Clusters 4 and 0 primarily encoded self-estrus state information, while clusters 3, 2, and 1 effectively decoded both self-estrus states and target-sex information (Figures 5L and 5N), indicating a complex representation of mixed information. The notably lower correlation R-value of all distinct subpopulations in responses to male versus female sniffing during estrus highlighted mPFC^Cacna1h+^ neurons’ improved sex differentiation in the estrus state compared to diestrus (Figures S12D-S12E). Moreover, clusters 1, 2, and 3 showed superior performance in target-sex decoding during estrus over diestrus (Figure 5N). Interestingly, subpopulations in males paralleled those in females (Figures S12F-S12H), suggesting a conserved subpopulation architecture in both sexes.

Collectively, these results robustly demonstrate that mPFC^Cacna1h+^ neurons effectively represent self-estrus states and distinguish the sex of sniffing targets, with specific subpopulations encoding distinct aspects of multifaceted information. These findings underscore the key role of the mPFC in integrating internal estrous states with external social cues, providing neural mechanism for top-down modulation of adaptive sociosexual behaviors.

### Ovarian hormone-induced *Cacna1h* upregulation drives estrus-specific activity changes of mPFC^Cacna1h+^ neurons

To investigate the molecular and cellular mechanisms underlying estrus-specific activity changes in mPFC^Cacna1h+^ neurons, we performed TRAPseq to identify the translational profiles of mPFC^Cacna1h+^ neurons across the estrous cycles^59^ (Figure 6A and S13A). Hierarchical clustering identified five gene clusters with distinct expression patterns (Figure 6B). Cluster 3 genes, exhibiting peak expression during estrus, were significantly enriched in female-specific functions related to reproduction, synaptic plasticity, and calcium ion-dependent exocytosis (Figures 6C and 6D). Notably, the fold enrichment of these gene ontology (GO) terms in cluster 3 was substantially higher than in other estrous phases (Figure 6D). This suggests a female-specific function of mPFC^Cacna1h+^ neurons related to sex and the estrous cycle.

**Figure 6.**
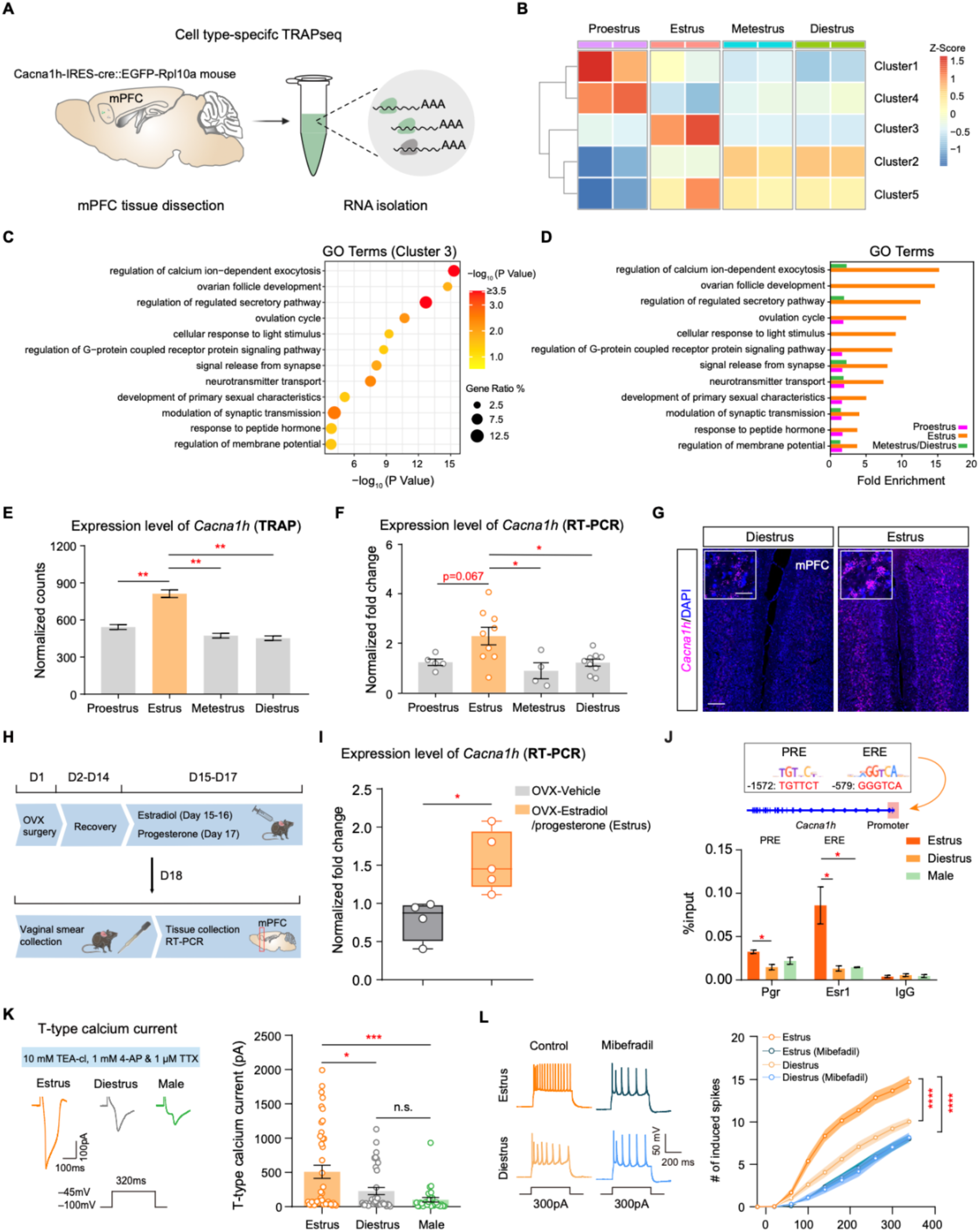
Ovarian hormone-induced *Cacna1h* upregulation drives estrus-specific activity of mPFC^Cacna1h+^ neurons. (A) TRAP profiling of translating mRNAs in mPFC^Cacna1h+^ neurons using the Cacna1h-IRES-Cre::EGFP-Rpl10a mouse line. (B) Hierarchical clustering of gene expression in mPFC^Cacna1h+^ neurons from females at four estrous phases. Five gene clusters are differentially expressed across the estrus cycle. (C) GO analysis of DEGs in cluster 3 of estrus-sensitive genes. (D) Comparative analysis of DEG enrichment in GO categories across estrous stages. Fold enrichment is shown as zero when the p-value is not significant. (E) TRAP counts of *Cacna1h* mRNA enrichment in mPFC^Cacna1h+^ neurons from females at different estrus phases (n = 2 samples per group; one-way ANOVA, Bonferroni’s multiple comparisons test; **p < 0.01). (F) RT-PCR of *Cacna1h* in the mPFC of females at four estrous stages (n = 4-9 mice per group; one-way ANOVA, Tukey’s multiple comparisons test; *p < 0.05). (G) Verification of *Cacna1h* levels in the mPFC of estrus and diestrus females using RNAscope. Scale bars, 50 µm [inset] and 200 µm. (H) Experimental timeline of ovariectomy (OVX) and ovarian hormone supplementation. (I) RT-PCR of *Cacna1h* in the mPFC of OVX females receiving ovarian hormones or vehicle (n = 4-6 mice per group; two-tailed unpaired t-test; *p < 0.05). (J) Chromatin immunoprecipitation (ChIP) analysis of estrogen receptor (ER) and progesterone receptor (PR) binding to hormone response elements in the *Cacna1h* promoter during the estrous cycle. The *Cacna1h* promoter contains predicted progesterone (PRE) and estrogen (ERE) response elements (top). Quantification of ER and PR binding affinity to EREs and PREs in estrus and diestrus females, represented as the percentage of input DNA recovered after immunoprecipitation with ER and PR antibodies (bottom). (K) T-type calcium current recording strategy (top). Representative traces of T-type calcium currents at -45 mV in mPFC^Cacna1h+^ neurons from estrus and diestrus females and males (bottom left). Quantitation of T-type current amplitudes (n = 29-42 cells per group; one-way ANOVA, Bonferroni’s multiple comparisons test; *p < 0.05, ***p < 0.001, n.s, p > 0.05) (right). (L) Representative traces of induced spikes from mPFC^Cacna1h+^ neurons with and without mibefradil in estrus and diestrus females (left). Summarized data of induced spikes (n = 22-33 cells per group; two-way ANOVA, Bonferroni’s multiple comparisons test; ****p < 0.0001) (right).

Interestingly, *Cacna1h* gene in cluster 3, which encodes the Cav3.2 T-type calcium channel and may drive the estrus-dependent synaptic potentiation under membrane hyperpolarization, reached its highest expression level during the estrus phase (Figure 6E), as confirmed by quantitative real-time PCR (RT-PCR) and RNAscope methods (Figures 6F-6G). Moreover, *Cacna1h* mRNA levels increased dramatically in ovariectomized female mice receiving ovarian hormones to induce estrus (Figures 6H-6I). The *Cacna1h* gene promoter region included hormone response elements binding estrogen and progesterone receptors, with significantly stronger affinity for Pgr and Esr1 in estrus females compared to diestrus females and males (Figure 6J). These findings indicate that ovarian hormones directly regulate *Cacna1h* expression via a genomic pathway during estrus. Notably, this dynamic expression pattern was specific to *Cacna1h* and not observed for the other T-type calcium channel genes, *Cacna1g* and *Cacna1i* (Figures S13C and S13D).

The estrus-specific *Cacna1h* upregulation suggested enhanced Cav3.2-mediated T-type currents in mPFC^Cacna1h+^ neurons. To confirm this, we measured the T-type calcium current amplitude in mPFC^Cacna1h+^ neurons and found that the amplitude was significantly larger in mPFC^Cacna1h+^ neurons of estrus females compared to diestrus females and males (Figure 6K). This increased calcium influx influenced mPFC^Cacna1h+^ neuron firing frequency, as evidenced by higher spike counts at each injected current step (>100 pA) during estrus. Blocking T-type currents with mibefradil reduced firing rates and eliminated the estrus-dependent difference in excitabilty (Figure 6L), demonstrating that Cav3.2-mediated T-type currents drive the estrus-specific hyperactivity of mPFC^Cacna1h+^ neurons.

The hyperactivity and hyperpolarized RMP of mPFC^Cacna1h+^ neurons during estrus suggest that T-type calcium current-induced rebound activation may contribute to this process. To investigate the rebound activity of mPFC^Cacna1h+^ neurons following inhibition from presynaptic OxtrINs in vivo, we optogenetically triggered oxytocin release by expressing ChR2 in the paraventricular nucleus (PVN) while recording the calcium activity of mPFC^Cacna1h+^ neurons using fiber photometry (Figure S14A). To isolate oxytocin’s primary effect mediated by OxtrINs, we selectively blocked excitatory synaptic transmission and CRF system functions, eliminating potential confounding factors including the activation of excitatory Oxtr+ neurons in the mPFC (Figure S14) and the influence of CRF peptide released from PVN neurons on mPFC neurons with abundant CRF receptor 1 expression^41^. Optogenetic stimulation of PVN induced two distinct increases in mPFC^Cacna1h+^ neuronal activity: an immediate peak at light onset, likely reflecting disinhibition among inhibitory synapses, and a subsequent rebound activity at light cessation (Figures S14B). This rebound activity was more pronounced in estrus than in diestrus (Figures S14D). Oxtr blockade reduced the amplitude of rebound activity in estrus (Figures S14D), underscoring the crucial role of oxytocin/OxtrINs in mediating the enhanced rebound activity of mPFC^Cacna1h+^ neuron during estrus.

Taken together, our findings reveal that the estrus-specific elevation of *Cacna1h*, driven by ovarian hormones through a genomic pathway, leads to a remarkable rise in T-type currents, culminating in estrus-specific alterations in the electrophysiological properties of mPFC^Cacna1h+^ neurons. Furthermore, oxytocin, acting through OxtrINs, modulates the rebound activity of mPFC^Cacna1h+^ neurons in an estrus-dependent manner. These findings provide the molecular and cellular mechanisms underlying the estrus-specific functions of mPFC^Cacna1h+^ neurons in females.

### *Cacna1h* in the mPFC mediates estrus-specific and sexually opposing regulation of sociosexual behaviors

To investigate the role of *Cacna1h* in sociosexual behaviors, we generated a conditional *Cacna1h* knockout (cKO) mouse line (Cacna1h^fl/fl^) using CRISPR/Cas9 genome editing (Figures S15A-S15B). Expressing cre virus in the mPFC of this mouse line significantly reduced *Cacna1h* expression (Figures S15C-S15E) and decreased T-type calcium current amplitudes to 16.7% of the control (Figure S15F), confirming efficient deletion of *Cacna1h* at both genetic and electrophysiological levels.

In females, mPFC-specific *Cacna1h* cKO resulted in a significant reduction in male-directed sniffing (Figure 7A) and decreased preference for male targets exclusively during estrus, without affecting females at other estrous stages (Figures 7C and S16B). Testing each female mouse across different estrous cycles revealed a consistent, estrus-phase-specific reduction in male-directed sniffing behavior (Figure 7B). In freely moving social encounters, estrus females with mPFC *Cacna1h* deletion exhibited reduced investigation of males and increased avoidance responses to male approaches (Figure 7D). Moreover, *Cacna1h* deletion in the mPFC significantly reduced sexual receptivity in estrus females, as indicated by a decreased lordosis quotient (Figure 7E). Notably, conditional knockout of *Cacna1h* in the female mPFC did not alter the overall length of the estrous cycle or the duration of individual phases (Figure 7F), suggesting that *Cacna1h* directly modulates sociosexual behaviors during estrus, rather than through alterations in the estrous cycle itself.

**Figure 7.**
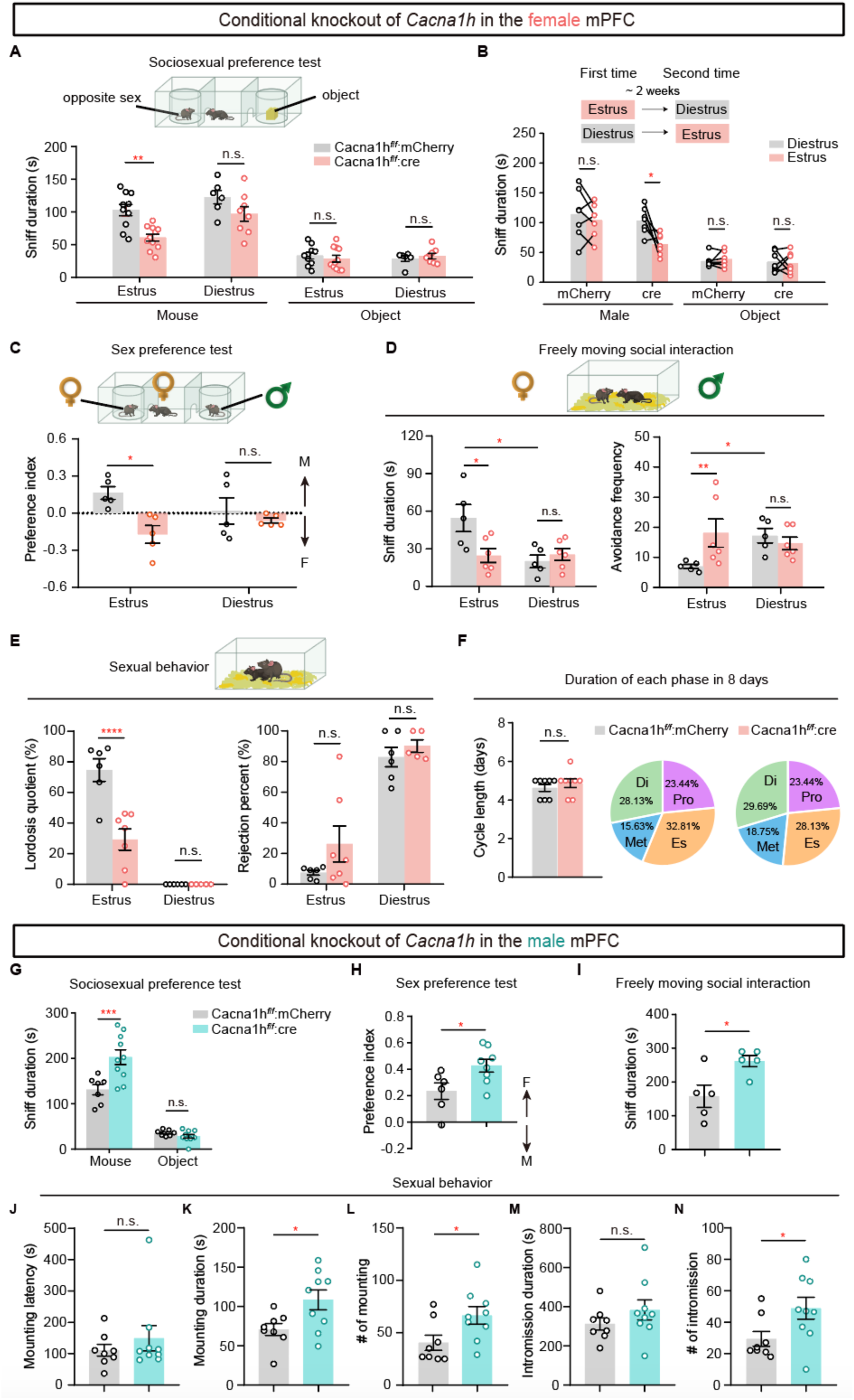
*Cacna1h* in the mPFC regulates sociosexual behaviors in estrus-specific and sexually dimorphic manners. (A) mPFC *Cacna1h* deletion in female mice decreased their sniffing time towards males during estrus, but not diestrus (n = 6-11 mice/group; Two-way ANOVA, Bonferroni multiple comparisons test, ** p < 0.01). (B) Each female mouse was examined twice for sociosexual preference during both estrus and diestrus phases, with a minimum two-week interval (n = 7 mice/group; Two-tailed paired t-test, * p < 0.05). (C) mPFC *Cacna1h* ablation in estrus females significantly reduced sex preference towards males (n = 5 mice/group; Two-way ANOVA, Bonferroni multiple comparisons test, ** p < 0.01). (D) Conditional knockout of *Cacna1h* in estrus females significantly reduced freely moving social interaction and increased avoidance frequency towards males (n = 5-6 mice/group; Two-way ANOVA, Bonferroni multiple comparisons test, * p < 0.05). (E) Ablation of *Cacna1h* in estrus females significantly reduced lordosis quotient in response to male mounts (n = 5-7 mice/group; Two-way ANOVA, Bonferroni multiple comparisons test, ** p < 0.01). (F) Estrus cycle length did not significantly differ between Cacna1h^fl/fl^::Cre and Cacna1h^fl/fl^::mCherry females. Pie charts show phase durations over 8 days (n = 8/group; Mann-Whitney test, p > 0.05; Proestrus, purple; Estrus, orange; Metestrus, blue; Diestrus, green). (G) mPFC *Cacna1h* deletion in male mice increased sniffing time towards females (n = 7-10 mice/group; Two-way ANOVA, Bonferroni multiple comparisons test, *** p < 0.001). (H-I) mPFC *Cacna1h* ablation in males significantly increased preference for females in the three-chamber test (H) and social interaction with females during freely moving encounters (I) (n = 5-8 mice/group; Two-way ANOVA, Bonferroni multiple comparisons test, ** p < 0.01). (J-N) Conditional knockout of *Cacna1h* in males increased mounting and intromission behaviors toward females (n = 8-9 mice/group; Two-way ANOVA, Bonferroni multiple comparisons test, n.s., p > 0.05, * p < 0.05).

In contrast, deletion of *Cacna1h* in the male mPFC enhanced sociosexual interest, as evidenced by increased duration of female-directed sniffing (Figures 7G and 7I) and elevated preference for female counterparts (Figure 7H). Furthermore, mPFC *Cacna1h* deletion led to heightened sexual behaviors in males, including increased mounting and intromission behaviors toward females (Figures 7K-7N). Notably, mPFC *Cacna1h* deletion did not affect the time spent in same-sex social interaction or object investigation in either sex (Figures S16C-S16D). Moreover, it did not alter locomotor activity, as assessed by total distance moved in the open field test, or baseline anxiety, as measured by open arm entries and duration in the elevated plus maze test and center duration in the open field test in both sexes (Figure S17). These results suggest that *Cacna1h* gene specifically regulates opposite-sex sociosexual behaviors, without influencing sociability, locomotor activity, or baseline anxiety.

Taken together, these findings demonstrate that dynamic *Cacna1h* expression in the mPFC provides the molecular basis for the estrus-dependent and sex-specific regulation of sociosexual interests and sexual behaviors by mPFC^Cacna1h+^ neurons.

### mPFC^Cacna1h+^ neurons exert sexually dimorphic top-down control of sociosexual behavior via AHNc-descending pathways

The anterior hypothalamic nucleus (AHN) is a key sexually dimorphic region across several species^43–49^. Our previous findings revealed that the estrous cycle modulates AHNc neuronal responses to sociosexual behavior and synaptic plasticity of AHNc-projecting mPFC neurons (Figures 1F-1L). Furthermore, *Cacna1h* exhibits high enrichment in mPFC neurons projecting to the AHNc (Figures 1M). Based on these observations, we hypothesized that AHNc-descending pathways mediate the sex-specific function of mPFC^Cacna1h+^ neurons in regulating sociosexual behavior. To test this hypothesis, we employed a circuit-specific chemogenetic approach to selectively inhibit the DREADD-expressing axonal terminals of mPFC^Cacna1h+^ neurons in the downstream AHNc region (Figures S18A-S18F). Silencing the projections of mPFC^Cacna1h+^ neurons in the AHNc via cannula-guided CNO infusion specifically reduced the social sniffing time toward males in estrus females but not in non-estrus females (Figure S18G). Intriguingly, inhibition of the same mPFC^Cacna1h+^ neurons-AHN pathway elicited reversible effects on sociosexual behavior in males, promoting their investigation of female conspecifics (Figure S18H). Collectively, these findings demonstrate that mPFC^Cacna1h+^ neurons mediate estrous-dependent and sexually dimorphic regulation of sociosexual behavior through descending projections to the AHN.

## DISCUSSION

We identified a population of primary estrous-sensitive neurons in the mPFC that express *Cacna1h* gene. The Cacna1h-defined L5_PT neurons form a circuit with OxtrINs and AHN, and integrate internal hormonal states with external social cues to guide sexually dimorphic sociosexual behaviors. The mPFC^Cacna1h+^ neurons encode opposite-sex investigation, showing enhanced activity in estrus females but predominant suppression in males. These neurons promote estrus-specific opposite-sex interactions and mating behaviors in females, while suppress these behaviors in males through their projections to the AHN. Notably, mPFC^Cacna1h+^ neurons exhibit biased representation of opposite-sex cues in estrus females and males, and mix-represented self-estrous state with target-sex information in females. Mechanistically, ovarian hormones upregulate *Cacna1h*-encoded T-type calcium channels in mPFC^Cacna1h+^ neurons during estrus, enabling enhanced rebound excitation following inhibition from presynaptic oxytocin receptor-expressing interneurons. This modulation drives estrus-specific activity changes and mediates sexually dimorphic effects of mPFC^Cacna1h+^ neurons on sociosexual behaviors. Our study provides direct demonstration of how the mPFC integrates internal physiological states and external social cues to orchestrate adaptive innate social behaviors, unveiling a novel top-down neural mechanism that regulates estrus-dependent and sexually dimorphic sociosexual behaviors. This discovery advances our understanding of the neural basis underlying the complex interplay between internal and external factors in shaping context-specific social behaviors (Figure S19).

### Estrus-sensitive mPFC neurons integrate internal states and external social cues to guide adaptive sociosexual behaviors

The mPFC is well-positioned to potentially integrate social cues and physiological states and guide adaptive social preferences, given its established roles in executive function^60,61^, information integration^14^, behavioral flexibility^15–17^, and social behavior regulation^18–22^. Despite the mPFC’s involvement in estrus-dependent male investigation^3^, the cortical mechanisms governing its dynamic control of male-directed preferences in females over the reproductive cycle remain largely unexplored. Notably, while the mPFC guides male preference for females through biased encoding of female cues, non-specific excitatory mPFC neurons in females exhibit similar proportions of male- and female-preferring cells^19^. This intriguing observation highlights the necessity for cell-type-specific investigations and underscores the existence of sex differences in the neural mechanisms underlying sex encoding and sociosexual preference.

Our study addresses this critical gap by identifying a distinct population of estrous-sensitive mPFC^Cacna1h+^ neurons that exhibit sexually dimorphic encoding of opposite-sex cues and regulate estrus-specific sociosexual behaviors in females. We demonstrate that mPFC^Cacna1h+^ neurons periodically update the representation of social cues based on internal estrous states, exhibiting a marked shift toward excitation in their populational responses to male targets during estrus. Furthermore, these neurons show improved decoding accuracy for the sex of conspecifics during estrus compared to diestrus (Figures 4 and 5). This estrous-dependent enhancement in social cue representation may serve as a neural mechanism for prioritizing sociosexual interactions with potential mates when females are most receptive, ensuring the optimization of reproductive success.

Importantly, we uncover the critical role of mPFC^Cacna1h+^ neurons in encoding internal estrous states in females. Distinct subpopulations of these neurons exhibit unique activity patterns across the estrous cycle, with all subpopulations robustly encoding internal estrous state information and some also encoding the sex of conspecifics (Figures 5). These findings reveal the heterogeneous and specialized functions of mPFC^Cacna1h+^ neurons in the mixed representation of social and physiological information, providing direct evidence to support the hypothesis that mPFC integrates internal states and external social cues to guide appropriate social decisions^62^.

Our study identifies Cacna1h+ neurons as a key subpopulation mediating the interplay between hormonal states and sex-specific processing of social information, providing novel insights into the top-down regulation of adaptive innate social behaviors. This discovery advances our understanding of the neural basis of innate social behaviors and highlights the mPFC as a central hub for integrating diverse streams of information to flexibly guide context-specific, sexually dimorphic sociosexual interactions.

### Cellular and molecular mechanisms driving the dynamic encoding of estrous states and target sex in mPFC^Cacna1h+^ neurons

Our findings demonstrate that mPFC^Cacna1h+^ neurons encode internal estrus state and dynamically update the representation of target sex across the estrous cycle. However, the cellular and molecular mechanisms underlying these dynamic encoding processes in sociosexual behaviors remain unknown. Notably, mPFC ovarian hormone receptor expression levels do not correlate with changes in sex-hormone regulated IEG networks across mPFC subpopulations (Figure 1D-1E). Single-cell RNA sequencing identified mPFC^Cacna1h+^ neurons as the most sensitive to ovarian hormone changes among mPFC neural types, as evidenced by the most dramatic changes in sex-hormone regulated IEG networks across estrous states (Figure 1E). Our results further indicate that Cav3.2 T-type calcium channels, encoded by *Cacna1h*, are the primary driving force in generating estrus-specific activity patterns in mPFC^Cacna1h+^ neurons, thus linking internal physiological state switches to adaptive sociosexual behaviors.

We showed that OxtrINs monosynaptically connect with AHN-projecting neurons enriched with *Cacna1h* (Figure S4C). During estrus, ovarian hormones upregulate *Cacna1h* via the genomic pathway, increasing the expression of Cav3.2 T-type channels in mPFC^Cacna1h+^ neurons (Figures 6I-6J). Social sniffing^63^ or PVN stimulation triggers oxytocin release in the mPFC, activating OxtrINs and hyperpolarizing their postsynaptic mPFC^Cacna1h+^ neurons (Figure S14A-S14B). This hyperpolarization allows more Cav3.2 channels to recover from voltage-dependent inactivation, resulting in heightened rebound activation of mPFC^Cacna1h+^ neurons specifically during estrus (Figures 6K-6L, and S14D). Consequently, when estrus females investigate males, the activity of mPFC^Cacna1h+^ neurons shifts from inhibition to excitation, updating the representation of male cues in the mPFC (Figure 4). Furthermore, the varied T-type currents observed in layer 5 mPFC pyramidal neurons may account for the heterogeneous activity patterns of mPFC^Cacna1h+^ subpopulations between estrus and diestrus (Figures 5K). Ultimately, the upregulation of Cav3.2 channels by ovarian hormones, combined with the actions of oxytocin on OxtrINs, is the critical mechanism driving estrus-specific activity patterns in mPFC^Cacna1h+^ neurons, enabling these neurons to dynamically encode social cues and internal estrous states, ultimately promoting sociosexual behaviors exclusively during estrus.

Despite the impact of sex hormones on a variety of genes, Cav3.2 channels act at an early stage to membrane hyperpolarization during social approach due to their low-voltage activated properties^53,64^. Cav3.2 activation initiates a cascade of events: depolarization of the membrane potential, opening high-voltage gated channels, and change neuronal excitability and synaptic efficacy. The Cav3.2-mediated mechanism precisely modulates mPFC^Cacna1h+^ neural activity based on hormonal fluctuations, enabling adaptive female preference toward males tailored to the specific demands of the estrous cycle. Further investigation into T-type calcium channels in females may uncover additional estrous-dependent and sex-specific behavioral adaptations to internal states, with implications for studying disorders related to sexual behaviors.

### Sexually dimorphic top-down roles of the mPFC in sexual behavior

Sexual behavior exhibits notable sexual dimorphism in both normal sexual behaviors and disorders^7,65–67^. Hyposexuality affects women approximately twice the rate of men^68–71^, while hypersexuality is more than two times prevalent in men^72–77^. Despite these observed differences, the neural mechanisms underlying this sexual dimorphism remain unexplored. Neuroimaging studies show increased prefrontal cortex (PFC) activity during visually evoked sexual arousal^78–83^. In males, enhanced PFC activity correlates with sexual inhibition^73,84^, while PFC dysfunction is linked to hypersexual behaviors, which are more commonly observed in men^75,85–87^. Animal studies support the mPFC’s role in modulating sexual activity, as lesions in this region decrease sexual behaviors in rats^88,89^. In contrast, women’s sexual interest fluctuates with estrogen levels throughout the menstrual cycle. During ovulation, characterized by peak estrogen and progesterone levels, women display increased mPFC activity in response to erotic stimulation compared to other cycle phases^90,91^. These differences suggest a sexually dimorphic role for the mPFC in regulating sexual behavior but the underlying neural mechanisms remain largely unexplored.

Our study uncovers the sexually dimorphic and estrus-dependent roles of mPFC^Cacna1h+^ neurons in regulating opposite-sex sexual behaviors. These neurons exhibit pronounced sex differences in gene expression (Figure S1F), neural response to opposite-sex social cues (Figures 4D-4M), and biased representation of opposite-sex information between estrus females and males (Figures 4H-4P). Manipulating mPFC^Cacna1h+^ neurons or *Cacna1h* expression in the mPFC demonstrates their sexually dimorphic role in orchestrating sexual behaviors, promote sexual behavior in females while primarily inhibiting it in males (Figures 2 and 3). Opposite-sex cues excite these neurons in estrus females but predominantly suppress them in males (Figures 4D-4L), providing a neural substrate for the differential regulation of sociosexual behaviors between sexes and highlighting the dynamic processing of social stimuli across sexes and physiological states. Our findings reveal a neural mechanism of sexually dimorphic top-down regulation of sexual behaviors by mPFC^Cacna1h+^ neurons, implying that dysfunction of these neurons and the *Cacna1h* gene may contribute to hypersexuality in males and hyposexuality in females. This suggest that T-type calcium channels could be potential targets for developing sex-specific therapies for sexual dysfunctions.

Our study further demonstrates that mPFC^Cacna1h+^ neurons exert sexually dimorphic top-down control of sociosexual preference via AHN-descending pathways (Figure S18). The AHN has strong bidirectional connectivity with VMHvl^Esr1+^ neurons^92^, which play a critical role in promoting female sexual behaviors during estrus^4,29,30^. These connections may facilitate the top-down regulation of the transition from sociosexual interaction to sexual receptivity in females during ovulation. However, the specific role of the AHN-VMHvl^Esr1+^ circuit in estrus-linked reproductive behavior warrants further investigation.

## ACKNOWLEDGMENTS

We thank Cuidong Wang at Rockefeller University for technical assistance with TRAP experiments; the Rockefeller University Genomics Resource Center for the help with TRAPseq data analysis; Dr. Diane Lipscombe at Brown University for kindly providing the Cacna1h-IRES-cre knock-in mouse line. This work was supported by STI2030-Major Projects (2021ZD0203000 (2021ZD0203002)), National Science Foundation of China (32171016) and Leon Levy Neuroscience Fellowship (to K.L.).

## AUTHOR CONTRIBUTIONS

Y.W., X.S., F.Z., K.L. and X.C performed the behavioral manipulations, electrophysiology, fiber photometry recordings and analyzed related data; K.L. performed the TRAPseq experiments and X.S. analyzed the TRAPseq data; X.S. and F.Z. conducted the single-cell RNAseq experiment. X.S., Y.Y., and D.M. analyzed single-cell RNAseq data; F.Z. conducted the RT-PCR, OVX and ChIP experiments; L.W. and Y.W. performed Miniscope calcium imaging; Y.Z., J.M., B.Miao., G.Q., and X.J. analyzed the data of Miniscope imaging; G.Q., N.Y. and P.R helped with the experiments and data analysis of fiber photometry recording; Y.W. and K.L. prepared the figures; K.L. wrote the manuscripts with inputs from other co-authors; K.L., N.T. and I.I-T designed and supervised the study.

## STAR METHODS

### KEY RESOURCES TABLE

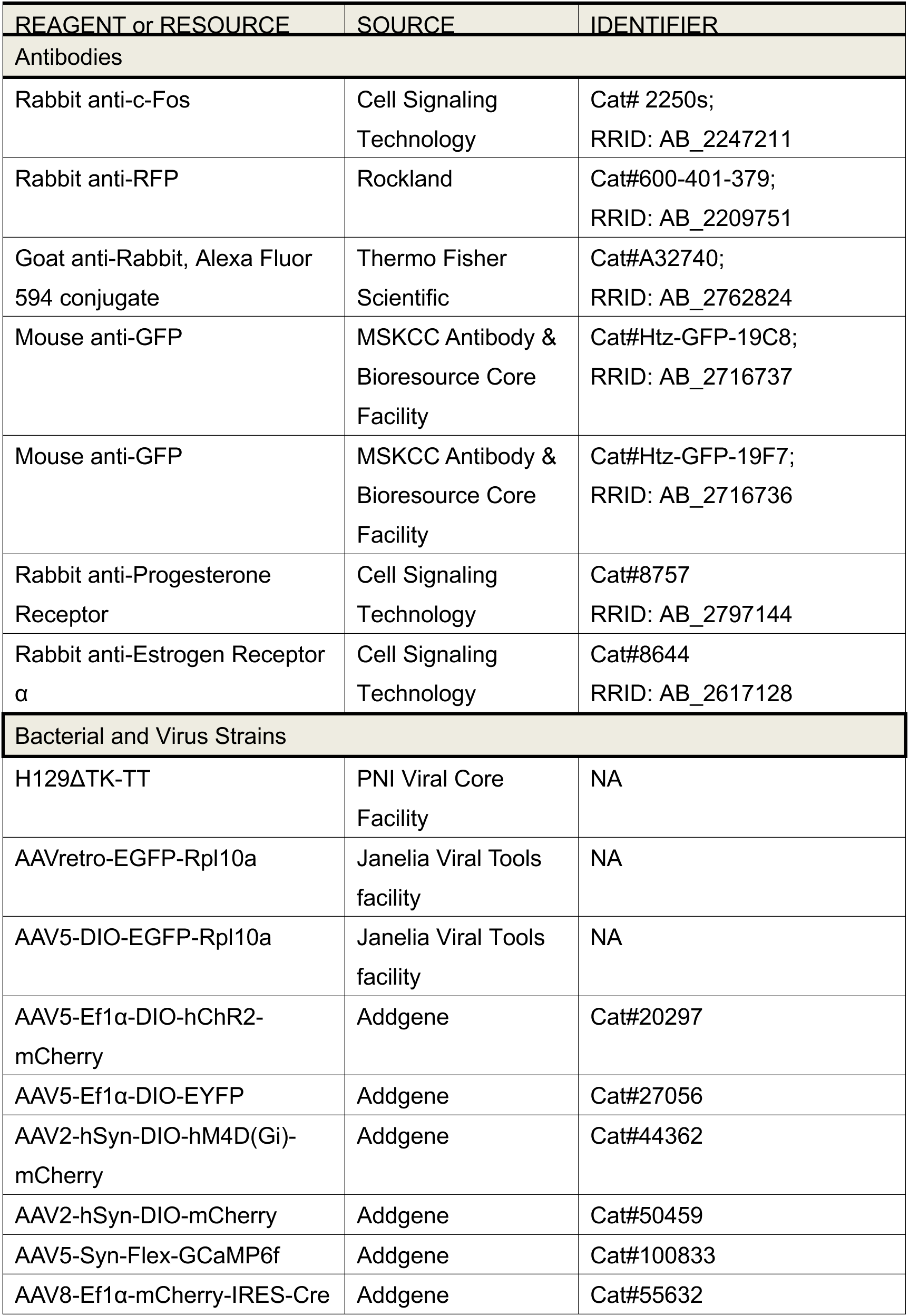
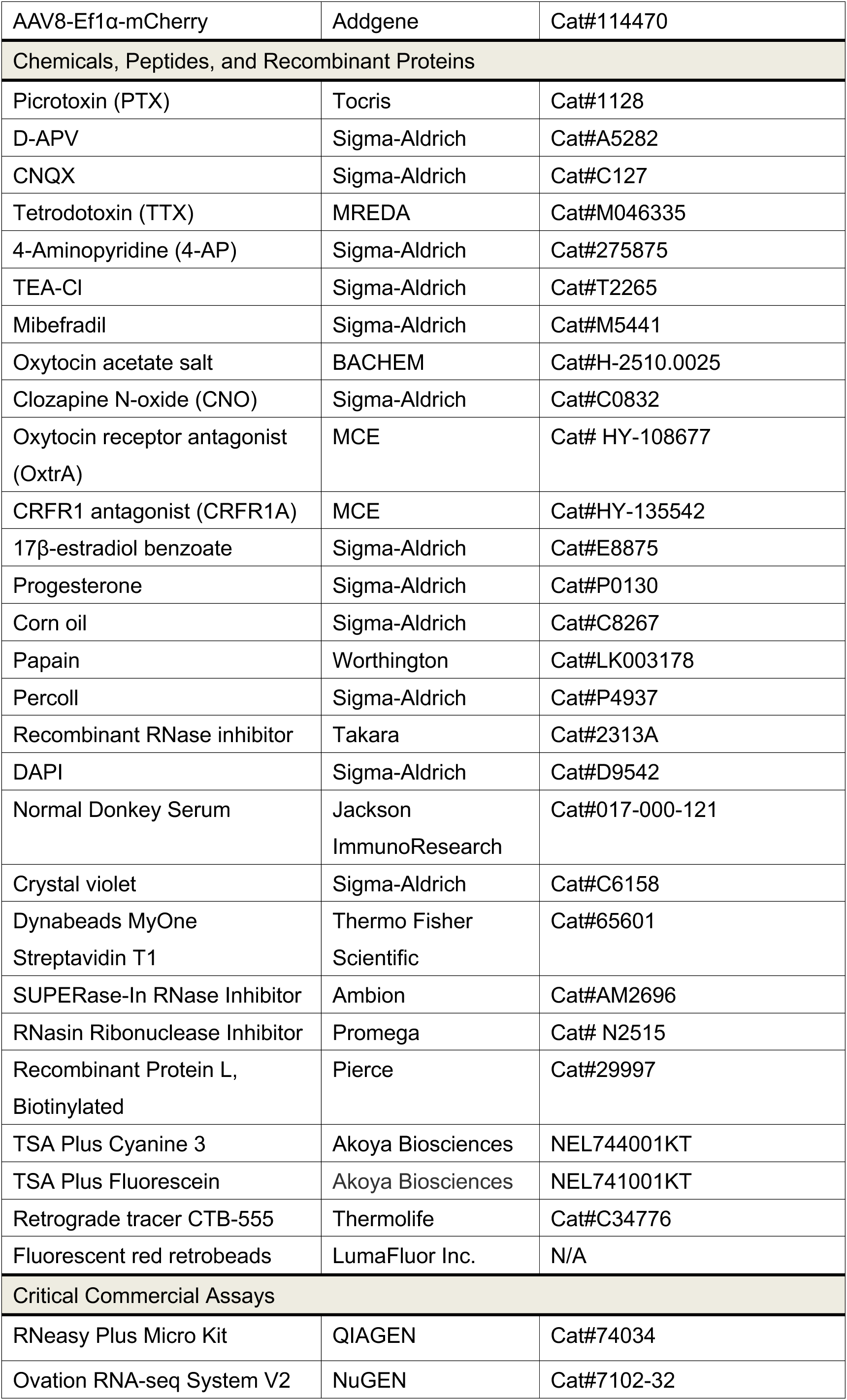
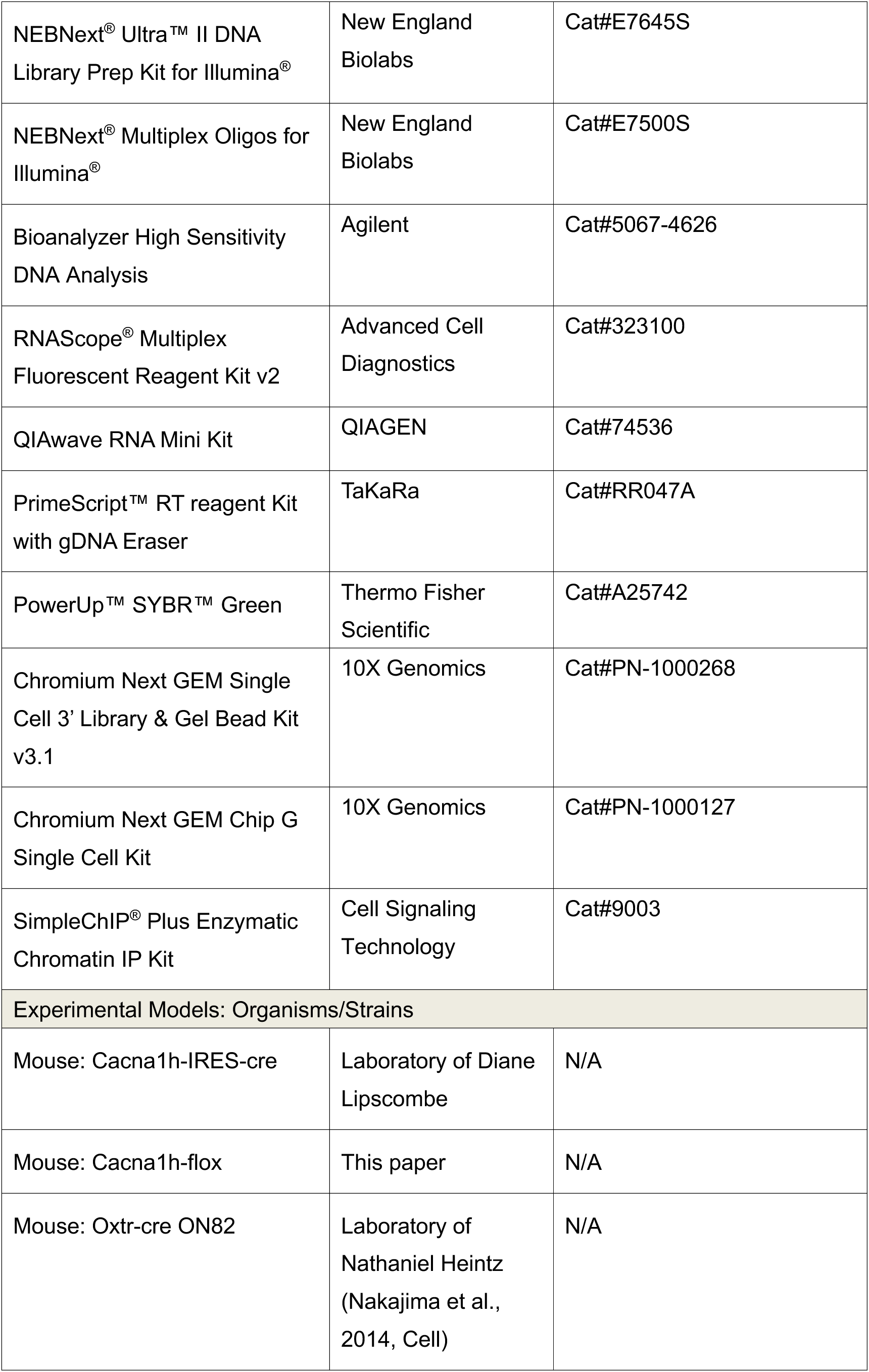
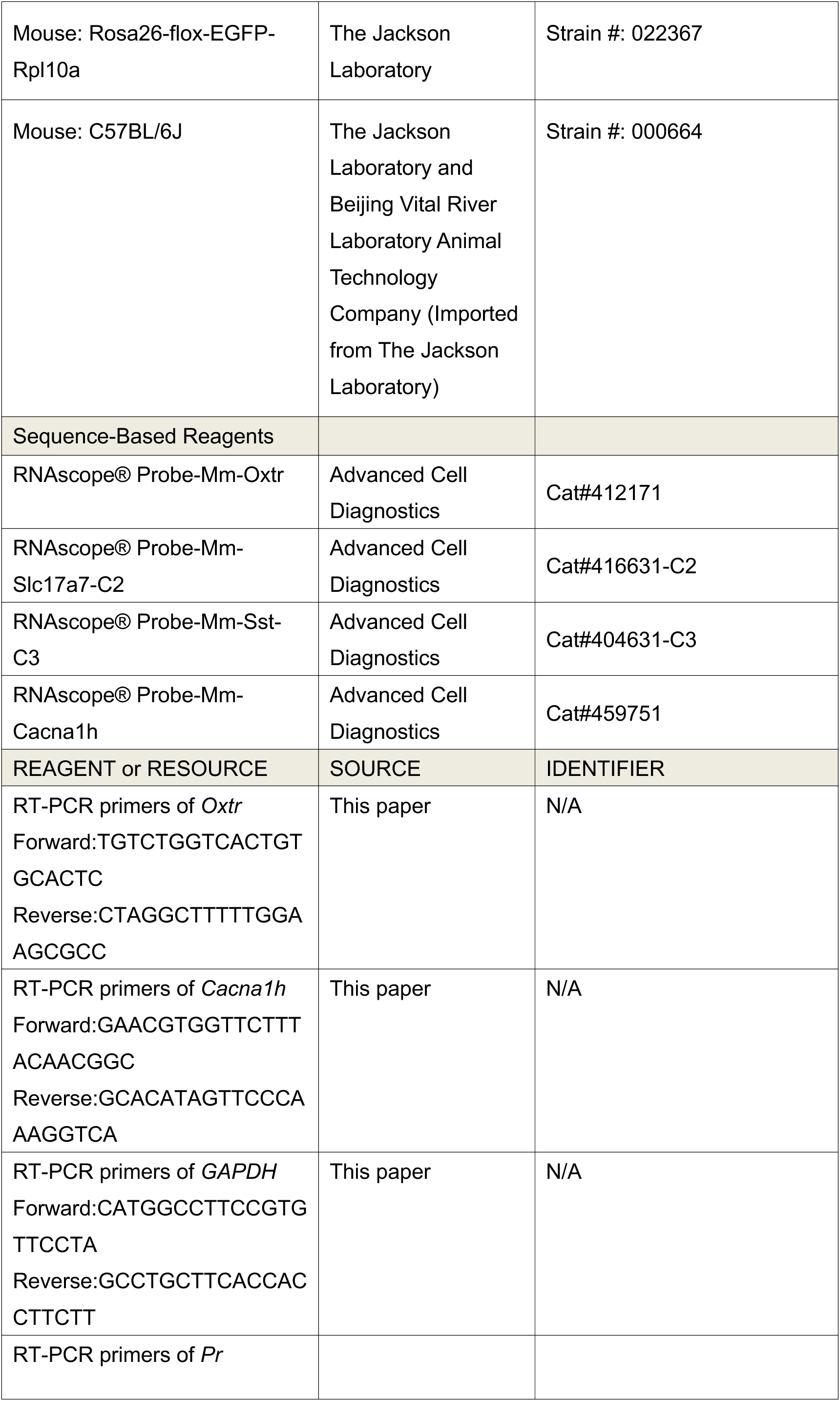
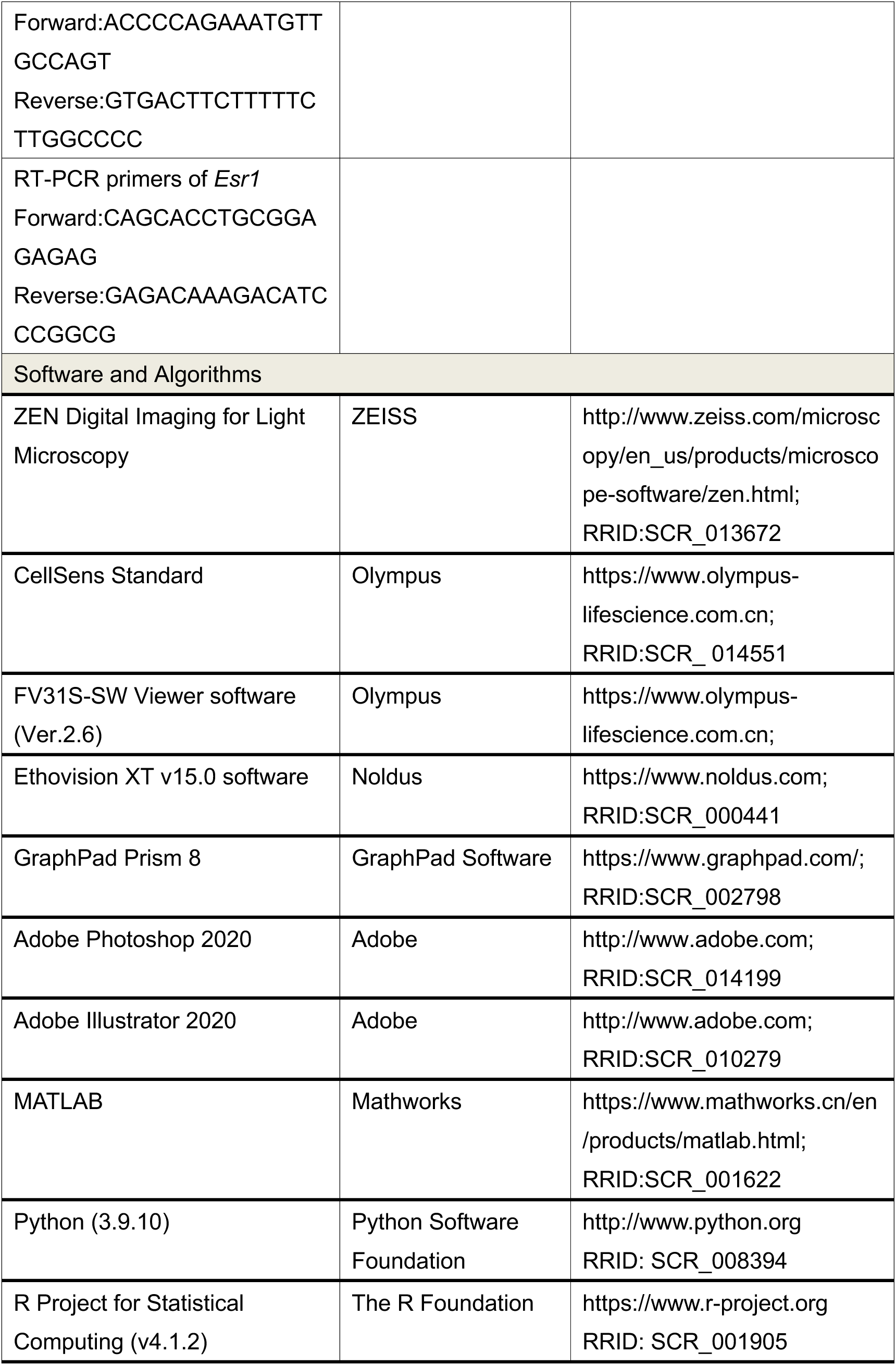

### RESOURCE AVAILABILITY

#### Lead contact

Further information and requests for reagents should be directed to and will be fulfilled by the lead contact, Kun Li.

#### Data and code availability

- The scRNAseq and TRAPseq data has been uploaded at Gene Expression Omnibus.
- All MATLAB, Python and R codes used in this paper are available from the lead contact upon request.

### EXPERIMENTAL MODEL AND SUBJECT DETAILS

#### Mice

All experimental procedures in this study were performed in accordance with the Tsinghua or Rockefeller University guidelines and animal protocols, which were approved by Tsinghua University or The Rockefeller University Institutional Animal Care and Use Committee (IACUC). Wild-type (C57BL/6J, 8 weeks) mice were purchased from The Jackson Laboratory (Strain #: 000664) and Beijing Vital River Laboratory Animal Technology Company (Imported from The Jackson Laboratory, Strain #: 000664). Oxtr-cre ON82 mouse line was generated at the Rockefeller University^3^. Cacna1h-IRES-cre knock-in mouse line was kindly provided by Dr. Diane Lipscombe at Brown University. The conditional knockout mouse line of *Cacna1h* (Cacna1h^flox/flox^) mouse was generated by Cyagen Biosciences Company. Rosa26-flox-EGFP-Rpl10a line was purchased from Jackson Laboratories (Strain #: 022367). Cacna1h-IRES-Cre mouse was crossed with Rosa-flox-EGFP-Rpl10a line to generate the Cacna1h-EGFP-Rpl10a (Cacna1h-TRAP) line. All the mice were same-sex group housed (4-5 mice per cage) in a 12-h light-dark cycle with unlimited access to water and food.

### METHOD DETAILS

#### Identification of the female estrous cycle

The estrous cycle of female mouse is divided into four stages: proestrus, estrus, metestrus, and diestrus, which were determined by cytological evaluation of vaginal smears according to the published literature^93,94^. To evaluate the mouse estrous cycle, the tail of female mouse was gently and firmly grasped to raise the rear body and the vaginal opening of vaginal canal was rinsed by sterile saline with a 200 μl pipette tip. The pipette tip with approximately 100 μl of residual liquid was then placed close to the opening of vaginal canal. The liquid was aspirated slowly and released gently five to eight times to collect the vaginal lavage solution. Even though mice are less susceptible to vaginal stimulation-induced pseudopregnancy, the pipette tip should never be inserting inserted into the vaginal canal to avoid the effect of vaginal stimulation, and all operations were performed with extreme care and delicacy. The vaginal smears were dried on slides and stained with 0.1% crystal violet solution to determine the cytology present. The criterion for the proestrus stage was the presence of round nucleated epithelial cells with relatively uniform appearance and size. The estrus stage is defined by the presence of exclusively anucleated keratinized epithelial cells. At the early stage of estrus phase, the anucleated keratinized epithelial cells are smaller and are typically arranged in loose clusters or sheets. As estrus phase progresses, the cell numbers of keratinized epithelial cells increase, and the cells appear larger and more evenly dispersed and can be seen in stacks or layers. The metestrus stage was characterized by a combination of anucleated keratinized epithelial cells and neutrophils. The diestrus stage was the presence of a large number of neutrophils and few numbers of nucleated epithelial cells.

#### Behavioral testing

##### Sociosexual preference test

The sociosexual preference test of male and female mice were performed in a three-chamber social interaction box as previously described ^3,41^. The apparatus is a rectangular box with three chambers. Subject mice are allowed free access to each chamber through the open middle section on the dividing walls. Prior to the behavioral test, both the subject mice and age-matched, opposite-sexed target mice were acclimated to the testing environment for at least one hour. The sociosexual preference test was comprised of two sessions: first habituation session and second test session. During the habituation session, the subject mouse was put into the middle chamber and able to freely explore all three chambers of the apparatus for 10 min. During the second test session, a stranger target mouse of opposite sex and a novel lego were placed into the wire cups located in left or right side of the chamber randomly. The subject mouse was allowed to freely investigate the three chambers for another 10 min. We defined the ‘Sniff duration’ as the cumulative amount of time that the software detected nose-point of the subject mouse toward the wire cup in the social interaction zone, which is a circular zone surrounding the wire cup containing the mouse or lego (Ethovision XT v15.0 software, Noldus). The three-chamber apparatus was sequentially cleaned using 84 disinfectant (5%) and purified water after the behavioral test of each mouse. The vaginal smears of female subjects were collected following their behavioral tests.

For sociosexual preference test combined with cell type-specific chemogenetic manipulation, the CNO was completely dissolved in DMSO and then diluted to a final concentration of 1 mg/ml in saline. Thirty minutes before the habituation session, the CNO was administrated intraperitoneally at a dose of 1 mg/kg 30 minutes before the habituation session. For pathway-specific chemogenetic manipulation, the DMSO dissolved CNO was diluted to a final concentration of 3 μM in artificial cerebrospinal fluid (ACSF). A dose of 500μl CNO was infused into the AHNc through the internal cannula 30 minutes prior to behavioral tests.

##### Sex preference test

To test the mouse sex preference between male and female target mice, the sex preference test was performed in a three-chamber social interaction box as previously described^29^. Prior to the behavioral test, subject mice were single housed for at least one week, and both the subject and target animals were habituated to the chamber for 30 min per day for at least two days. The sex preference test was comprised of two sessions: first habituation session and second test session. During the habituation session, the subject mouse was put into the middle chamber and able to freely explore all three chambers of the apparatus for 10 min. During the second test session, a sexually experienced strange male mouse and a strange female mouse were placed into the wire cups located in left or right side of the chamber randomly. The subject mouse was allowed to freely investigate the three chambers for another 10 min. The sniff duration for male or female target mice were manually scored by Ethovision XT v15.0 software (Noldus, Netherlands). The sex preference index (female) = sniffing male - sniffing female/sniffing male + sniffing female. The sex preference index (male) = sniffing female - sniffing male/sniffing female + sniffing male.

##### Sexual behavior test

All sexual behavior experiments were performed during the circadian dark phase. Prior to the behavioral test, subject mice were single housed for at least one week. And on the day of testing, both the subject mice and age-matched, opposite-sexed target mice were acclimated to the dark environment illuminated only with infrared lamps for at least one hour.

To measure the sexual behavior of a male mouse, a hormonal-primed and high-receptive female mouse was intruded into the male home-cage, male mouse was allowed to interact with female for up to 30 min or until male ejaculation. The mounting, intromission and ejaculation of males were manually scored by Ethovision XT v15.0 software (Noldus, Netherlands).

To measure the sexual receptiveness of an estrus or diestrus female mouse, firstly a single-housed and sexually experienced male mice were primed by another receptive female 30 min before the testing, and then the subject estrus or diestrus female mouse was intruded into the male home-cage. Female mouse was allowed to interact with male for up to 30 min or until male ejaculation. The lordosis quotient of females was calculated as the ratio between lordosis events and male mounts with intromission. The rejection percent of females was calculated as the ratio between female quick escape events and male mounts or mount attempts.

##### Freely moving social interaction test

Prior to the behavioral test, subject mice were single housed for at least one week, and both the subject mice and age-matched, opposite-sexed target mice were habituated to the cylindric testing arena (35 cm diameter × 30.5 cm height) for 30 min per day for at least two days. On the day of testing, a subject mouse was entered into the testing arena and habituated for 5-10 min to minimize anxiety and stress. Then an opposite-sexed target mouse was intruded into the arena and their social interaction was recorded for up to 10 min. The social behaviors of subject mouse including sniffing or avoidance were manually scored by Ethovision XT v15.0 software (Noldus, Netherlands).

##### Elevated plus maze

The apparatus comprises a central platform (5 × 5 cm), two open arms (30 × 5 cm), and two enclosed arms (30 × 5 × 15 cm). The maze was placed 40 cm above the floor and the light intensity on the maze was measured by light meters and adjusted to 110 lux at the start of each experimental day. The test mouse was placed in the central platform facing an open arm and was allowed to move freely in the maze for 10 min. The distance moved and the time spent in the open arms were automatically analyzed by Ethovision XT v15.0 software (Noldus, Netherlands).

##### Open field test

Mice were individually introduced into the central zone of the open field apparatus (40 × 40 × 40 cm). The mice were allowed to freely explore their surroundings. Parameters including total distanced traveled and the amount of time spent in the central area of the open field were recorded and automatically analyzed for 5 min by Ethovision v15.0 software (Noldus, Netherlands).

#### Stereotaxic surgeries

Mice were maintained anesthesia and performed surgeries on a stereotactic frame (RWD, China). Viruses were injected into either the mPFC (AP, 2.00 and 1.80 mm from the bregma; ML, ±0.35 mm from the midline; DV, -1.75 mm from the dura) , the AHNc (AP, -0.60 mm from the bregma; ML, ±0.55 mm from the midline; DV, -4.95 mm from the dura) or the PVN (AP, -0.59 mm from the bregma; ML, +1.10 mm from the midline; DV, -4.80 mm from the dura) with calibrated glass pipettes connected to an infusion pump (WPI, USA) at a rate of 20 nl min^−1^. After the surgery, mice were placed on a heating pad for recovery and were allowed to stay in home cage for another 7 weeks before behavioral assays. For chemogenetic manipulation, the cannulae were implanted above the AHNc (AP, -0.60 mm; ML, ±0.55 mm; DV, -4.25 mm; 250 μm inner diameters) and attached to the skull with adhesive dental cement. For optogenetic or fiber photometry experiment, the optic fiber was implanted in the mPFC (0.2 mm dorsal to the virus injection site, 400 μm core, NA 0.48) or the PVN (0.2 mm dorsal to the virus injection site, 200 μm core, NA 0.48). For mini-endoscopic imaging, two weeks after virus injection, the cortex above the injection sites was removed with an aspirator and the Graded-Index (GRIN) lens (diameter 1.0 mm, length 4.38 mm, catalog no. NEM-050-06-08-520-S-1.0p) was implanted (AP, 2.00 mm; ML, -0.40 mm; DV, -1.55 mm). The GRIN lens was secured by Krazy Glue and dental cement. The surface of the GRIN lens was protected by Kwik-Sil.

H129ΔTK-TT virus (PNI Viral Core Facility, 100 nl for each site into the mPFC), AAV5-Ef1α-DIO-hChR2-mCherry-WPRE (Addgene, 20297-AAV5, 5.25 × 10^12^ vg ml^−1^, 300 nl for each site into the mPFC), AAVretro-EGFP-Rpl10a (Janelia Viral Tools facility, 500 nl for each site into the AHNc), AAV5-Ef1α-DIO-EYFP (Addgene, 27056-AAV5,1.0 × 10^12^ vg ml^−1^, 300 nl for each site into the mPFC), AAV2-hSyn-DIO-hM4D(Gi)-mCherry (Addgene, 44362-AAV2, 1.1 × 10^12^ vg ml^−1^, 200 nl for each site into the mPFC), AAV2-hSyn-hM3D(Gq)-mCherry (Addgene, 44361-AAV2, 1.8 × 10^13^ vg ml^−1^, 1:5 diluted, 200 nl for each site into the mPFC), AAV2-hSyn-DIO-mCherry (Addgene, 50459-AAV2, 1.8 × 10^12^ vg ml^−1^, 200 nl for each site into the mPFC), AAV1-Syn-Flex-GCaMP6f-WPRE-SV40 (Addgene, 100833-AAV1, 2.1 × 10^13^ vg ml^−1^, 500 nl unilaterally infused into the mPFC), AAV8-Ef1α-mCherry-IRES-Cre (Addgene, 55632-AAV8, 2.1 × 10^13^ vg ml^−1^, 100 nl for each site into the mPFC), AAV8-Ef1α-mCherry (Addgene, 114470-AAV8, 2.5 × 10^13^ vg ml^−1^, 100 nl for each site into the mPFC), AAV5-DIO-EGFP-Rpl10a (Janelia Viral Tools facility, 300 nl for each site into the mPFC), AAV2-hSyn-DIO-mCherry (Addgene, 50459-AAV2, 1.8 × 10^13^ vg ml^−1^, 200 nl for each site into the mPFC), AAV2/9-hSyn-DIO-jGCaMP7b-WPRE-pA (Taitool, S0592-9-H5, 1.73x10^13 vg/ml, 1:5 diluted, 250 nl for each site into the mPFC), AAV2/9-hSyn-ChrimsonR-tdTomato-WPRE-pA (Taitool, S0217-9, 2.0x10^12 vg/ml, 200 nl into the PVN), CTB-555 (Thermolife, C34776, 1.0 mg ml^−1^, 150 nl unilaterally delivered into the AHNc), Red retrobeads (LumaFluor, Inc., 1.0 mg ml^−1^, 100nl for each site into the AHNc) were aliquoted and stored at -80°C until use.

#### Brain slice electrophysiological recordings

##### Brain slice preparation

Adult mice were deeply anesthetized with avertin (2.5% w/v, i.p.) and perfused with 20 mL chilled dissection buffer (oxygenated with 95% O_2_, 5% CO_2,_ 110.0 mM Choline chloride, 25.0 mM NaHCO_3_, 1.25 mM NaH_2_PO_4_, 2.5 mM KCl, 0.5 mM CaCl_2_, 7 mM MgCl_2_, 25.0 mM Glucose, 11.6 mM Sodium ascorbate, 3.1 mM Pyruvate acid. After decapitation, coronal mPFC slices were sectioned to a thickness of 300 μm in chilled dissection buffer using a VT1200s vibratome (Leica). The brain slices were incubated in oxygenated ACSF (118 mM NaCl, 2.5 mM KCl, 26 mM NaHCO_3_, 1 mM NaH_2_PO_4,_ 10 mM glucose, 1.3 mM MgCl_2_ and 2.5 mM CaCl_2,_ oxygenated with 95% O_2_ and 5% CO_2_) to recover for 60 minutes at 32°C and then kept at room temperature until transferring to a recording chamber (Warner Instruments) for electrophysiological recordings.

##### Whole-cell patch-clamp recordings

Brain slices were continuously perfused at 2-3 ml/min with the oxygenated ACSF solution. Whole-cell patch-clamp recordings were performed in the mPFC layer 5 pyramidal neurons with or without fluorescence labeling at 30°C using a temperature control system (HCT-10, ALA). The patch pipette resistance with 5-8 MΩ was pulled with a horizontal micropipette puller (P1000, Sutter Instruments) from glass capillaries (World Precision Instruments, Inc.). Signals were amplified using MultiClamp 700B, filtered at 2 kHz, and sampled at 10 kHz using Digital 1550B (Axon Instruments). The series resistance and capacitance were compensated automatically after a stable Gigaseal was formed. Data analysis was performed offline with Clampfit 11.1 (Molecular Devices). For voltage-clamp recording, the patch pipette was filled with internal solution containing (in mM) :115 CsMeSO_3_, 20 CsCl, 10 HEPES, 2.5 MgCl_2_, 4 Na_2_ATP, 0.4 Na_3_GTP, 10 Na-phosphocreatine, and 0.6 EGTA. To measure sEPSCs and sIPSCs, cells were clamped at −70 mV and 10 mV respectively. A total of 30 sweeps were taken at individual potential, and each sweep lasted 10 seconds. Series and membrane resistances were monitored by giving a 5 mV pulse. Recordings in which series resistance exceeded 30 MΩ were excluded. For current-clamp recording, the internal solution contained (in mM): 130 K-gluconate, 5 KCl, 10 HEPES, 2.5 MgCl_2_, 4 Na_2_ATP, 0.4 Na_3_GTP, 10 Na-phosphocreatine, 0.6 EGTA. RMP was measured by reading Vm under I = 0 mode right after breaking into cells. To measure current-induced firing rate of mPFC layer 5 pyramidal cells, sequential currents were injected from -20 pA to 340 pA in a 40 pA step during 500 ms. Oxytocin (100 nM, BACHEM, H-2510.0025) and mibefradil (10 nM, Sigma-Aldrich, M5441) were perfused with ACSF to measure their effects on RMP or current-induced firing rate.

To measure light-evoked postsynaptic currents, 470 nm blue light (25 mW, 5 ms) was delivered to the patched cells in the mPFC brain slice by an optic fiber attached to a laser. Layer 5 pyramidal neurons were clamped at -70 mV for light evoked-EPSCs (eEPSCs) and 10 mV for light evoked-IPSCs (eIPSCs). Picrotoxin (100 μM, Tocris, 1128) was perfused with ACSF to validate the inhibitory postsynaptic currents, CNQX (20 μM, Sigma-Aldrich, C127) and D-APV (50 μM, Sigma-Aldrich, A5282) were used to confirm the excitatory postsynaptic currents. Peak amplitude was determined by subtracting the baseline value. Synaptic latency was calculated as the interval between the start of light stimulation and the current onset time. To access the paired pulse ratio, two brief light pulses with 150 ms intervals for eEPSCs and 350 ms intervals for eIPSCs were delivered to the mPFC slices. For the purpose of isolating monosynaptic responses, eEPSCs and eIPSCs were elicited by a single 5ms-light pulse in the presence of 4-AP (1 mM, Sigma-Aldrich, 275875) and TTX (1 μM, MREDA, M046335).

To isolate T-type calcium currents pharmacologically, 1 μM TTX, 10 mM TEA-Cl (Sigma-Aldrich, T2265), and 1 mM 4-AP were perfused with ACSF in the acute mPFC slices. The cells were clamped at -110 mV. A depolarization step with 300 ms duration was applied from holding potentials of -110 mV to -45 mV. The T-type currents were identified by subtracting traces at two different holding potentials. The amplitude of T-type currents was determined as the peak current in subtracted traces.

To confirm the silencing effect of CNO on cells expressing hM4Di, CNO (3 μM, Sigma-Aldrich, C0832) was perfused with ACSF on mPFC slices infected with AAV-DIO-hM4Di-mCherry viruses. mCherry tagged mPFC cells were recorded for 10 minutes before and after CNO perfusion, and then for additional 5 minutes following the CNO washout.

#### RNA sequencing

##### scRNA-seq and data analysis

The mPFC tissues were extracted using a brain matrix from two estrus female mice, two diestrus female mice, and two male mice. The dissected tissue was then cut into small pieces and transferred to a 15 ml tube containing 20 U/mL pre-activated papain (Worthington, LK003178, 37°C for 30 min) and 0.1 U/uL recombinant RNase inhibitor (Takara, 2313A). Papain digestion was performed at 34℃ for 20 min. Following digestion, the papain solution was replaced with ice-cold, carbogen-bubbled ACSF (pH 7.2-7.4, 125 mM NaCl, 2.5 mM KCl, 25 mM NaHCO3, 1.25 mM NaH2PO4, 25 mM glucose, 1 mM MgCl2, and 1 mM CaCl2, oxygenated with 95% O2 and 5% CO2) containing 1% fetal bovine serum (FBS) and 0.1 U/μL recombinant RNase inhibitor. The tissue pieces were dissociated using gentle pipetting through 1000μL and 200μL tips, to obtain single-cell suspensions. Subsequently, the suspensions were filtered through a 35μm cell strainer (Corning, 352235) to remove debris, and then layered onto 30% Percoll (Sigma, P4937, diluted by ACSF containing 1% FBS and 0.1 U/μL recombinant RNase inhibitor), followed by centrifugation at 700g for 10 minutes at 4 ℃ . After two washes, the cells were resuspended in 1X PBS with 0.02% BSA and 0.1 U/μL recombinant RNase inhibitor. The cell viability and concentration were determined, and 15000 single cells per sample were uploaded onto 10 × Genomics 3′ library chips, using the Single Cell 3′ Library & Gel Bead kit (v.3.1), as per the manufacturer’s instructions.

The resulting libraries were sequenced on an Illumina NovaSeq 6000, and the reads were aligned and quantified using the CellRanger (version 3.0.2)^95^ pipeline on the reference mouse genome (mm10). Downstream analysis was performed using the R package Seurat (version 4.0.2)^96,97^, where genes detected in fewer than 3 cells were removed, and cells with nCount_RNA > 1000 and nCount_RNA < 30000, and nFeature_RNA > 500 and nFeature_RNA < 7000 were retained. Potential low-quality cells were filtered out based on a mitochondrial or ribosomal gene percentage greater than 20%. To correct for batch effects, the expression matrixes across all three conditions (estrus, diestrus, and males) were integrated using FindIntegrationAnchors and IntegrateData functions of Seurat. We then identified 5000 highly variable genes using FindVariableGenes and performed principal component analysis (PCA) with RunPCA. Dimension reduction was performed with t-distributed stochastic neighbor embedding (t-SNE) using RunTSNE. Next, we performed unsupervised cell clustering with FindClusters (resolution = 0.2) and identified cluster markers using FindAllMarkers with the default parameters. We used the highly-expressed marker gene Snap25 to identify the neuron clusters. A total of 5705 high-quality neurons were extracted from total cells, including 827 neurons from estrus, 1901 neurons from diestrus, and 2977 neurons from males. To annotate subclusters of mPFC neurons, clustering analysis was performed with FindClusters by setting the parameter resolution to 0.5. We then identified subcluster markers using well-known marker genes of the whole cortex or mPFC single cell RNA-seq and the dataset from Allen Brain Atlas^34,35,37^. Finally, we used FindMarkers with the default parameters to find differentially expressed genes (DEGs) among different estrus/sex state comparisons (estrus vs diestrus, estrus vs males, diestrus vs males), and defined DEGs lists with a cut-off p-value < 0.05. To elucidate how ovarian hormone fluctuation impact individual cell types, de gene screening was performed by the p-value cutoff of estrus vs diestrus DEGs from each neuronal cell type, and then the gene clustering/ordering was conducted by euclidean distance for heatmap layout using R package ComplexHeatmap (version 2.6.2)^98^. Heat map showed the identified cell-type-specific gene blocks of each neuron cluster. We used DAVID^99^ to perform GO enrichment analysis, and filtered enriched Biological Process terms with GeneCount ≥ 2 and p-value < 0.05. We removed synonymous and unassociated GO terms for clarity.

In order to define the neurons with high IEGs expression, and find the estrous cycle- or sex-specific activated clusters, a set of the IEGs regulated with estrogen or progesterone were selected (*Fos, Jun, Junb, Npas4, Nr4a1, Btg2, Tns1, Ptgs2, Per1, Per2, Mcl1, Dusp6, Irs2, Kdm6b, Ifrd1, Cdkn1a* and *Pim1*)^38–40^, the 90th percentile expression for each IEG of estrus, diestrus or male samples was calculated. The neuron was defined “IEG^high^ cell” if it expressed any of the IEGs above this 90th percentile^100^. ΔIEG^high^ cell percentage among different estrus/sex state comparisons (estrus vs diestrus, estrus vs males, diestrus vs males) was calculated by the difference of the IEG^high^ cell percentage of each comparison.

##### Pathway-specific TRAP-seq and cell type-specific TRAP-seq

For pathway-specific TRAP-seq, retrograde viruses rAAV2-EGFP-Rpl10a (Janelia Viral Tools facility, 500 nl for each site) were injected into the AHNc of wild type mice. The mPFC regions from three male and five female mice with AHNc virus injection were used for independent TRAP replicates. For cell type-specific TRAP-seq, 3 male and 8 female Cacna1h-cre::EGFP-Rpl10a mice of four distinct estrous cycles were used for independent TRAP replicates. The TRAP procedure was performed as described previously^101^. The mPFC cortices were dissected in dissection buffer (1X HBSS, 2.5 mM HEPES-KOH 35 mM Glucose, 4 mM NaHCO_3_, RNase-free water, freshly added 100 μg/mL cycloheximide) using a brain matrix one month after retrograde viral infection. The tissues were immediately homogenized in an ice-cold homogenization buffer (150 mM KCl, 10 mM HEPES pH 7.4, 5 mM MgCl_2_, 0.5 mM dithiothreitol (Sigma-Aldrich, D9779), 100 μg/mL cycloheximide (Inalco, 1758-9310), 20 U/μL SUPERase-In RNase Inhibitor (Ambion, AM2696), 40 U/μL RNasin Ribonuclease Inhibitor (Promega, N2515), and EDTA-free protease inhibitors) in the cold room using a motor-driven Teflon glass homogenizer. Polyribosomes were immunoprecipitated by monoclonal anti-EGFP antibodies (MSKCC Antibody & Bioresource Core Facility. a mixture of clones 19C8 and 19F7) that have been conjugated to Dynabeads MyOne Streptavidin T1 (Thermo Fisher Scientific, 65601), which were pre-incubated with Recombinant Protein L (Pierce, 29997) for 16 hours at 4°C on an end-over-end rotator. The RNeasy Plus Micro Kit (QIAGEN, 74034) was used to extract and purify RNAs from ribosomes according to the manufacturer’s instructions. The RNA quality was evaluated using an Agilent 2100 Bioanalyzer (Agilent Technologies, USA). Then, using a total of 5 ng mRNAs from each IP and INPUT sample, cDNA was synthesized and further amplified using the Oviation RNA-seq System V2 Kit (NuGEN,7102-32). cDNA fragments of 200 bp were end-repaired and ligated with adapters for NextSeq 500 technology using NEBNext Ultra™ II DNA Library Prep Kit (New England Biolabs, E7645L) and NEBNext^®^ Multiplex Oligos (New England Biolabs, E7500S) for Illumina. The cDNA libraries were sequenced on the NextSeq 500 System (Ilumina Inc., San Diego, CA, USA). The quality of libraries was evaluated by using Bioanalyzer High Sensitivity DNA Analysis (Agilent, 5067-4626) on the Aglient 2200 TapeStation system.

Sequencing reads were aligned to the UCSC mm10 reference genome with STAR (version 2.3.0e_r291). Aligned reads were quantified by the ‘htseq-count’ module of ‘HTSeq’ framework (version 0.6.0). Differential gene expression analysis was performed by using DESeq2 (version 1.34.0). Genes were identified as differentially expressed with a false discovery rate (FDR) adjusted p value < 0.05 and log_2_Fold Change > 1.2 or < -1.2. To identify all DEGs (differentially expressed genes) linked with female estrous cycle, we performed pairwise comparisons of four samples collected at different estrus phases (proestrus, estrus, metestrus and diestrus) and extracted genes differentially expressed in at least one pairwise comparison. The DEGs were hierarchically clustered into 5 clusters with distinct expression patterns across the estrous cycle. The expression levels of *Cacna1h*, *Cacna1g* and *Cacna1i* in mPFC^Cacna1h+^ neurons at different estrus phases were evaluated by their normalized counts. We used DAVID (Huang et al., 2009) to perform the GO enrichment analysis, and enriched Biological Process terms were filtered with GeneCount ≥ 2 and p value < 0.05. The synonymous and unassociated GO terms were removed for clarity. Fold Enrichment was calculated by GeneRatio/BgRatio.

#### TF motif analysis

To identify the putative TF binding sites in the promoter of *Cacna1h*, we scanned the promoter region (±2kb) of *Cacna1h* by FIMO (MEME suite v5.4.1) to identify TF motifs occurrences with parameters as follows: P value < 10^-4^, a first-order Markov background model, and position weight matrices (PWMs) from the mouse HOCOMOCO motif database (v11).

#### Chromatin immunoprecipitation (ChIP) assays

ChIP assay was performed using the SimpleChIP® Plus Enzymatic Chromatin IP Kit (Cell Signaling cat # 9003) according to the manufacturer’s protocol. Using the Bioruptor® Pico sonication device (Diagenode) to break nuclear membrane. Equal volumes of chromatin were immunoprecipitated with either antibody against PR (Cell Signaling, cat # 8757T), Esr1 (Cell Signaling cat # 8644T), mouse IgG as a negative control, or mouse H3 as a positive control. Primers 5’-ACCCCAGAAATGTTGCCAGT-3’ and 5’-GTGACTTCTTTTTCTTGGCCCC-3’ were designed to amplify the region of Pr -binding site of *Cacna1h* promoter. Primers 5’-CAGCACCTGCGGAGAGAG-3’ and 5’-GAGACAAAGACATCCCGGCG-3’ were designed to amplify the region of Esr1 -binding site of *Cacna1h* promoter.

#### Calcium imaging

##### Photometry setup and recording

The fiber photometry system used for recording GCaMP6f signals was built as previously described ^102,103^. In brief, GCaMP6f fluorescence (473 nm) and isosbestic autofluorescence signals (405 nm) were excited by the fiber photometry system using 473 nm (Thorlabs M470F3) and 405 nm (Thorlabs M405FP1) LEDs, and collimated into a dichroic mirror holder with a 425 nm long pass filter (Thorlabs DMLP425R). This is further passed into another 495 nm long pass dichroic (Semrock FF495-Di02-25x36) to direct the light from LEDs into the fiber patch chord (Doric lenses, 400 μm core, 0.48 NA) via a 10x/0.5NA Objective lens (Nikon CFI SFluor 10X, Product No. MRF00100). The light intensity at the interface between the fiber tips was adjusted to 10∼50 μW. The emitted GCaMP6f fluorescence is transmitted back from the same cable, redirected by the 495 nm long pass dichroic, to a GFP emission filter (Semrock FF01-520/35-25) and focused onto a high sensitivity sCMOS camera (Prime 95b, Photometrics) for amplification and recording. A custom made JK flip-flop is used to alternate between 405nm and 470nm which takes the trigger input from the sCMOS and alternatively trigger the two LEDs. Bulk activity signals were collected using the PVCAM software, and data were further post-processed and analyzed using custom MATLAB scripts.

To synchronize animal behaviors with GCaMP6f signals, the behavior-relevant TTL inputs were triggered by Ethovision XT software (Noduls, Netherlands). Before each behavioral test, subject mouse was allowed to habituate to the fiber patch cord in their home cage for approximately 10 min. Following that, the GCaMP6f signals (F473 nm) and the autofluorescence signals (F405 nm) were alternatively recorded while the animals were performing sociosexual preference test.

##### Fiber Photometry data processing and analysis

To analyze the recording data, the signal was processed at a frame rate of 7.5 Hz. The calcium independent signal from 405nm fluorescence was scaled to best fit the 470nm signal using the least squared regression. After subtracting this scaled 405 nm fluorescence from the 470nm signal, we get the ΔF/F by dividing this value by the scaled 405 nm signal; Fn(t) = 100 × [F470(t)–F405fit(t)]/F405fit(t). Baseline of each mouse was determined and corrected by MATLAB function “msbackadj” which estimated baseline over 200 frame sliding window and used a spline method to regress varying baseline values to the window’s data points, then adjusted the baseline of the peak signals. This method normalizes GCaMP6f signals (F473 nm) with autofluorescence signals (F405 nm) and controls movement and bleaching artifacts over the course of calcium recording. The sniffing events during sociosexual interaction test were defined as the mouse nose actively smelling target mouse or lego located in the wire cups in the interaction zone. Behavioral events were manually time-stamped on the recorded animal tracking videos using EthoVision program (Noduls).

For the peri-event time histogram (PETH) analysis, the average activity from -5 s to 3 s around behavioral bout onset of each mouse was calculated across all trials. Then mean F_0_ and the standard deviation SD_0_ of baseline signals were calculated from -5 s to -3 s prior to each behavior bout onset. And the ΔF/F Z-Score of a given behavior was constructed by [Fn(t)–F_0_]/SD_0_. The change of mean fluorescence was defined as the averaged ΔF/F Z-Score between 0 s to 3 s minus the averaged ΔF/F Z-Score between -3 s to 0 s. As to the trial-by-trial analyses, ΔF/F Z-score of each trial was processed by the same method but without averaging F_0_ across trials. Post-processing and plotting of data were performed with the custom MATLAB scripts.

##### Miniscope imaging

Mice were allowed to recover for 1 week after the implantation and the amoxicillin (10 mg/kg) was administrated in drinking water. Besides, during the recovery, subcutaneous injection of Meloxicam (10 mg/kg) and dexamethasone (0.2 mg/kg) was provided every day. 2-3 weeks after the implantation of the GRIN lens, mice were anesthetized again and GCaMP expression was checked by customized miniscope V4 system (UCLA miniscope, from OpenEphys). The Miniscope was slowly lowered to the surface of the GRIN lens. If clear GCaMP-expressing neural shapes were in focus, the baseplate was fixed with dental cement. Before the behavioral testing, mice habituated wearing the Miniscope while freely moving at least 3 times within 1 week.

##### Extraction of Single Cell Calcium Signals

The raw calcium fluorescence videos were processed with MiniAn^58^, an open-source package based on CNMF. Most parameters were set as recommended. After obtaining the footprint, we extracted the raw neural signals from the preprocessed videos. The footprints and traces were then manually checked. Cells with abnormal footprints or signals are excluded from analysis. The pipeline based on MATLAB is available at https://github.com/lance11320/cc1h_ms.

##### Single cell response analysis

To identify responsive neurons, we applied the receiver operating characteristic (ROC) analysis. Behaviors at each time point were converted to a binary form respectively. This binary data was applied to the ΔF/F of each neuron to classify the specific behavior at each time point. True positive rate and false positive rate were calculated and used to construct the ROC curve. The area under the ROC curve (auROC) indicated how well the neuron was modulated by specific behavior. Then we applied a circularly permuting of random shift 1000 times to the calcium signals as a null distribution and calculated the auROC. A neuron was identified as an activated neuron if its auROC exceeded the 97.5th percentile of null distribution. A neuron was identified as an inhibited neuron if its auROC was less than 2.5th percentile of null distribution. Any neuron with multiple response was classified as a mixed neuron. These activated neurons, inhibited neurons and mixed neurons composed responsive neurons.

To obtain the average single cell responses, the behavior bouts that lasted less than 4s or had an interval less than 5s compared with the previous bout were excluded from analysis. The activity of each cell from 5s before to 5s after the behavior bouts onset was averaged to obtain the average response of each cell during different behavior bouts. Average response was normalized according to the mean activity Fm and standard deviation s.d. over the 2s baseline window between 5s and 3s before bout onset: z-scored ΔF/F = (F(t)-Fm)/s.d..

##### Classifying sniffing object with support vector machine (SVM)

For time-evolving decoding, we constructed a linear SVM classifier with LibSVM for each 10 frames with randomly selected 15 (in sociosexual preference test) or 25 (in sex preference test) neurons to decode the trial type (sniffing Lego or sniffing mouse). Only sessions with no less than 5 trials (in sociosexual preference test) or 10 trials (in sex preference test) were selected for training the classifier (trials with gap less than 3 seconds and duration less than 1 seconds are excluded from all the analysis). We then randomly selected 5 trials for classifying mice against Lego and 10 trials for classifying male against female in each trial type for each category. The accuracy is reported as mean accuracy of 5-fold cross validation. The decoding process above was repeated 50 times.

For statistics, we constructed a SVM classifier for data from 0-2s activity (60 frames in total and down sampled into 20 frames) after sniffing onset with all the other parameters the same as above.

To determine whether the lower accuracy reported in male or diestrus female mice during sociosexual behavior is due to less neuron recorded or less proportion of neurons contributing to decoding, we constructed a pseudo neuron population for decoding. For each session, we randomly selected 5 trials and aligned the activity of neurons from all the sessions at each sniffing onset. We then randomly selected from 1 to 100 neurons to train a decoder. The decoding was repeated 30 times, with accuracy reported as mean accuracy of leave-one-out-cross-validation (LOOCV).

##### Analysis of population activity in low dimensional space

To determine the contribution of neural data from each trial type to SVM classification in sex preference behavior, we project the neural data onto SVM hyperplane. This method draws parallels with the concept of coding direction, which best separates the response vectors of different trial type as described in previous literatures^104–106^. A linear SVM classifier was trained with all recorded neurons on the 0-2s neural data from each trial. Therefore, for a n-neurons population we obtained an n x 1 SVM weight or hyperplane direction vector (HD) which best separates the response vectors of sniffing male target and female target. Given that the activity vector of each frame is denoted as x, the projected activity of each frame is then formulated as HD^T^.x. The absolute value of the projected data signifies the distance to the classification plane, where a larger distance indicates a more substantial contribution to the classification. Additionally, when interpreting the hyperplane direction as SVM weight, lower projection results may suggest that neurons responsive to this trial type are assigned smaller weights in the classifier, thereby contributing less to the classification. This implies that these neurons may respond to both trial types, but this complexity is difficult to discern by hard thresholding auROC.

To visualize the distribution of dataset, the neural data of each trial is projected onto a 2-dimensional principal component plane. The plane was calculated based on 0-2s activity of each trial and activity of baseline period.

##### Decoding estrus and diestrus state with each behavior

A linear SVM decoder of each mouse is trained to distinguish estrus and diestrus state with neural activity in specific behavior. In each mouse, only neurons that are recorded in both estrus and diestrus state are selected for decoding. In each behavior, each sample in the training and testing set is the one trial neural activity of all neurons and 0-4s after behavior onset. The label of each sample is estrus (0) or diestrus (1). For balancing number of samples (trials) in each class, we randomly select the minimum number of trials in estrus and diestrus 50 times and calculate the mean accuracy (Figure 5F, 5K). For balancing the feature dimension of each behavior, the behavior onset time of each home cage and habituation trial is randomly sampled 100 times because these two behaviors last long time and have no behavior onset. So, the number of trials of home cage and habituation behavior is 100.

##### Decoding sex of social targets with estrus or diestrus neural activity

A linear SVM decoder of each mouse is trained to distinguish sniffing female and sniffing male behavior with neural activity in specific self-estrus state. In each mouse, only neurons that are recorded in both estrus and diestrus state are selected for decoding. In each state, each sample in the training and testing set is 0-4s neural activity in one trial of all neurons relative to behavior onset during sniffing female or male behavior. The label of each sample is sniffing female (0) or sniffing male (1). For balancing number of samples (trials) in each class, we randomly select the minimum number of sniffing female and male trials 50 times and calculate the mean accuracy (Figure 5C).

##### Clustering and find subpopulations

Select neurons that are recorded both in estrus and diestrus state. After selection, 162 neurons of all mouses remain (Figure 5L). To detect subpopulations of mPFC^Cacnah1+^ neurons, we concatenate each neuron’s neural activity during different behaviors in estrus and diestrus condition (Figure 5L) and used K-means algorithms for clustering. The number of clusters is determined by combining Sum of Squared Errors (SSE) and calculating Silhouette Coefficient. SSE is the sum of squares of the distances from each point to its nearest cluster center. When different numbers of clusters are selected, SSE typically decreases as the number of clusters increases, because more clustering means that the points in each cluster are closer to their centers. The Silhouette Coefficient combines the degree of clustering density with the degree of separation and provides a measure for each sample, ranging from -1 to 1. The higher the Silhouette Coefficient is, the better the clustering effect is, which indicates that the sample is more suitable for its own clustering and not suitable for the adjacent clustering. Select the K value for the point where SSE starts to slow down (Elbow Method) and the average Silhouette Coefficient has a local maximum (Figure S9 B,C). Before this point, increasing the number of clusters significantly reduced SSE; after this point, however, the contribution of increasing the number of clusters to reducing SSE became less obvious.

##### Optogenetic activation and fiber photometry recording experiment

AAV2/9-hSyn-DIO-jGCaMP7b-WPRE-pA was injected into the medial prefrontal cortex (mPFC), and AAV2/9-hSyn-ChrimsonR-tdTomato-WPRE-pA was injected into the paraventricular nucleus (PVN) of Cacna1h-cre female mice, followed by optic fiber implantation in the mPFC. The CRFR1 antagonist NBI-27914 (250 μl, 1 μg/μl, 10 mg/kg; Cat#HY-135542) was administered intraperitoneally, and calcium signals of mPFCCacna1h+ neurons were recorded 30 min after drug treatment. Subsequently, a mixture of CNQX (250 μl, 0.125 μg/μl, 1 mg/kg; Cat#A5282) and D-APV (250 μl, 1 μg/μl, 10 mg/kg; Cat#C127) was administered intraperitoneally, and recordings were performed 10 min later. For Oxtr blockade, the OxtrA L-368,899 (250 μl, 1 μg/μl, 10 mg/kg; Cat# HY-108677) was injected intraperitoneally, and recordings were made 20 min later. PVN neurons were optogenetically activated with 589 nm yellow light (2 mW, 3 s square wave) after a 2-min baseline. The calcium signal of mPFCCacna1h+ neurons was collected using a Doric photometry recording system (blue light: 470 nm, 20 μW; purple light: 405 nm, 2 μW). Custom MATLAB code was used to extract and analyze the GCaMP signal surrounding each stimulation. Peak amplitudes of the first signal peak during stimulus onset and the rebound peak after light cessation were quantified.

#### RNAscope

RNAScope was performed following the procedure described in the RNAScope Multiplex Fluorescent Reagent Kit v2 (Advanced Cell Diagnostics, 323100) with minor modifications. In brief, mice were perfused with 4% paraformaldehyde solution (PFA). Mouse brains were post-fixed in 4% PFA overnight and dehydrated in a 30% sucrose solution at 4°C for 36 h. The mPFC region was sliced in 14-μm sections by cryostat and mounted on Fisher brand slides. The sections were air dried at room temperature and then stored at -80°C until use. On the day of staining, the slides were rinsed with PBS after drying for 1 hour at 65°C. The sections were treated with RNAscope® Hydrogen Peroxide solution to eliminate non-specific background signals. Then the sections were recovered in pre-heated Target Retrieval Reagent and digested by RNAscope® Protease III. To detect the target RNAs, commercial RNA-specific probes (ACDbio RNAscope®) were hybridized with the sections. In order to magnify the probe signals, sections were further sequentially incubated in a cascade of signal amplification reagents. TSA Plus Cyanine 3 and TSA Plus Fluorescein (Akoya Biosciences, NEL744001KT and NEL741001KT) were linked to the probes to visualize target genes. To access the percentage of colocalization between target genes and mCherry^+^ signals, the slides were blocked by 3% Normal Donkey Serum (Jackson ImmunoResearch, 017-000-121) for 1 hour at room temperature following the RNA hybridization protocol. Then, the slides were incubated with primary anti-RFP (Rockland, 600-401-379, 1:500) antibody for 16 hours at 4°C, followed by secondary Alexa-594 antibody (Thermo Fisher Scientific, A32740, 1:1000) for another 1 hour at room temperature. Finally, the sections were counterstained with DAPI (Sigma-Aldrich, D9542, 1:10000) and mounted with antifade mounting solutions in preparation for imaging. In accordance with the ACD guidelines, the RNA levels of target genes identified by RNAscope probes were manually quantified by counting the number of punctate dots in individual cells.

#### Immunohistochemistry

Animals were deeply anesthetized by avertin (2.5% w/v, i.p.) and perfused with PBS followed by 4% PFA. The brains were post-fixed in 4% PFA overnight and dehydrated in a 30% sucrose solution at 4°C. For immunofluorescent staining, the brains were cryosectioned into 40 µm (Leica, CM3050S), blocked with 3% Normal Donkey Serum diluted in PBST (0.1% Triton X-100 diluted in PBS), and incubated with primary anti-c-Fos (Cell Signaling Technology, 2250s, 1:500) antibody for 36 hours at 4°C. Goat anti-Rabbit Alexa 594 (Thermo Fisher Scientific, A32740, 1:1000) was used as the secondary antibody. All sections were imaged using Olympus IXplore Spin Spinning Disk Microscope with 10X objective lens. For cell counting, the images were registered to the standard Allen Mouse Brain Reference Atlas to analyze the c-Fos numbers in different brain regions.

#### Ovariectomy surgery and hormonal priming

Bilateral ovaries were removed by cautery excision. After the incisions were sutured, analgesic meloxicam (4 mg/kg) was administered subcutaneously to the surgical sites for 3 days. Two weeks after ovariectomized (OVX) mice have recovered, they were randomized into control and experimental groups. In order to induce the estrous cycle in the experimental group of OVX mice, 17β-estradiol benzoate (Sigma-Aldrich, E8875) and progesterone (Sigma-Aldrich, P0130) were supplied for three days^107^. Briefly, the experimental group of OVX mice were injected subcutaneously with 10 μg of 17β-estradiol benzoate suspended in 50 μl corn oil (Sigma-Aldrich, C8267) on day 1, followed by 5 μg of 17β-estradiol benzoate dissolved in 50 μl corn oil on day 2, and 50 μg of progesterone suspended in 50 μl corn oil on day 3. Female controls received equivalent amounts of vehicle at identical timepoints. Twenty-four hours after progesterone supplement, the estrus stages of experimental group of OVX females were determined by vaginal smear cytology. Finally, mPFC tissues were dissected from the OVX females of control group and estrus experimental group for RT-PCR.

#### Real-Time PCR

The mPFC tissues were dissected from wild-type mice, Cacna1h-flox mouse line and ovariectomized female mice. Total RNAs were extracted and purified by the QIAwave RNA Mini Kit (QIAGEN, 74536). The first strand cDNAs were reverse transcriped from RNAs by PrimeScript™ RT reagent Kit with gDNA Eraser (TaKaRa, RR047A). Quantitive real-time PCR was performed using PowerUp™ SYBR™ Green (Thermo Fisher Scientific, A25742) in a BioRad system. RNA quantification was analyzed by the delta Ct method with *Gapdh* as an endogenous control. Brain samples of different groups in a same experiment were processed in parallel.

#### Quantification and statistical analysis

Statistical parameters including the sample size, scale bars, statistical test used, and statistical significance were reported in the Figures, the Figure Legends, and the supplementary Table 1. All the data in the figures were judged to be statistically significant when p < 0.05 by two-tailed student’s *t*-test, One-way ANOVA or Two-way ANOVA. In figures, asterisks indicate statistical significance (*, p < 0.05; **, p < 0.01; ***, p < 0.001; ****, p < 0.0001) as compared to controls. For the student’s *t*-test, if the data satisfied the normality, parametric methods were used. But if the data did not follow the normal distribution, the nonparametric Mann-Whitney test was used. All values are presented as mean ± SEM unless otherwise stated. Age-matched mice were randomly assigned to the control or experimental groups. The mice in control group were age-matched littermates whenever possible. All statistics and data analysis were performed by Prism, MATLAB, and R.

---

**Figure S1.**
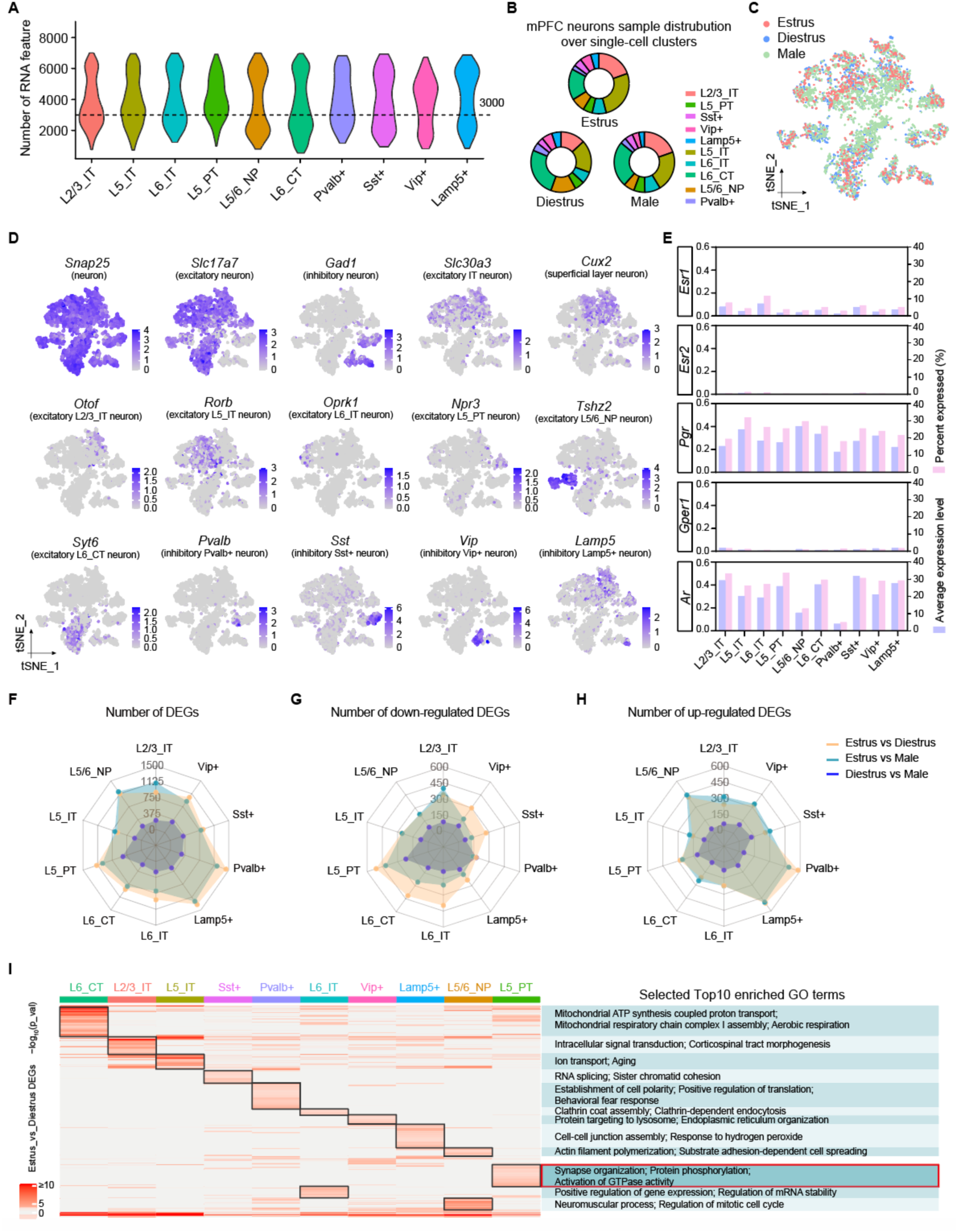
Cell-Type-Specific Gene Expression Analysis in Single-Cell RNA-Seq Data. (A) Violin plot showing the number of genes detected in each neural type. (B) Donut plot showing the proportion of mPFC neuron clusters under the different sample conditions. (C) t-SNE visualization of mPFC neuron clusters, colored according to different sample conditions. (D) t-SNE visualization of marker genes expressed in 10 neural clusters identified in the mPFC. The expression level is depicted from gray (low) to blue (high). (E) The expression level and percentage of hormone receptor genes including Esr1, Esr2, Pgr, Gper1, and Ar, in each mPFC neural clusters (Esr1, estrogen receptor 1; Esr2, estrogen receptor 2; Pgr, progesterone receptor; Gper1, G protein-coupled estrogen receptor 1; Ar, androgen receptor). (F-H) Radial Plot of DEGs in Neural Cluster Comparisons: Showcases counts of all (F), down-regulated (G), and up-regulated DEGs (H) across estrus vs. diestrus, estrus vs. male, and diestrus vs. male. (I) Heat map shows gene modules of cell-type-specific DEGs in estrus versus diestrus comparison of each neuron cluster. Right, selected Top10 enriched GO terms.

**Figure S2.**
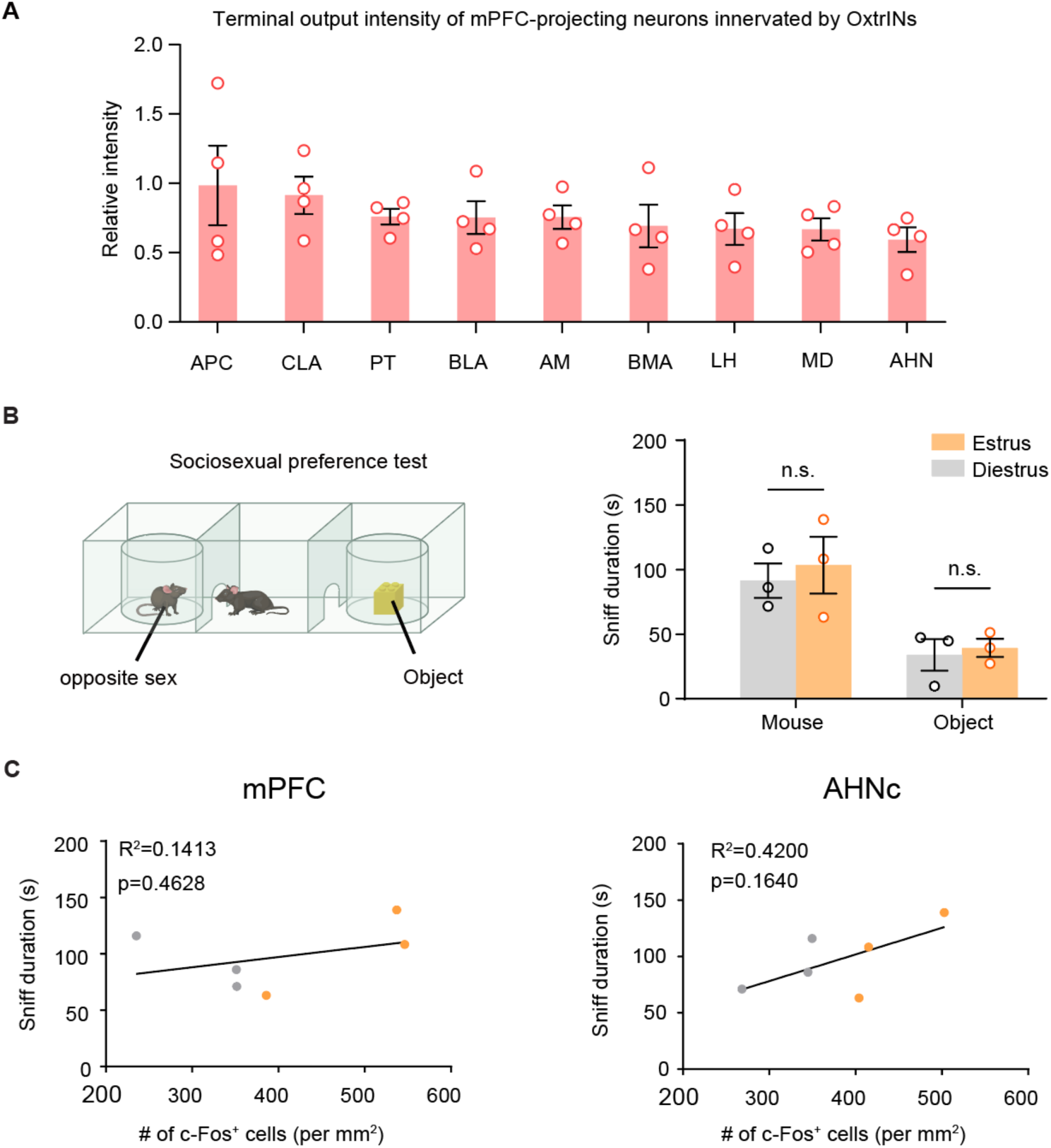
Mapping downstream targets of OxtrIN-Innervated projection neurons in mPFC and analyzing the correlation between social behavior and c-Fos Expression. (A) Quantitative analysis of infection intensity in neurons labeled by the H129ΔTK-TT virus, which receive inputs from OxtrIN-innervated projection neurons located in the mPFC. N = 4 mice. (B) Analysis of male-directed sniffing duration in female mice used for c-Fos labeling. (C) Correlation of c-Fos expression and male-directed sniffing duration in mPFC (left) and AHNc (right).

**Figure S3.**
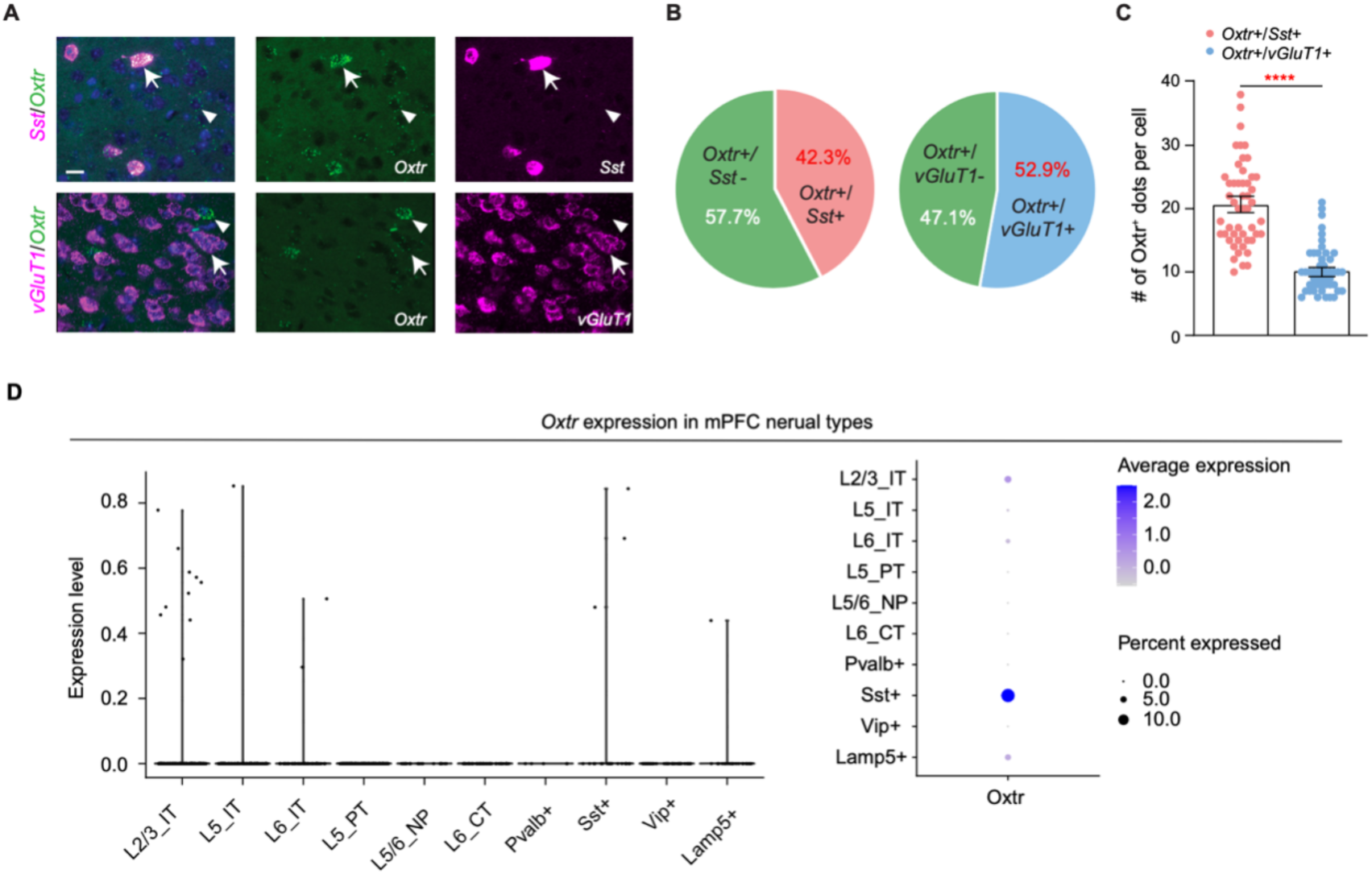
Characterization of the cell identity of Oxtr+ neurons. (A) RNAscope in situ hybridizations of endogenous Sst, vGluT1 and Oxtr in the mPFC. Representative images showing the co-localization of Sst (magenta) and Oxtr (green) mRNAs in the mPFC neurons (Top, arrow indicates Sst+/Oxtr+ neurons; arrowhead indicates Sst-/Oxtr+ neurons; scale bar, 20 µm; DAPI, blue). Representative images showing the co-expression of vGluT1 (magenta) and Oxtr (green) mRNAs in the mPFC neurons (Bottom, arrows indicate vGluT1+/Oxtr+ neurons; arrowheads indicate vGlut1-/Oxtr+ neurons). (B) Pie charts showing the percentage of endogenous Sst+/Oxtr+ neurons and vGlut1+/Oxtr+ neurons in the mPFC respectively. (C) Quantitation of Oxtr+ dots as shown by RNAscope signals in Sst+/Oxtr+ neurons and vGlut1+/Oxtr+ neurons. N = 50 cells per group. Two-tailed unpaired t-test; **** p < 0.0001. (D) Violin plot illustrating the expression levels of the Oxtr gene (left), and a dot plot demonstrating the percentage of cells expressing the Oxtr gene (right) across distinct mPFC neuronal types.

**Figure S4.**
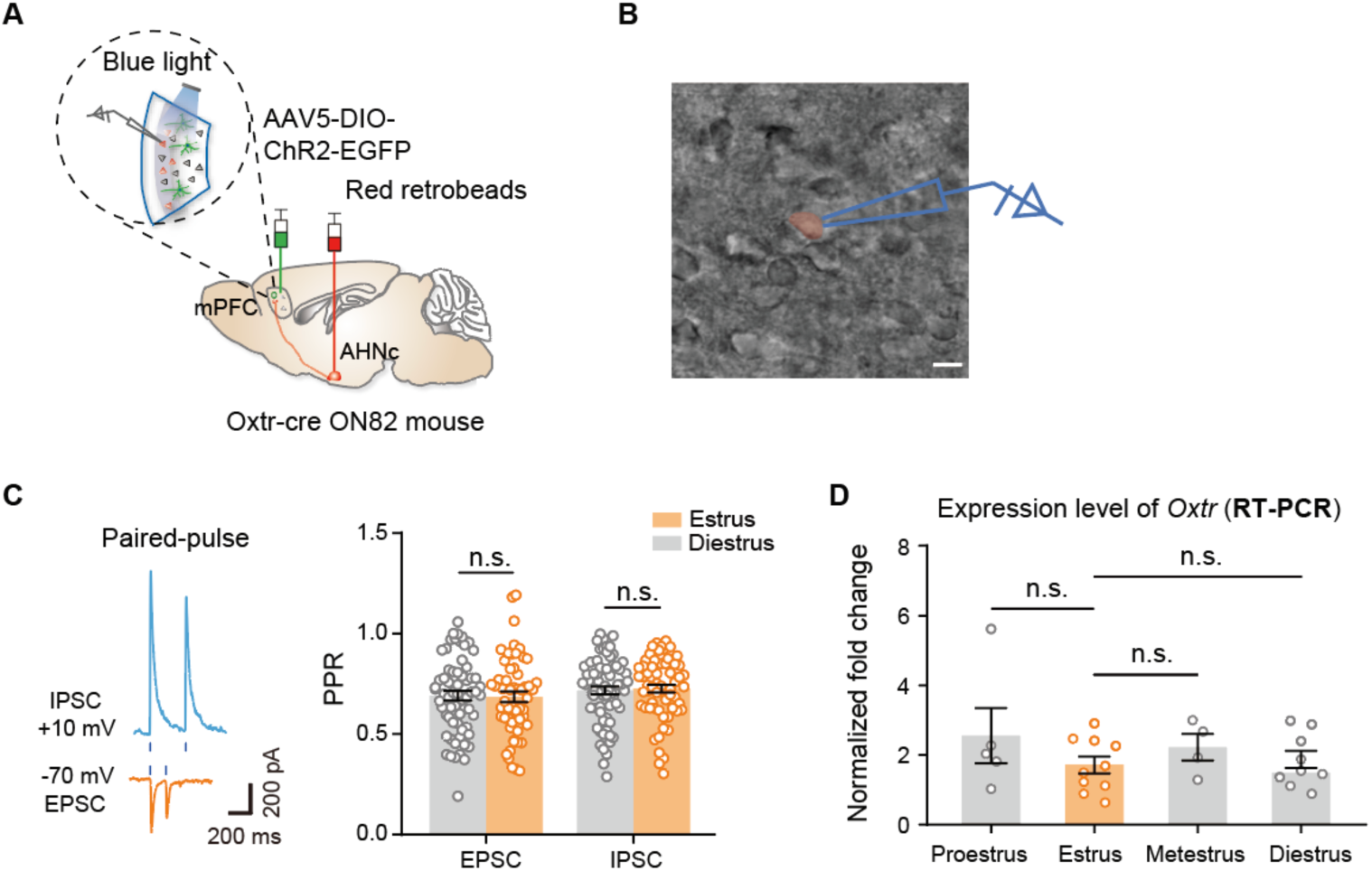
Synaptic transmission between OxtrINs and mPFC^AHN-Projecting^ neurons and mPFC oxtr mRNA expression across estrous phases in female mice. (A) Schematic diagram of the strategy for measuring the paired pulse ratio in mPFC^AHN-^_projecting neurons._ (B) Representative image showing the recording of layer 5 neurons in mPFC brain slice (40×, scale bars, 20 µm). (C) Representative traces of IPSC and EPSC elicited by paired light pulses in AHNc-projecting mPFC layer 5 pyramidal neurons (left). Quantification of the paired-pulse ratio (PPR) of IPSC and EPSC in mPFC^AHN-projecting^ neurons from females in estrus and diestrus groups. The PPR is defined as the second evoked amplitude divided by the first evoked amplitude. N = 56 to 67 cells from 3 mice for each group (right). Two-way ANOVA, Bonferroni multiple comparisons test, n.s. = not significant, p > 0.05. (F) RT-PCR of oxtr mRNA expression in the mPFC of females across four distinct estrous phases. N = 4 to 10 mice for each group. One-way ANOVA, Bonferroni multiple comparisons test, n.s. = not significant, p > 0.05.

**Figure S5.**
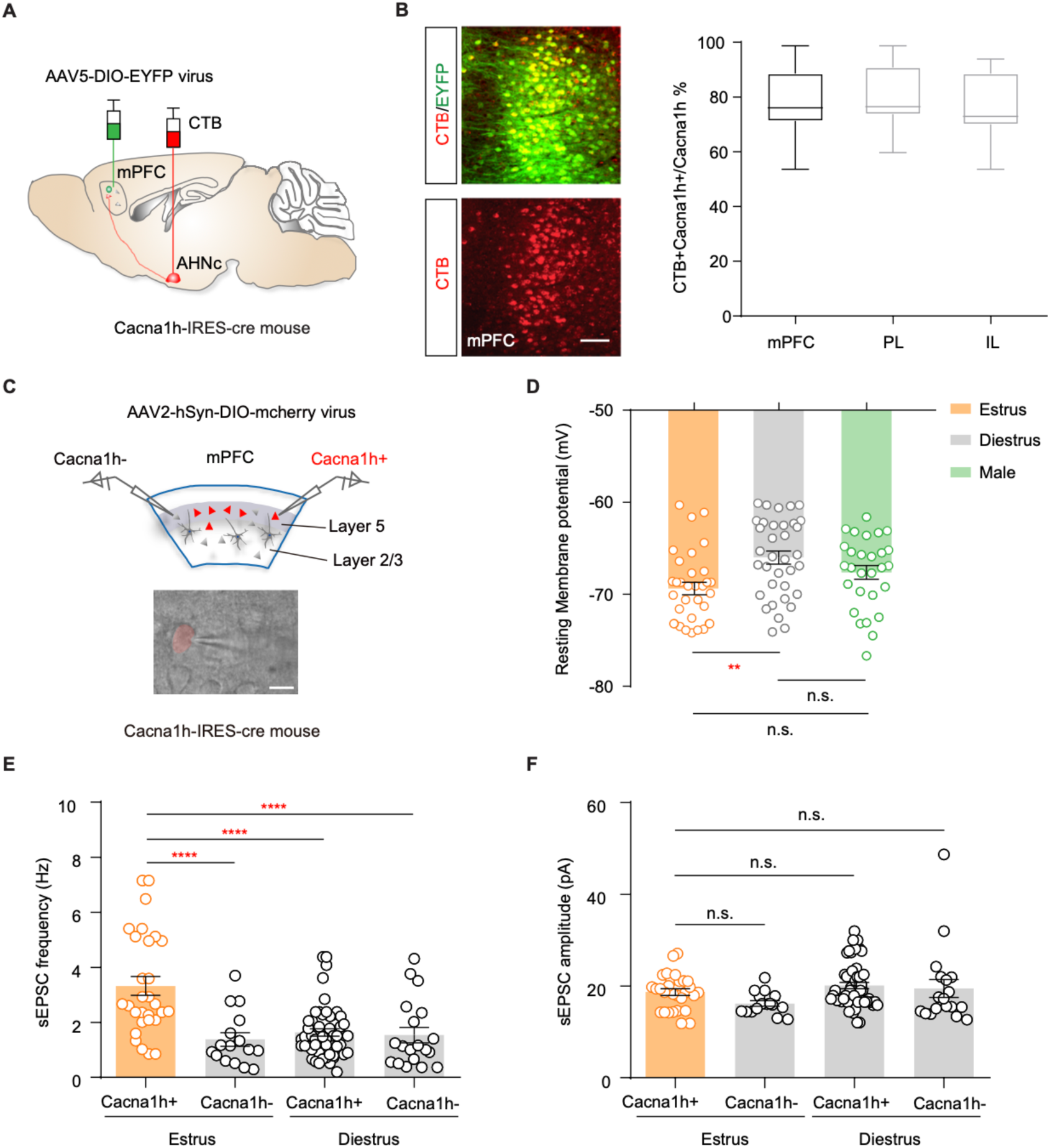
Electrophysiological characterization of mPFC^Cacna1h+^ neurons in estrus and diestrus females. (A) A schematic illustrating the injection of AAV5-DIO-EYFP into the mPFC and CTB-555 into the AHNc of Cacna1h-IRES-cre mouse. (B) Co-localization of EYFP-labeled mPFC^Cacna1h+^ neurons and CTB-positive mPFC^AHNc-projecting^ neurons in the mPFC (left panel); Quantitative analysis of the ratio of AHNc-projecting mPFC^Cacna1h+^ neurons to the total population of *Cacna1h*-expressing neurons in the prelimbic (PL), infralimbic (IL), and overall mPFC regions. (right panel). Scale bar, 100 µm. (C) mPFC scheme showing the recordings in Cacna1h+ neurons and Cacna1h-neurons in mPFC. AAV-DIO-mCherry viruses were introduced into the mPFC of Cacna1h-IRES-cre mouse to label Cacna1h+ neurons. Whole-cell slice recordings were conducted on both mCherry-labeled (Cacna1h+) and unlabeled (Cacna1h-) neurons within the mPFC. Scale bar, 20 µm. (D) Summarized data of the RMP recorded from mPFC^Cacna1h+^ neurons in the indicated groups. N = 19 to 35 cells per group. One-way ANOVA, Bonferroni’s multiple comparisons test, * * p < 0.01, n.s. = not significant, p > 0.05. (E-F) Quantification of sEPSCs frequency (E) and amplitude (F) of mPFC^Cacna1h+^ neurons and mPFC^Cacna1h–^ neurons in estrus and diestrus females. N = 16 to 51 cells in each group. One-way ANOVA, Bonferroni multiple comparisons test, **** p < 0.0001, n.s. = not significant, p > 0.05.

**Figure S6.**
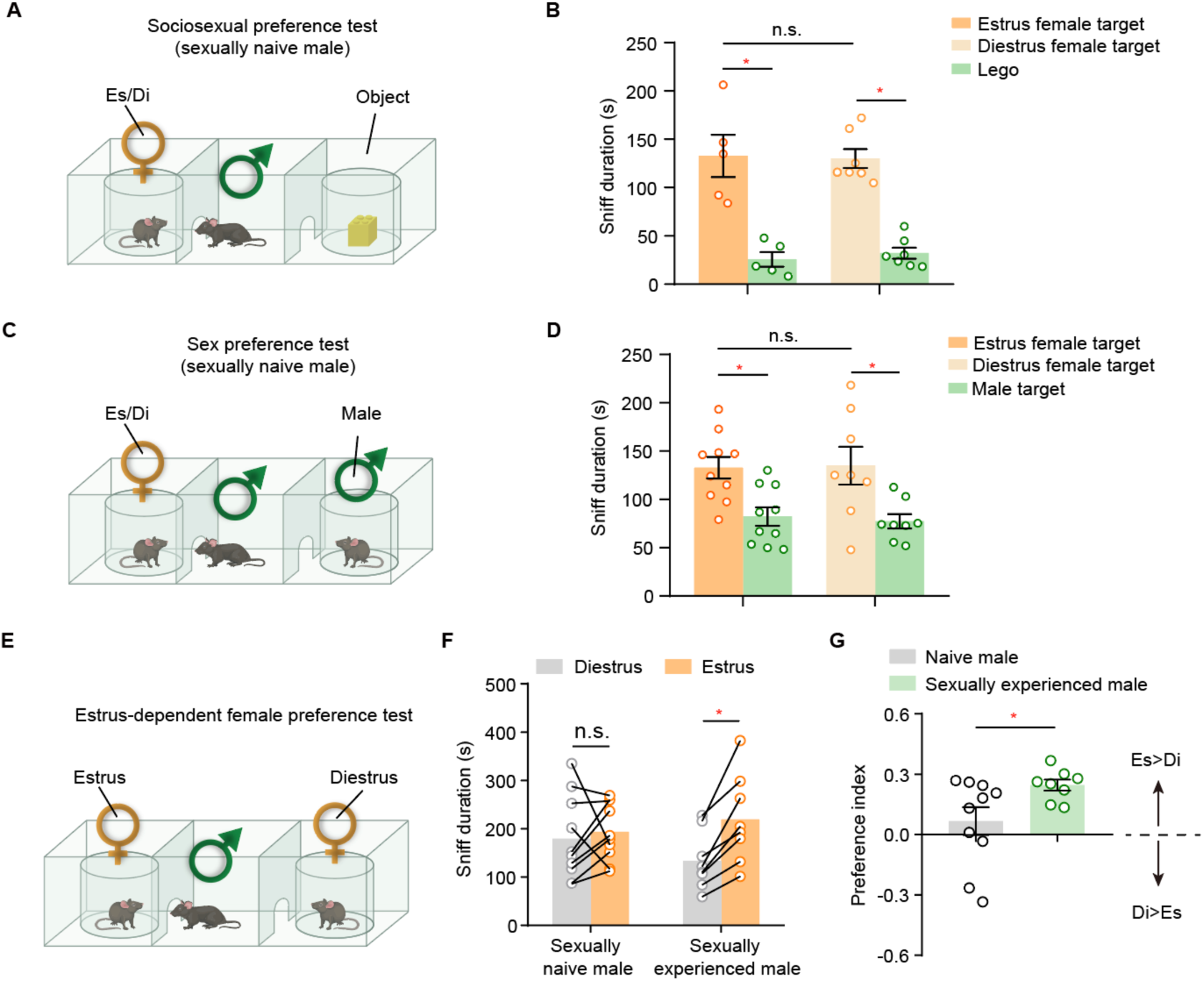
Assessing social preference for estrus and diestrus females in sexually naive and experienced males. (A) Schematic illustrating sexually naive males sniffing an estrus female or diestrus female and an object in sociosexual preference test. (B) Quantification of male sniffing behaviors towards female or object in sociosexual preference test. N = 5 to 10 mice for each group. Two-tailed paired *t*-test, n.s. = not significant, p > 0.05, * p < 0.05. (C) Schematic illustrating sexually naive males sniffing an estrus female or diestrus female and a male in the sex preference test. (D) Quantification of sniff duration by sexually naive males toward estrus or diestrus females and males in sex preference test. N = 5 to 10 mice for each group. Two-tailed paired *t*-test, n.s. = not significant, p > 0.05, * p < 0.05. (E) Schematic illustrating sexually naive and experienced males sniffing estrus and diestrus females during a mating preference test. (F-G) Quantification of sniff duration toward estrus and diestrus females (F) and calculation of preference index (G) in sexually naive and experienced males during the estrus-dependent female preference test. N = 8-10 mice for each group. Two-tailed paired t-test, n.s. = not significant, p > 0.05, * p < 0.05.

**Figure S7.**
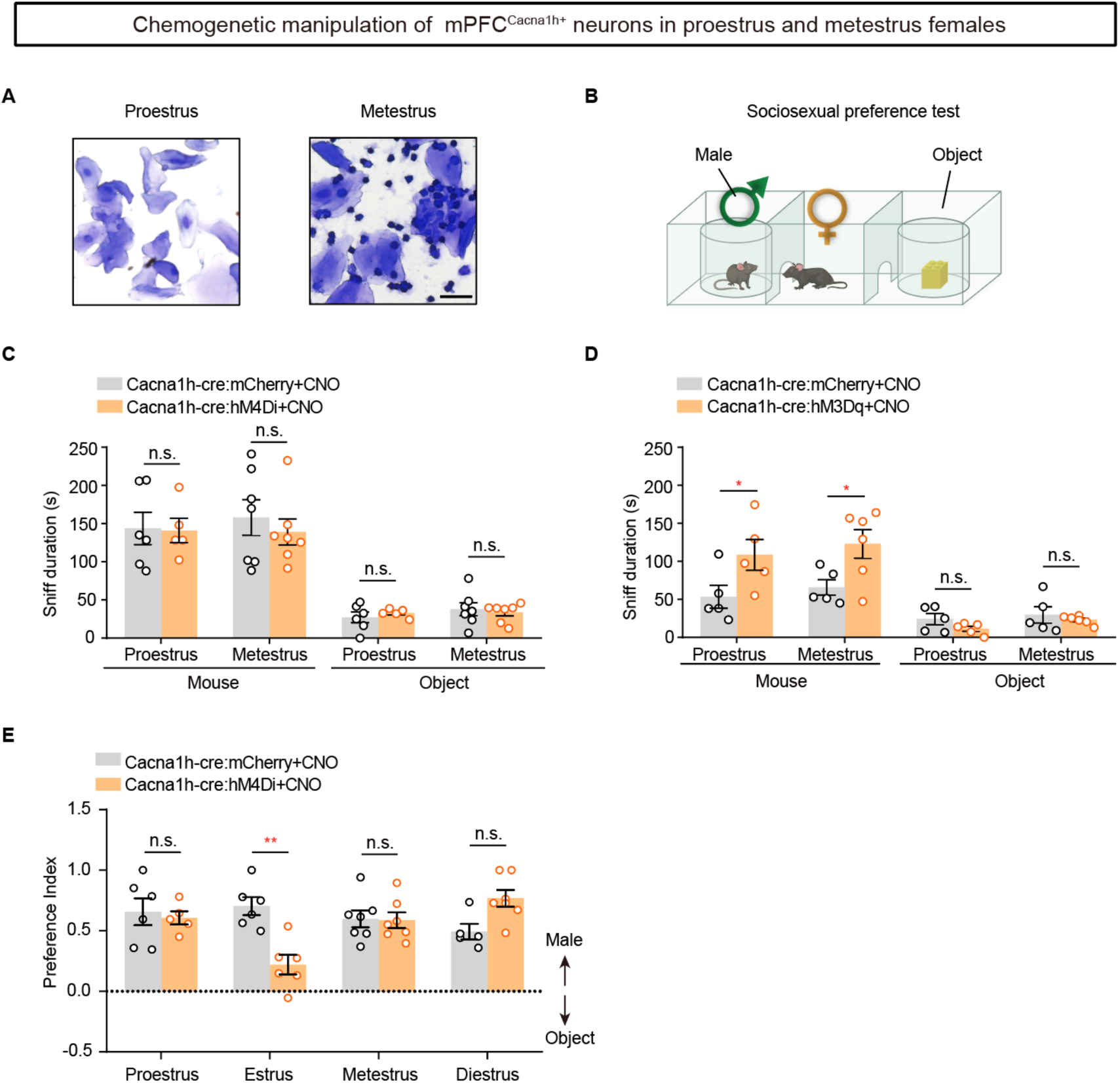
Impact of chemogenetic manipulation of mPFC^Cacna1h+^ neurons on female sociosexual interest in proestrus and metestrus phases. (A) Representative images illustrating proestrus and metestrus stages identified through vaginal cytology. Scale bar, 25 µm. (B) Schematic of chemogenetic manipulation of mPFC^Cacna1h+^ neurons in sociosexual preference assays. (C) Chemogenetic inhibition of mPFC^Cacna1h+^ neurons in female mice does not alter their investigation toward males during proestrus and metestrus phases. N = 5 to 7 mice for each group. Two-way ANOVA, Bonferroni multiple comparisons test, p > 0.05. (D) Chemogenetic activation of mPFC^Cacna1h+^ neurons in females increases the social interest towards males during both proestrus and metestrus phases. N = 5 to 6 mice for each group. Two-way ANOVA, Bonferroni multiple comparisons test, * p < 0.05. (E) Chemogenetic inhibition of mPFC^Cacna1h+^ neurons in female mice reduces the social preference index toward males only during estrus phase, but not in other phases. N = 5 to 7 mice for each group. Two-way ANOVA, Bonferroni multiple comparisons test, p > 0.05.

**Figure S8.**
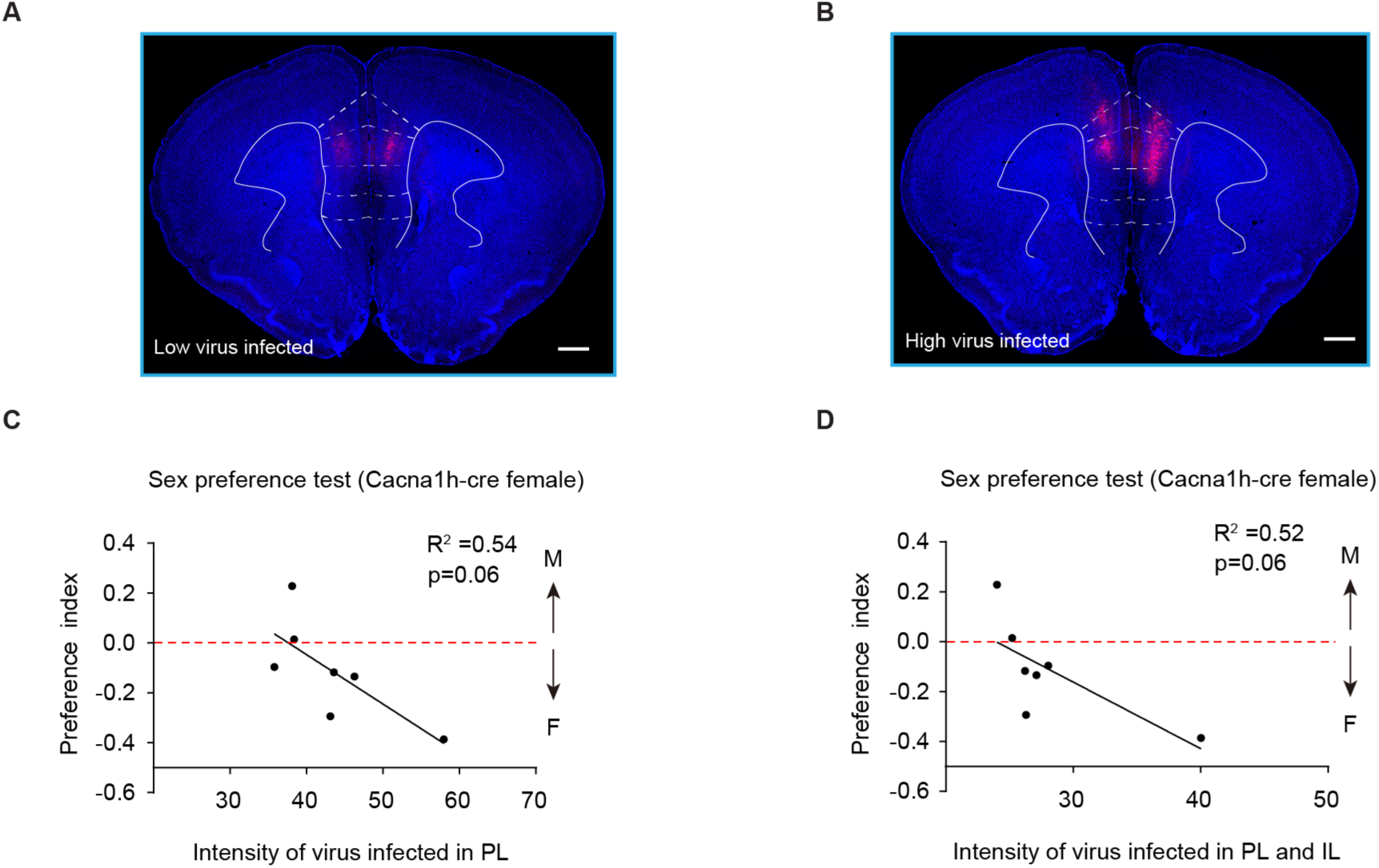
Correlation of viral infection with behavioral modifications in estrus females. (A-B) Representative confocal images of Cacna1h-Cre::hM4Di mice showing low (A) and high (B) hM4Di-mCherry virus expression in the mPFC. Scale bars, 500 μm. (C-D) Graphs illustrating the intensity infected by hM4Di-mCherry virus in the prelimbic (PL) (C) and the combined PL and infralimbic (IL) regions (D) of estrus females, plotted against the preference index.

**Figure S9.**
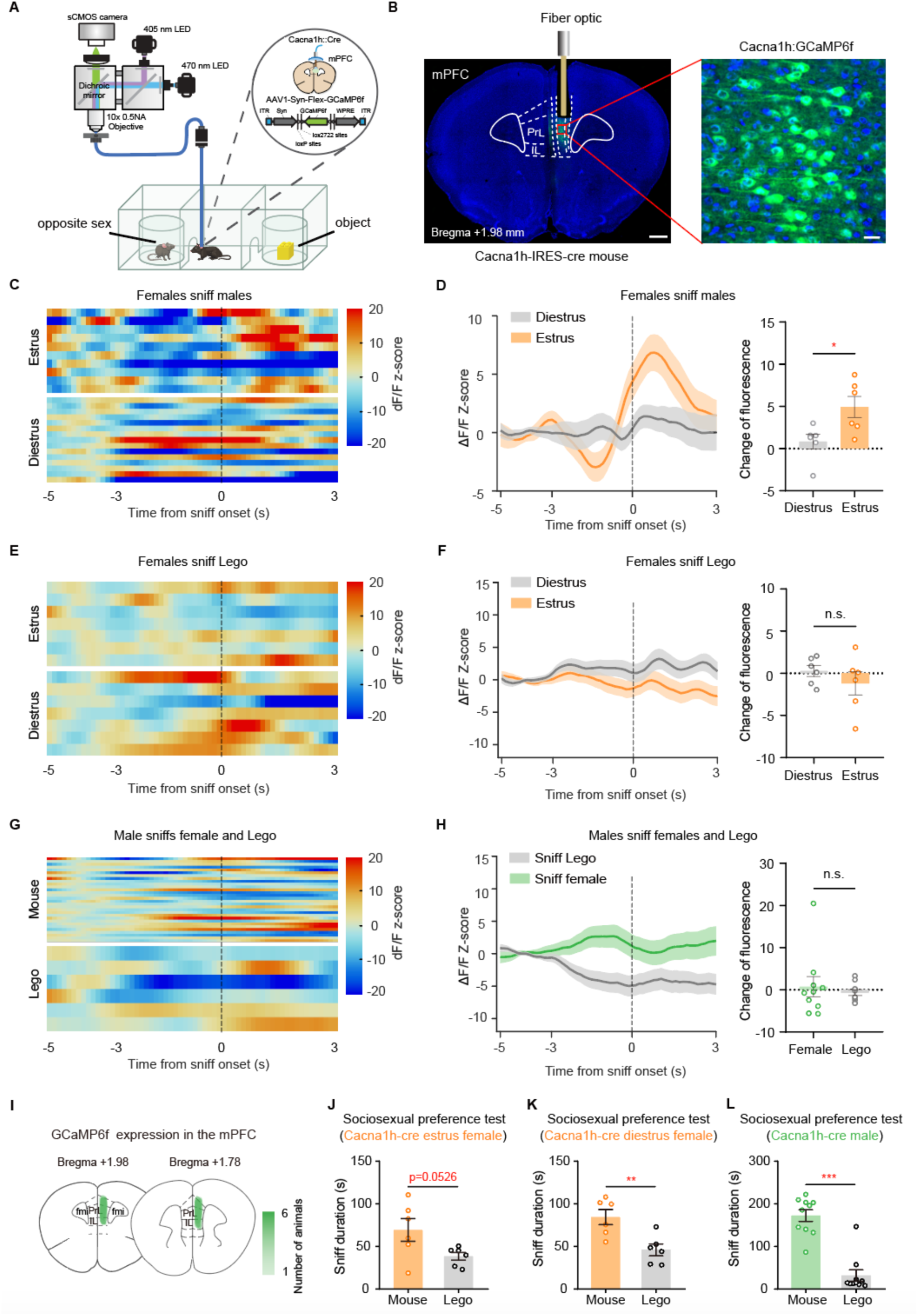
Calcium response in mPFC^Cacna1h+^ neurons to sniffing the opposite sex and objects across estrus and diestrus phases in females and males. (A) Scheme illustrating the fiber photometry setup to record calcium signal of mPFC^Cacna1h+^ neurons during the sociosexual preference test. (B) Coronal section showing the expression of AAV1-Syn-Flex-GCaMP6f in the mPFC of Cacna1h-IRES-cre mice with an optic fiber implanted above the virus infection site (left). The GFP-positive cells represent GCaMP6f-expressing mPFC^Cacna1h+^ neurons (right). Blue, DAPI; scale bars, 500 µm (left) and 20 µm (right). (C) Heat map of GCaMP6f signals in mPFC^Cacna1h+^ neurons as an estrus female (top) and a diestrus female (bottom) sniff a male. The GCaMP6f signal is aligned to the time point of sniffing onset. The color scale is determined by the normalized Z-score. (D) Peri-event plot of Z-scored dF/F GCaMP6f signals when estrus and diestrus females sniffing males (left). Quantitative analysis of change in calcium signals before and after the onset of sniffing males (right). n = 6 mice for each group. Solid lines indicate mean. Shaded areas indicate SEM. Two-tailed unpaired t-test; p = 0.0288, * p < 0.05. (E) Heat map of GCaMP6f signals in mPFC^Cacna1h+^ neurons as an estrus female (top) and a diestrus female (bottom) sniff object. (F) Peri-event plot of Z-scored dF/F GCaMP6f signals when estrus and diestrus females sniffing Legos (left). Quantitative analysis of change in calcium signals before and after the onset of sniffing objects (right). n = 6 mice for each group. Solid lines indicate mean. Shaded areas indicate SEM.Two-tailed unpaired t-test; n.s. = not significant, p > 0.05. (G) Heat map of GCaMP6f signals in mPFC^Cacna1h+^ neurons as a male sniff a female (top) and a object (bottom). (H) Peri-event plot of Z-scored dF/F GCaMP6f signals when males sniffing females and objects (left). Quantitative analysis of change in calcium signals before and after the sniffing onset (right). n = 10 mice per group. Solid lines indicate mean. Shaded areas indicate SEM. Two-tailed Mann Whitney test; n.s. = not significant, p > 0.05. (I) The sample brain slices illustrating the infection site of AAV1-Syn-Flex-GCaMP6f-WPRE-SV40 virus in the mPFC. (J-L) Quantification of sociosexual preference behavior in fiber photometry-recorded estrus female (J), diestrus female (K), as well as male mice (L). N = 6 to 10 mice for each group. Two-tailed unpaired t-test (J-K) and Mann-Whitney test (L), * * p < 0.01, * * * p < 0.001.

**Figure S10.**
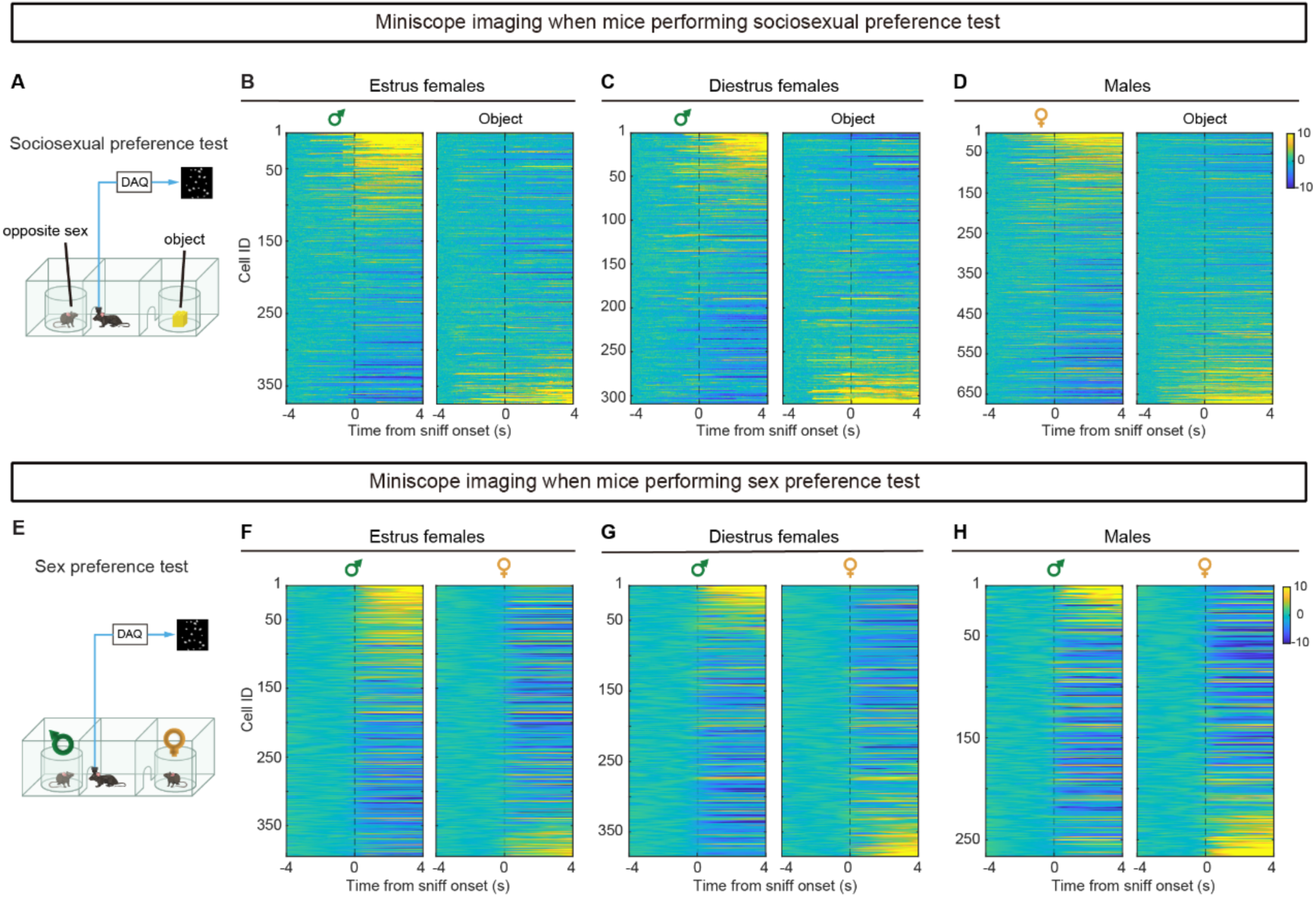
Miniscope imaging reveled the averaged activity of individual mPFC^Cacna1h+^ neurons during sociosexual and sex preference tests. (A) Schematic of the experimental setup for imaging neuronal activity during sociosexual preferene test. (B-D) Heatmap of average peri-event calcium signals in individual mPFC^Cacna1h+^ neurons as estrus females (B), diestrus females (C) or males (D) sniff mouse of opposite sex and object, ranked by maximum response during 0 to 4s after sniff onset. (E) Schematic of the experimental setup for imaging neuronal activity during the sex preference test. (F-H) Heatmap of mPFC^Cacna1h+^ neuronal responses to sniffing female and male conspecifics in estrus females (F), diestrus females (G) and males (H). Peri-event calcium signals aligned to sniff onset (0 s).

**Figure S11.**
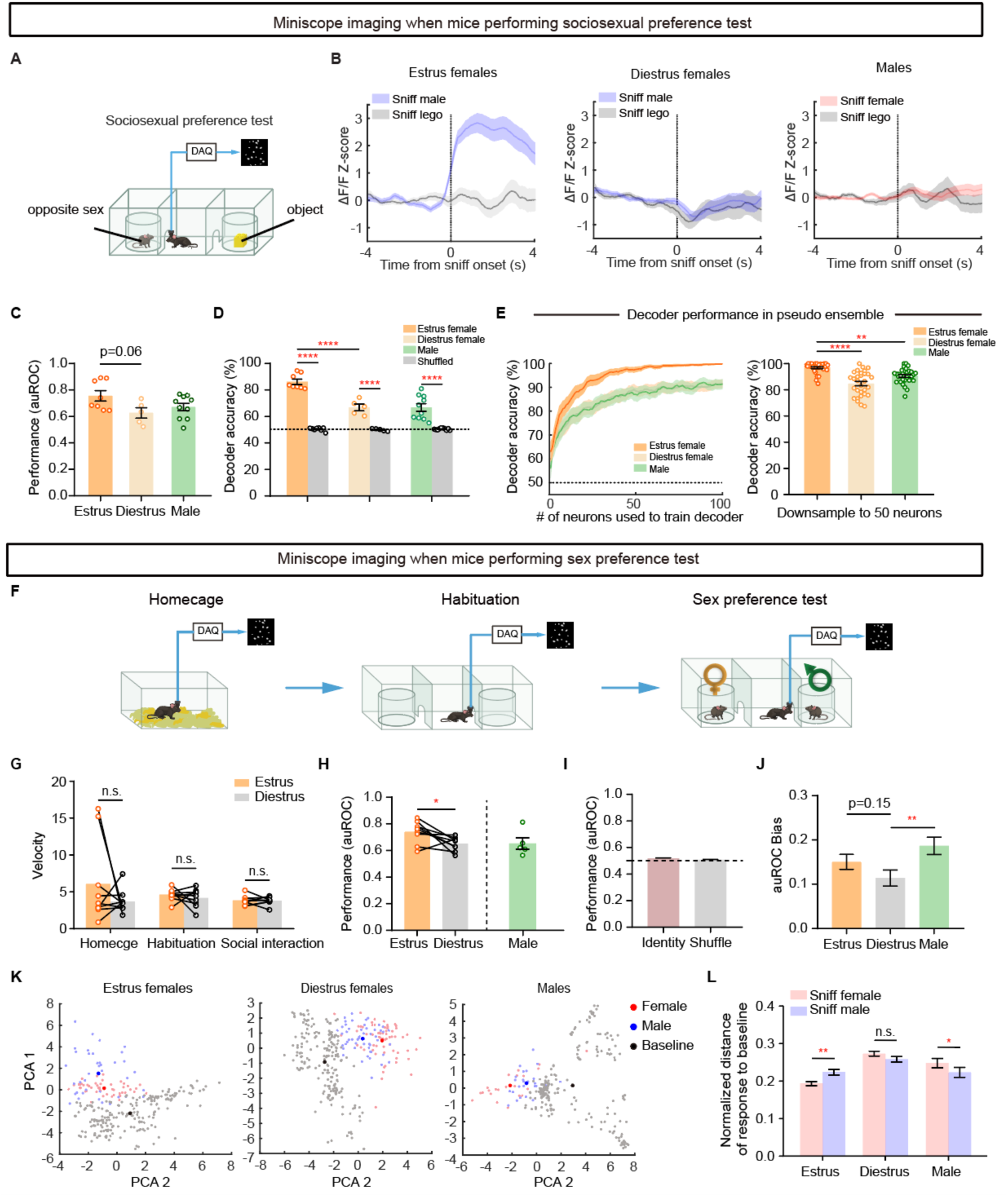
Miniscope imaging of mPFC^Cacna1h+^ neurons when mice performing sociosexual and sex preference tests. (A) Schematic illustrating calcium imaging of mPFC^Cacna1h+^ neurons via Miniscope during sociosexual preference testing. (B) Bulk calcium signals from recorded neurons in estrus, diestrus females, and male mice during sniffing of opposite sex and objects (C) Performance of unsupervised classifier (K-means clustering) distinguishing sniffing opposite sex vs. object using neural response during 0 to 2s relative to sniffing onset in estrus, diestrus females and males. (D) SVM decoder performance distinguishing sniffing opposite sex vs. object based on neural responses 0-2s post-sniff in estrus, diestrus females, and males. (E) Classifier performance as a function of ensemble size (1-100) trained with pseudo neural ensembles recorded in estrus, diestrus females, and males (left). Classifier performance in randomly sampled 50 neurons from pseudo neuron ensembles recorded in estrus, diestrus females and males (right). (F) Schematic illustrating calcium imaging of mPFC^Cacna1h+^ neurons with Miniscope in the home cage, during habituation in a three-chamber setup, and investigation in sex preference test. (G) Velocity of estrus and diestrus females in home cage, during habituation, and in social investigation phases. (H) SVM classifier performance distinguishing male vs. female targets based on neural responses 0-2s post-sniff onset in estrus females, diestrus females, and males. N=9 estrus females, N=9 diestrus females, N=5 males. (I) Performance of SVM classifier in differentiating same-sex targets to assess influence of mouse identity rather than sex on classifier accuracy, compared against label-shuffled data. N=20 animals, 8 estrus females, 8 diestrus females, and 4 males. (J) The auROC bias of same-sex excited neurons towards opposite-sex stimuli, calculated as |auROC -0.5|. N=38 estrus neurons, N=31 diestrus neurons, N=34 male neurons. (K) Example scheme of average male (blue) and female (red) sniffing trial activities within 2s post-onset and a 10s random baseline (gray, non-behavior time) projected onto a 2-dimensional principal component plane. (L) Euclidian distance comparison of all trails from baseline in estrus, diestrus females and males. N=417 female bouts vs. 353 male bouts in estrus females, 425 female bouts vs. 315 male bouts in diestrus females, 215 female bouts vs. 201 male bouts in males.

**Figure S12.**
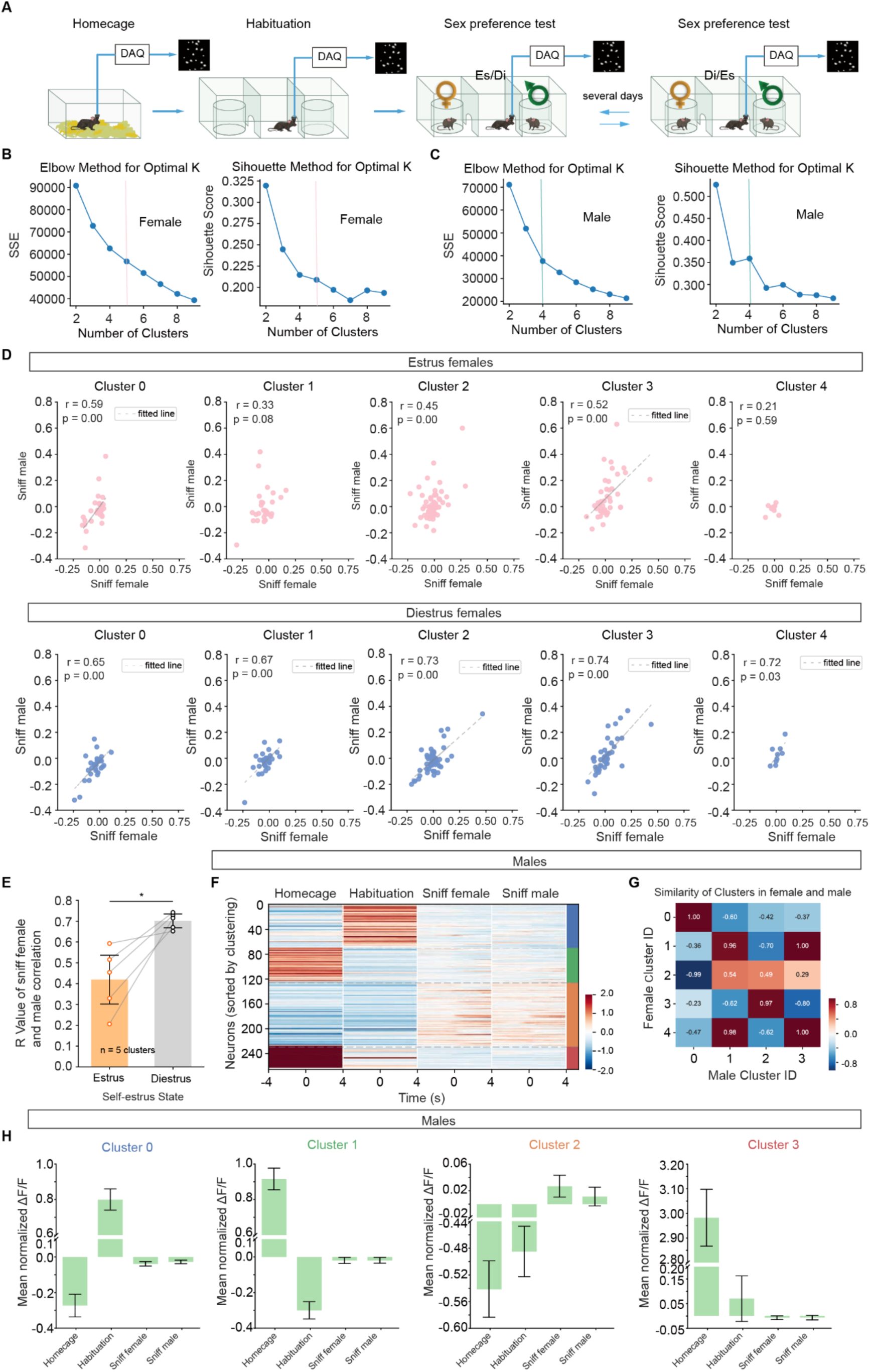
Distinct mPFC^Cacna1h+^ neuron clusters exhibit sex- and estrus-dependent responses to social stimuli. (A) Timeline of sex preference test. (B-C) K-means clustering evaluation for female (B) and male (C) mPFC^Cacna1h+^ neurons using sum of squared errors (SSE) and silhouette score. Optimal K determined by the elbow point in SSE and the local maximum in silhouette score. (D) Linear regression of mPFC^Cacna1h+^ neuron activity of each cluster in response to sniffing female and male stimuli during estrus (top) and diestrus (bottom). The fitted line is plotted only when r-value>0.5 and p-value<0.05. (E) Difference in correlation coefficients (r-values) between mPFC^Cacna1h+^ neuron activity during sniffing female and male stimuli across estrus states. (F) Clustering of mPFC^Cacnah1+^ neurons in male. Similar to Figure 5L, colors on the right indicate different clusters. (G) Similarity matrix of clusters identified in female and male mPFC^Cacna1h+^ neurons, based on Pearson correlation coefficients (H) Mean activity of mPFC^Cacna1h+^ neurons of each cluster in males.

**Figure S13.**
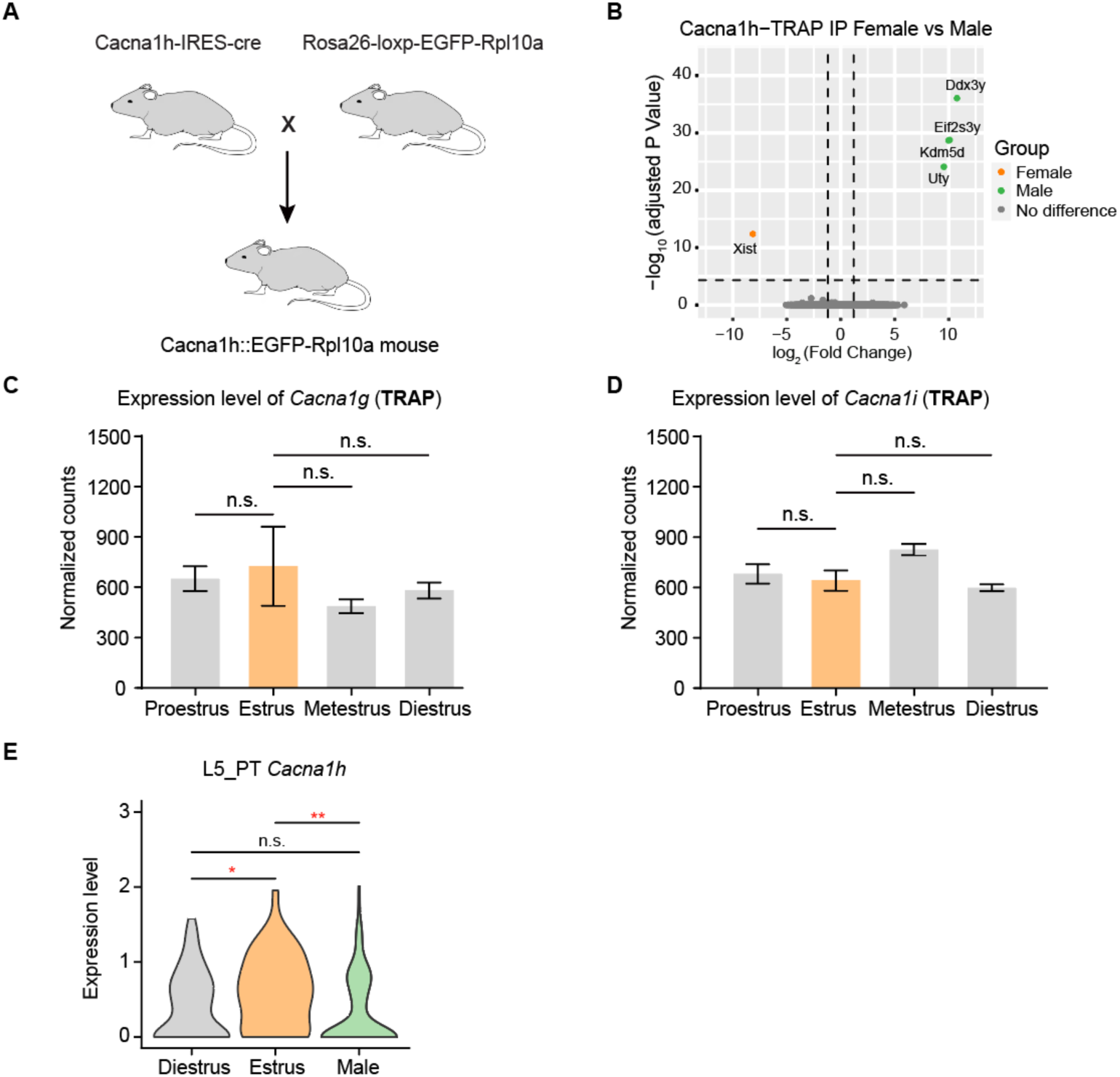
TRAPseq analysis of mPFC^Cacna1h+^ IP samples and RT-PCR verification of gene expression in the mPFC. (A) Cacna1h-IRES-cre mouse was bred to Rosa26-loxp-EGFP-Rpl10a mouse to generate Cacna1h-IRES-cre::Rpl10a-EGFP mouse. (B) Volcano plot for genes enriched in male versus female mPFC^Cacna1h+^ IP samples identified by TRAPseq. Green dots denote genes significantly enriched in male mPFC^Cacna1h+^ neurons with adjusted p < 0.05 and log_2_ fold change > 1.2, and orange dots denote genes significantly enriched in the female mPFC^Cacna1h+^ neurons with adjusted p <0.05 and log_2_ fold change < -1.2. N = 8 female samples and 3 male samples. (C-D) TRAP analysis of *Cacna1g* (C), and *Cacna1i* (D) expression in the mPFC of females across estrous phases. N = 2 to 4 samples per group. One-way ANOVA, Bonferroni multiple comparisons test, n.s. = not significant, p > 0.05. (E) Comparison of *Cacna1h* expression between diestrus, estrus female, and male in the L5_PT cell type identified by scRNA-seq.

**Figure S14.**
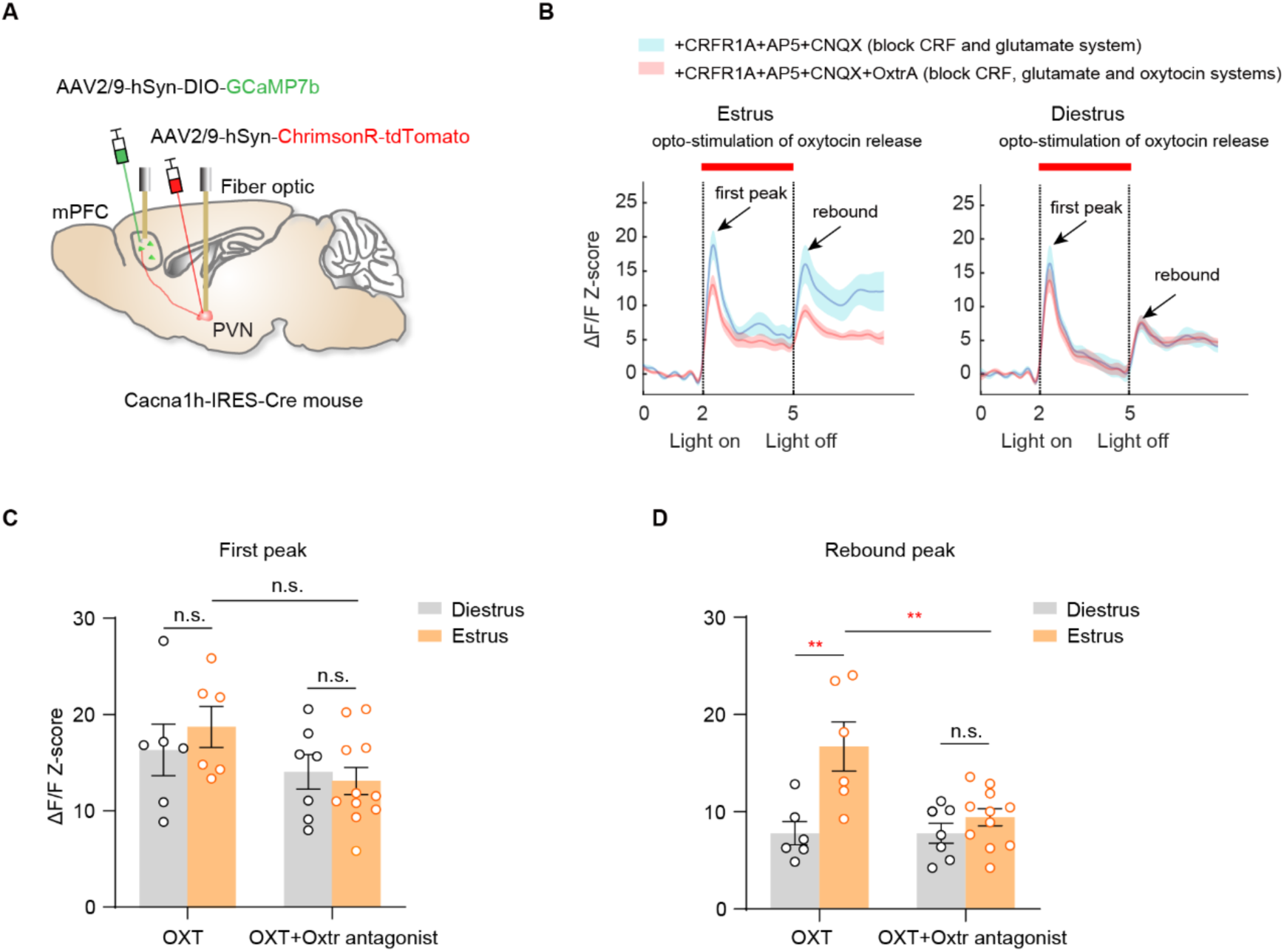
Oxytocin release elicits enhanced rebound activity in mPFC^Cacna1h+^ neurons during estrus compared to diestrus. (A) Schematic of viral injections and fiber implantations in Cacna1h-IRES-Cre mice. AAV2/9-hSyn-ChrimsonR-tdTomato was injected into the PVN for optogenetic activation of oxytocin release, and AAV2/9-hSyn-DIO-GCaMP7b was injected into the mPFC for calcium imaging of Cacna1h+ neurons. Optic fibers were implanted in the PVN for optogenetic stimulation and in the mPFC for fiber photometry recording of bulk Cacna1h+ neuron activity. (B) Average calcium activity of mPFC^Cacna1h+^ neurons in response to 3 s light stimulation of PVN neurons with pharmacological blockade of CRF, glutamate, or oxytocin signaling in estrus and diestrus female mice. The first peak following light onset suggests disinhibition of mPFC^Cacna1h+^ neurons, while the second peak after light offset indicates rebound activity. The right bar denotes the light stimulation period. (C-D) Quantification of Z-scored ΔF/F GCaMP7b signals for the first (C) and rebound (D) peaks of mPFC^Cacna1h+^ neuron activity in estrus and diestrus female mice. The amplitude of rebound activity, induced by oxytocin stimulation of inhibitory Oxtr-expressing mPFC neurons, was significantly higher in mPFC^Cacna1h+^ neurons during estrus compared to diestrus phases. Oxtr antagonist treatment abolished the increased rebound activity in estrus.

**Figure S15.**
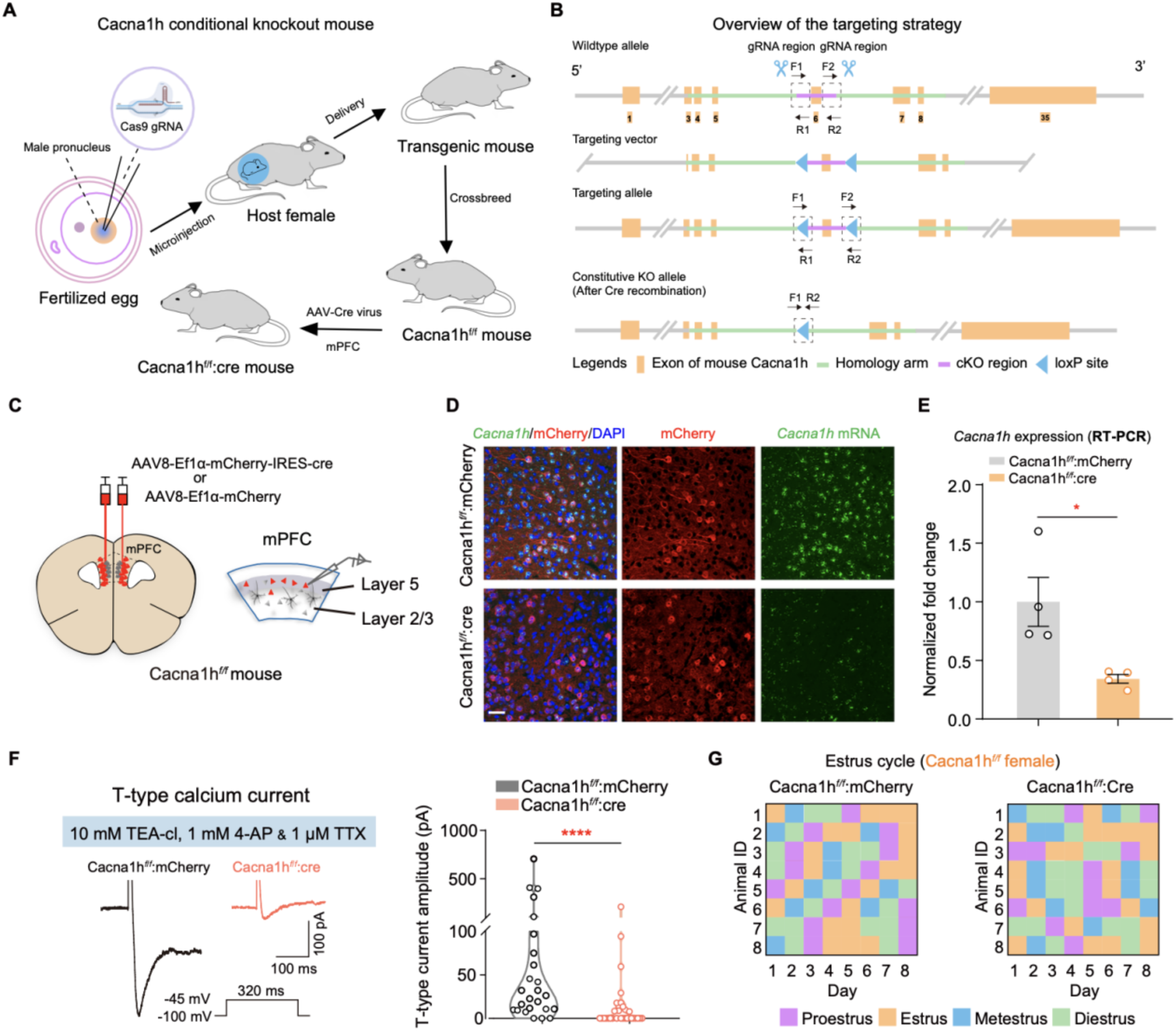
Generation and validation of Cacna1h-flox mice. (A-B) Schematic representation of conditional Cacna1h knockout in the mPFC (A). Overview of the strategy for constructing the Cacna1h^fl/fl^ mouse line to conditionally knockout *Cacna1h* (B). The targeting vector was designed to introduce loxP sites flanking the exon 6 of the *Cacna1h* gene. Through homologous recombination in embryonic stem cells, the loxP sites were inserted into the genome. Correctly targeted embryonic stem cell clones were selected and used to generate chimeric mice, which were then bred with wild-type mice to obtain heterozygous Cacna1h^flox/+^ mice. Further breeding of heterozygous mice resulted in the establishment of the homozygous Cacna1h^fl/fl^ mouse line, (C) Schematic illustrating the injection of AAV8-Ef1α-mCherry-IRES-cre or AAV8-Ef1α-mCherry into the mPFC of Cacna1h^fl/fl^ mouse. (D) Representative images of *Cacna1h* mRNA and virus expression in the mPFC of Cacna1h^fl/fl^ mice. Scale bar: 50 μm. (E) Quantification of *Cacna1h* mRNA expression in the mPFC using RT-PCR in Cacna1h^fl/fl^:mCherry and Cacna1h^fl/fl^:Cre groups. N = 4 mice for each group. Two-tailed unpaired *t*-test; * p < 0.05. (F) Electrophysiological validation of the conditional *Cacna1h* knockout in mPFC neurons. Representative traces of T-type calcium currents recorded from layer 5 pyramidal neurons in the mPFC slices of control (Cacna1h^fl/fl^) and conditional *Cacna1h* knockout (Cacna1h^fl/fl^; Cre) mice (left). Quantification of the peak T-type calcium current amplitude in layer 5 pyramidal neurons of the mPFC in control (Cacna1h^fl/fl^) and conditional Cacna1h knockout (Cacna1h^fl/fl^: Cre) mice (right). N = 25 to 30 cells per group. Two-tailed Mann-Whitney test, **** p < 0.0001. (G) Layout of the estrus states of Cacna1h-flox female mice assigned by vaginal smear in eight days (proestrus = purple; estrus = orange; metestrus = blue; diestrus = green).

**Figure S16.**
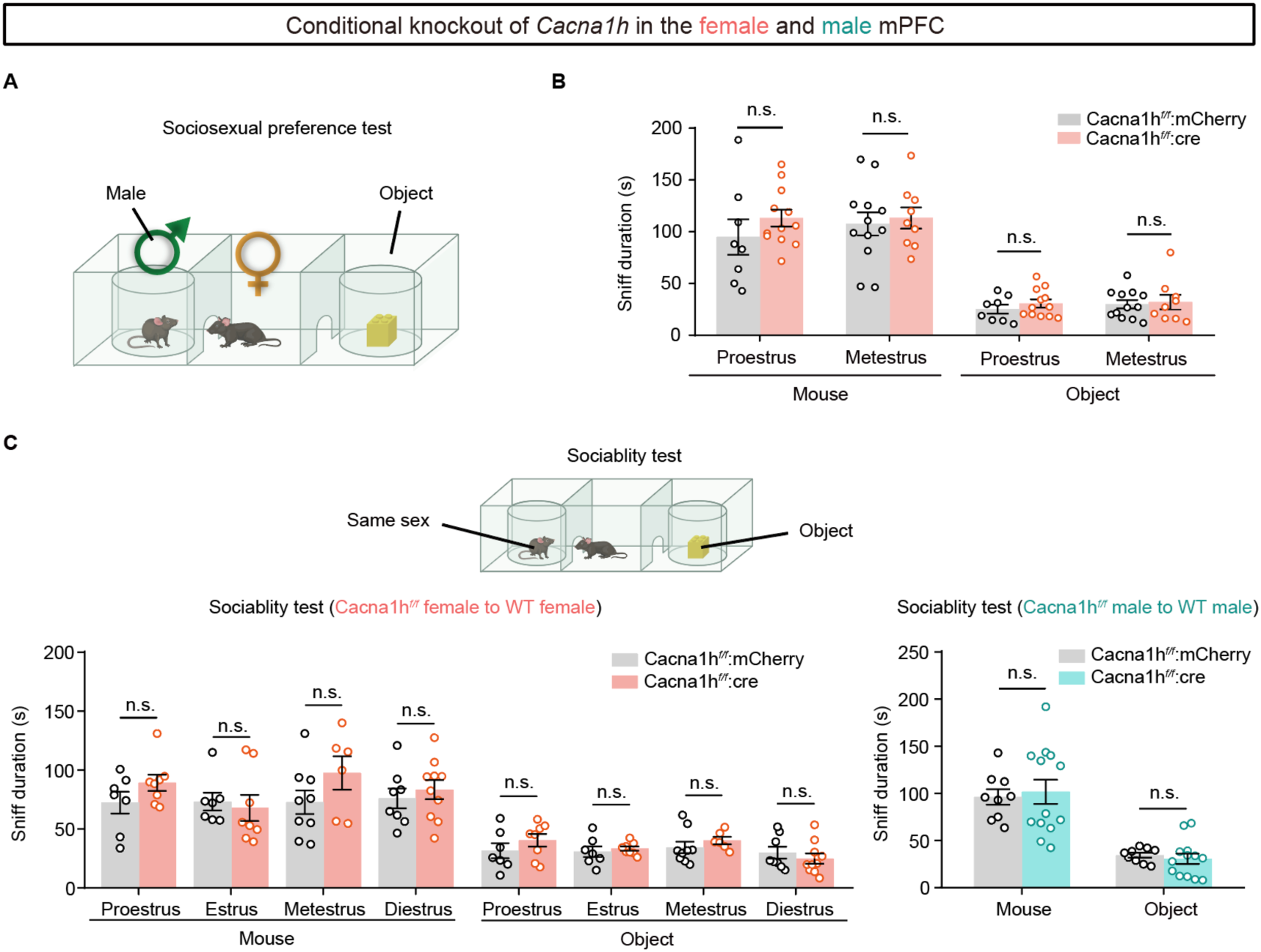
*Cacna1h* deletion in the mPFC did not affect opposite-sex investigation in non-estrus females or same-sex investigation in either sex. (A) The strategy of sociosexual preferene test. (B) Conditional deletion of *Cacna1h* in the mPFC of female mice does not affect the duration of sniffing opposite sex during proestrus and metestrus phases. N = 7 mice for each group. Two-way ANOVA, Bonferroni multiple comparisons test, p > 0.05. (C) Conditional deletion of *Cacna1h* in the mPFC of female and male mice did not affect sniffing same-sex social targets in females across extrous cycles (left) and males (right). N = 6 to 13 mice for each group. Two-way ANOVA, Bonferroni multiple comparisons test, n.s. = not significant, p > 0.05.

**Figure S17.**
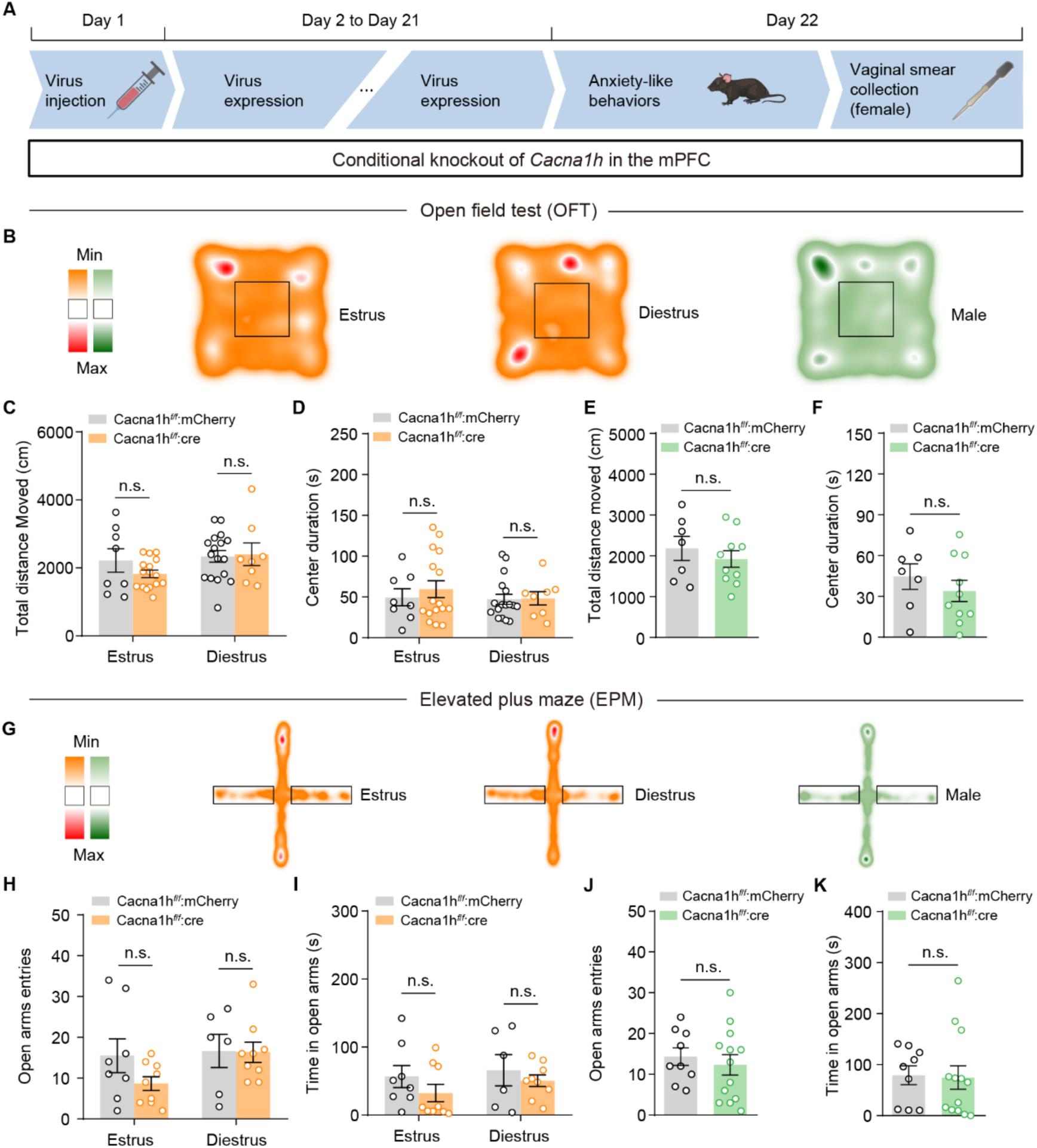
Conditional knockout of *Cacna1h* in the mPFC does not affect locomotor activity and anxiety-like behaviors in both female and male mice. (A) Experimental timeline of behavioral tests on female and male mice. (B) Representative heat maps showing all the paths traveled during the open field test (OFT) by each group. Red and dark green represent high dwell time in the compartment of open field box. Orange and light green represent low dwell time in open field box. (C-D) Conditional knockout of *Cacna1h* in the mPFC did not affect the total distance moved (C) and center duration (D) in the OFT in both estrus and diestrus female mice. N = 8 to 17 mice for each group. Two-way ANOVA, Bonferroni multiple comparisons tests, n.s. = not significant, p > 0.05. (E-F) Male mice with mPFC-specific *Cacna1h* deletion showed no differences in total distance moved (E) or center duration (F) during open field test compared to control males. N = 7 to 10 mice for each group. Two-tailed unpaired *t*-test, n.s. = not significant, p > 0.05. (G) Representative heat maps of all the paths traveled during the elevated plus maze (EPM) in both male and female groups. (H-I) mPFC-specific *Cacna1h* deletion does not affect the frequency (H) or duration (I) of open arm entries in the EPM test for both estrus and diestrus female mice. N = 6 to 9 mice for each group. Two-way ANOVA, Bonferroni multiple comparisons tests, n.s. = not significant, p > 0.05. (J-K) Conditional knockout of *Cacna1h* in the mPFC of male mice did not affect the open arm entries (J) and duration (K) in the EPM test compared to the control male mice. N = 9 to 13 mice for each group. Two-tailed unpaired *t*-test (J) and Mann-Whitney test (K), n.s. = not significant, p > 0.05.

**Figure S18.**
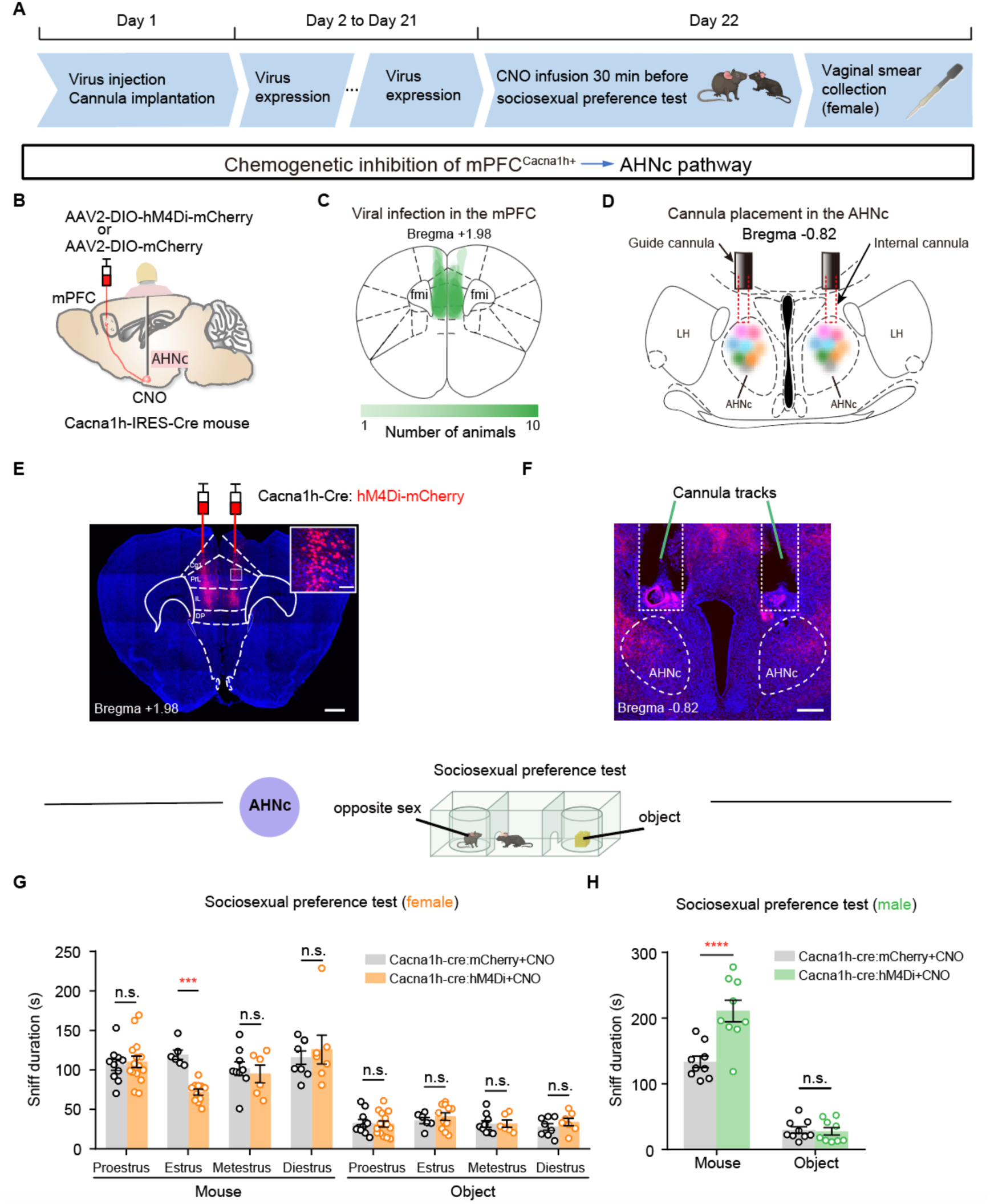
mPFC^Cacna1h+^ neurons drive sexually dimorphic top-down of control sociosexual interest via AHNc-descending pathways. (A) Experimental procedure for chemogenetic inhibition of mPFC^Cacna1h+^ neuronal projections to the AHNc during sociosexual preference test. (B) Schematic illustration of the pathway-specific chemogenetic inactivation strategy. (C) Overlay of AAV8-Ef1α-mCherry-IRES-Cre and AAV8-EF1α-mCherry expression in the mPFC of Cacna1h^fl/fl^ mice. Green intensity indicates the number of mice with viral expression in each area. (D) Cannula locations of chemogenetic inhibition experiments. (E-F) Coronal sections showing the expression of hM4Di-mCherry in the mPFC (E, scale bars, 100 µm [inset] and 500 µm) and cannula tracks above the mCherry+ axon terminals in the AHNc (F, scale bar, 200 µm). (G) Chemogenetic suppression of mPFC^Cacna1h+^ neurons-AHNc pathway decreased sociosexual interests specifically when female mice in estrus. N = 6 to 15 mice for each group. Two-way ANOVA, Bonferroni multiple comparisons test, p = 0.0002, * * * p < 0.001. (H) Chemogenetic inhibition of mPFC^Cacna1h+^ neurons-AHNc pathway in male mice enhanced their sociosexual preference for female mice. N = 9 mice for each group. Two-way ANOVA, Bonferroni multiple comparisons test, **** p < 0.0001.

**Figure S19.**
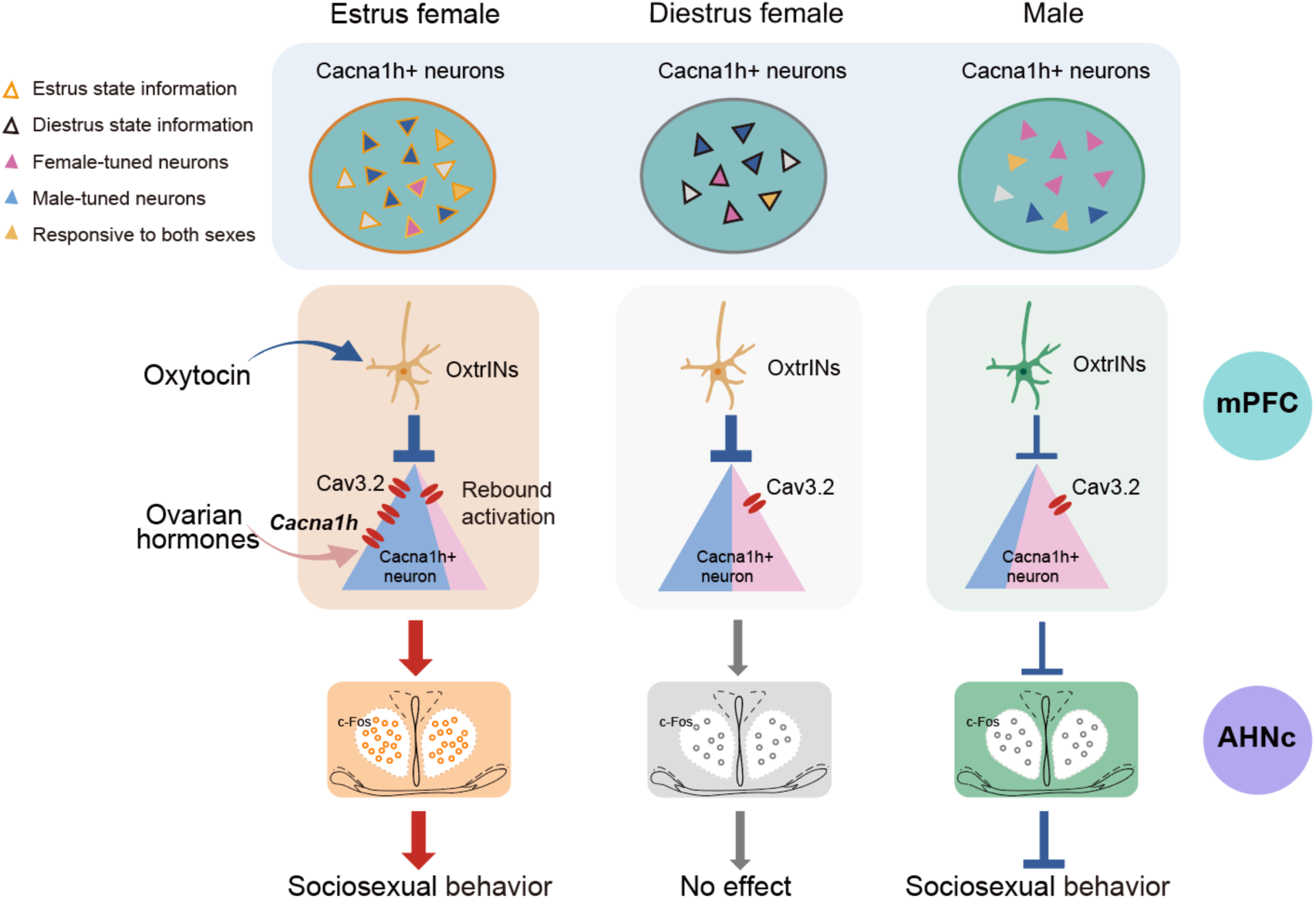
A model for the sexually dimorphic regulation of sociosexual behavior by mPFC^Cacna1h+^ neurons. In females, mPFC^Cacna1h+^ neurons exhibit mixed representation of internal estrous state and target-sex information. During estrus phase, these neurons are tuned to male cues and display more selective encoding of male-related information. Ovarian hormones upregulate *Cacna1h*-encoded T-type calcium channels via a genomic pathway, enabling enhanced rebound excitation following inhibition from oxytocin-responsive interneurons. This modulation drives estrus-specific activity changes in mPFC^Cacna1h+^ neurons, promoting sociosexual interests and sexual behavior through projections to the AHN. During diestrus, the low level of *Cacna1h* leads to insufficient T-type calcium channels, preventing the robust rebound activation observed in estrus females and ultimately diminishing selectivity for male cues compared to the heightened preference exhibited during estrus. Conversely, in males, mPFC^Cacna1h+^ neurons are biased towards the representation of female cues. However, due to low *Cacna1h* expression, these neurons exhibit a primarily inhibitory response to sociosexual stimuli, suppressing sociosexual interests and sexual behavior via the AHN pathway.

